# Role of lncRNAs related to NRs in the regulation of gene expression

**DOI:** 10.1101/2025.03.17.643776

**Authors:** Katerina Pierouli, Louis Papageorgiou, George P Chrousos, Elias Eliopoulos, Dimitrios Vlachakis

## Abstract

Long non-coding RNAs (lncRNAs) play a key role in regulating gene expression, influencing various cellular pathways by interacting with transcription factors, other RNA molecules like miRNAs, and DNA. In this study, we focused on the role of lncRNAs related to nuclear receptors (NRs), which are a family of transcription factors activated by ligands. NRs are involved in vital biological processes such as metabolism, immune response, reproduction, and development. We discovered six novel sequence motifs within lncRNAs that respond to multiple NRs suggesting they are not specific to a single receptor. One of the motifs, was complementary to miRNA hsa-miR-1908-3p, suggesting that lncRNAs containing this motif may function as miRNA sponges, regulating the expression of about 487 target mRNAs. The motifs were also found in key regulatory regions of the human genome, particularly on chromosome 19, including in Adeno-associated virus integration site 1 (AAVS1), a region with a high regulatory role in gene expression. Additionally, an evolutionary analysis was conducted revealing that these motifs are highly conserved across species, including *Mus musculus, Euglena gracilis,* and *Saccharomyces cerevisiae*, indicating their ancient origins. Based on these results, we propose that these motifs may represent ancestral binding sites for nuclear receptor precursors, and may act as backup mechanisms in modern organisms, ensuring the functional versatility and evolutionary conservation of NR-mediated gene regulation.

## 1. Introduction

By the sight of the central dogma of biology, RNA functions as an intermediate molecule helping transfer genetic information from the DNA level to the protein synthesis level. For many years, this process was considered the primary pathway for gene expression. However, over time, the functional importance of RNA transcripts and other non-coding regions, that are not directly encoded to proteins, has been recognized. Various of these non-coding RNAs (ncRNAs) are involved in several cellular functions, including the regulation of gene expression, alternative splicing, RNA processing, inhibition of translation, and mRNA degradation. In fact, RNA plays a crucial role in almost every level of genome regulation (1–3).

NcRNAs are typically divided into two main classes: structural and regulatory. Structural ncRNAs include ribosomal (rRNA), transfer (tRNA), and small nuclear RNAs (snRNA). On the other hand, regulatory ncRNAs are further classified into several subcategories such as microRNAs (miRNAs), Piwi-interacting RNAs (piRNAs), small interfering RNAs (siRNAs), and long non-coding RNAs (lncRNAs). In addition, a subclass of Promoter-Associated RNAs (PARs) that interacts with RNA promoters and is produced by genomic enhancer regions (enhancer RNAs, eRNAs) has recently been described(4).

LncRNAs are a highly diverse and heterogeneous class of ncRNAs, varying in their characteristics, locations in the genome, and functional roles. They are the most abundant class of non-coding transcripts, and they typically have low expression levels and are longer than 200 nucleotides (5). These RNAs can be categorized based on their location in the genome, into intronic, intergenic (lincRNAs), divergent, sense, and antisense lncRNAs (6, 7). Unlike small ncRNAs, lncRNAs are less conserved across species and their roles in the regulation of gene expression not still fully understood (8).

Even though the limited number of functional lncRNAs have been well characterized, they play essential roles in the regulation of gene expression at various stages. LncRNAs are involved in post-transcriptional regulation by controlling protein synthesis, RNA maturation, and RNA transport, as well as in gene silencing through modulation of chromatin structure. Because of the great variety of lncRNAs, categorizing them is challenging. However, several studies have classified them based on their molecular functions, as signaling molecules that relay temporal, spatial, and developmental information; as decoys that isolate various RNA molecules and proteins to inhibit their activity; as guides that direct epigenetic regulators and transcriptional factors to specific genomic loci; and as scaffolds that facilitate the formation of macromolecular complexes with a various functions (8). Importantly, individual lncRNAs may act through more than one mechanism, potentially leading to the combination of functions that create more complex functions in cellular procedures. Thus, understanding the shared features of these mechanisms is crucial for elucidating the functions of lncRNAs and their crucial roles in cellular regulation (8).

In addition to the aforementioned functions, several lncRNAs regulate gene expression by binding to miRNAs, acting as “miRNA sponges”. These lncRNAs contain multiple binding sites for one or more miRNAs and prevent their interaction with target mRNAs. As a result, the target mRNAs of the miRNAs are upregulated, increasing their expression (8, 9).

The direct and indirect interaction between lncRNAs and transcription factors are well-documented. For instance, the lncRNA Gas5, which is activated in the absence of growth factors, contains a hairpin structure sequence pattern that resembles the DNA-binding site of the glucocorticoid receptor (GR), a member of the nuclear receptors family (NRs). By acting as a decoy, Gas5 blocks the access of GR to DNA, thus inhibiting the transcription of GR metabolic target-genes (10). NRs are transcription factors that bind to DNA and regulate the expression of genes involved in processes such as homeostasis, stress response, and metabolism (11). Dysregulation of NRs has been reported in various diseases, including neurodegenerative diseases, cancer, metabolic syndromes, diabetes, obesity, and autism (12). For that reason, understanding the mechanisms of interaction between lncRNAs and NRs is critical for lighting up the complex molecular pathways involved and could lead to the development of more effective and specialized therapies for plenty of diseases.

To further explore the regulatory roles of lncRNAs in gene expression, we examined lncRNAs whose expression levels are altered following the activation of an NR. These lncRNAs were classified into two groups: those that are upregulated and those that are downregulated. Our aim was to explore whether any lncRNAs fail to exhibit a regulatory function. Considering the evolutionary conservation of lncRNAs, we hypothesized that the presence of sequence motifs within lncRNAs’ sequences could provide insight into their regulatory role. These motifs, if present with non-random frequency, may serve as signatures that contribute to the regulatory functions of the lncRNAs.

Using the MEME algorithm, we analyzed the lncRNA sequences from the aforementioned groups and identified six specific sequence motifs-three in each group. The potential roles of these motifs were investigated, providing insights into how lncRNAs containing these motifs may regulate a wide range of biological processes. . According to our results, one of the discovered motifs was found to be complementary to the miRNA hsa-miR-1908-3p, suggesting that lncRNAs containing this motif may act as sponges for this miRNA, regulating the expression of its target mRNAs. Additionally, all six motifs were detected in specific genomic regions of human genome, especially on chromosome 19, with a high percentage of occurrence. Among these regions on chromosome 19, the motifs were aligned with the Adeno-associated virus integration site 1 (AAVS1), a known hotspot for Adeno-associated virus (AAV) integration. Interestingly, many lncRNAs containing at least one of these motifs, participate in various biological pathways such as development, immune response, inflammation, cell cycle, regulation of gene expression, and homeostasis, highlighting the diverse roles of lncRNAs in cellular pathways. To explore the evolutionary significance of these motifs, we investigated their presence across various species, including *Mus musculus, Squamata (lizard), Danio rerio (zebrafish), Drosophila melanogaster, Caenorhabditis elegans, Escherichia coli (K12 MG1655), Saccharomyces cerevisiae, Bacillus subtilis (PY79) and Euglena gracilis,* which differ both evolutionarily and genetically. This cross-species analysis showed that these motifs are evolutionary conserved, suggesting their potential importance in fundamental biological processes. The conservation of these motifs across a wide range of species highlights their potential functional relevance, warranting further investigation into their roles in gene regulation and interaction with NRs.

## 2. Methods

In this study, we developed an in-house pipeline for the analysis of the collected dataset of lncRNAs with known expression levels, as documented in the literature. These lncRNAs are related to the activation and expression levels of NRs. This pipeline includes the following key steps:

1. *Literature review and data retrieval:* A comprehensive search in current literature was conducted to identify lncRNAs whose expression levels are known to be associated with the activation of NRs. In this stage, we also retrieved the corresponding RNA sequences for further analysis.
2. *Classification of lncRNAs*: The identified lncRNAs were classified into two groups. The first group includes lncRNAs whose expression is upregulated following the activation of an NR, while the second group includes lncRNAs whose expression is downregulated in response to the activation of an NR. Afterwards, conserved sequence motifs were identifiedwithin these two lncRNA sub-groups.
3. *Statistical evaluation of motif significance*: The identified motifs were statistically evaluated to determine their significance. This involved estimating the probability of these motifs occurring randomly within the entire human DNA.
4. *Exploration of motif functionality*: In this step, we explored the potential functions of the identified motifs, focusing on their role in gene regulation. This included investigating their potential to function as binding sites for miRNAs , examining their presence within genome regulatory regions and exploring their conservation through evolution. These analyseshelped us understand the broader roles of these lncRNAs in cellular processes.

A detailed description of the in-house pipeline is provided in the following subsections.

### 2.1 Data collection and filtering and annotation

The IDs of lncRNAs related to the NRs were collected from the literature according to the study of Foulds et al. (13). The IDs of lncRNAs had the following formats:

1. Name of lncRNA (e.g. MALAT-1)
2. ID such as AC145110.1
3. ID such as NR_026765
4. ID such as RP1-148H17.1

The first three ID formats correspond to different types of IDs detected at GenBank, a comprehensive genetic sequence database available at NCBI (ncbi.nlm.nih.gov). The fourth ID format derived from LNCipedia database (14), a novel database for human lncRNA transcripts and genes.

Based on these ID formats, the dataset of lncRNAs was classified into four subcategories. To retrieve the corresponding sequences for each subcategory, an in-house algorithm was developed using the MATLAB Bioinformatics Toolbox. The details of the four algorithms used for sequence retrieval are shown in **Data S1**.

Following the creation of the four individual algorithms for each type of lncRNA ID, we developed a unified algorithm designed to retrieve the sequences of the entire dataset, containing all ID formats simultaneously. The aggregate algorithm used for retrieving all lncRNA sequences is shown in **Data S2**.

### 2.2 Dataset classification and Motif exploration

In order to further classify the lncRNAs within the two sub-datasets, we investigated the presence of joint motifs across the lncRNA sequences in each subgroup. Following the collection, filtering, and annotation of the data, the MEME (Multiple EM for Motif Elicitation) software, Version 4.10.0 (15), was employed to identify joint motifs within the lncRNA sequences of each subgroup. MEME is a part of the ΜEME Suite (https://meme-suite.org/meme/), whichis a comprehensive set of tools for motif-based sequence analysis. This tool is designed to detect motifs or sequence patterns that occur repeatedly in a group of related nucleotide or protein sequences, finding common biological functions and structural features across different sequences. The term “pattern” or “motif” refers to a short, sequence that repeates within a group of similar sequences. The motifs have no gaps, while the MEME algorithm calculates how often each motif appears across sequences, allowing the detection of multiple motifs (15).

For the purposes of this study, we ran the MEME algorithm on a local server using the Linux operating system.

### 2.3 Statistical evaluation of the randomness of the occurrence of the discovered motifs

To calculate the probability of finding specific nucleotide bases of each motif at a random position within the 3,2 billion bases of the human DNA, we assumed that the occurrence of a nucleotide at one position in the motif’s sequence is independent of the occurrence of nucleotides at other positions. This assumption represents the simplest hypothesis, involving the fewest parameters. Thus, we applied the rule of probabilities for independent events, which defines the probability of multiple events occurring independently. Specifically, two events are considered independent if the occurrence or non-occurrence of one does not influence the probability of the other event’s occurrence or non-occurrence (16, 17).

To calculate the probability of both events A and B occurring together, we used the formula for independent events:

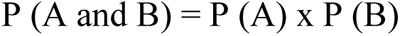

This approach was applied to assess the randomness of each motif existence within human DNA.

### 2.4 Exploration of the potential role of the identified motifs

A total of six motifs were discovered, three in each subgroup of lncRNAs. To explore the potential role of the collected lncRNAs in the regulation of gene expression, we focused on the lncRNAs that contained at least one of the discovered motifs. Since motifs are usually conserved through evolution and contribute crucial functions to the regions they occupy, we proceeded to explore the potential roles of these motifs in genome regulation, which could reflect the functions of the lncRNAs that contain them. For that purpose, we pursued two different approaches. The first included querying the miRBase database (https://www.mirbase.org/) to identify mature miRNAs with complementary sequences to these motifs, suggesting the possibility of miRNA-lncRNA interaction. The second approach included the exploration of the presence of these motif sequences across whole human DNA in order to determine their existence in other genomic regions, assess whether they were associated with regulatory regions, and explore potential roles in genomic regulation. In addition, we explored the evolutionary conservation of the motifs to determine the maintenance of their functions through evolution.

#### 2.4.1 Exploration of the miRNA-target genes’ interactions

The main purpose of this approach was to identify the potential role of lncRNAs containing the identified motifs in functioning as “sponges” for miRNAs, where the motifs serve as interaction/binding regions between lncRNAs and miRNAs. For that purpose, we used the blastn algorithm from miRbase, with the complement sequence of each motif as input. This allowed this algorithm to search for similarities between the motif sequences and known miRNA sequences in the miRBase. Afterward, we further investigated the predicted target genes of the identified miRNA using the bioinformatic tool MirTarget from the miRDB database (16), and the TargetScanS database (Release 8.0) (17–19).

Finally, we examined the location of the binding sites in the target mRNAs by using the multiple alignment algorithm ClustalW, using as input the mRNA sequence of the targets and the complementary sequence of miR-1908-3p.

#### 2.4.2 Exploration of motif sequences in whole human DNA

To investigate if these motif sequences also appear in other regions of the genome, we used the online tool FIMO (Find Individual Motif Occurrences) from the MEME Suite. This tool is an algorithm designed to identify specific matches of a given motif within a set of sequences, specialized for analyzing shorter sequences like motifs. Similar to a BLAST algorithm, FIMO searches for matches to each motif. When employed locally, motifs must be in MEME Motif Format; however, the online version of FIMO supports additional motif formats.

Afterwards, we developed an algorithm using the MATLAB Bioinformatics Toolbox to create a chromosome plot for each identified motif. This plot displayed the locations of the motif sequences by marking them with a red dot next to the corresponding regions on the chromosome where they were detected. The input for this algorithm was the output from the FIMO tool in .tsv format. The specific algorithm we developed is shown in **Data S3**.

We also used the FIMO tool to examine the presence of the identified sequence motifs in the genomes of various organisms, including *Mus musculus, Squamata (lizard), Danio rerio (zebrafish), Drosophila melanogaster, Caenorhabditis elegans, Escherichia coli (K12 MG1655), Saccharomyces cerevisiae,* and *Bacillus subtilis (PY79)*, to assess their evolutionary conservation. Additionally, we investigated the presence of these motifs in the genome of *Euglena gracilis*, a protist believed to possess an ancient form of NRs. Due to the unavailability of *Euglena’s* genome in the FIMO tool’s database, the “Matcher” tool available on the Galaxy platform (usegalaxy.org) was employed to check for potential sequence matches (20).

Galaxy is an open-source, web-based platform that aids computational biology research by providing a comprehensive suite of tools for data analysis. The “Matcher” tool within this platform compares two different sequences to identify regions of similarity, supporting both local and global alignments. This tool enables the identification of conserved genomic regions across species and helps in detecting functionally important motifs that have been preserved through evolution. This tool is widely used in genomics and bioinformatics for tasks such as gene annotation, mutation analysis, and the discovery of conserved regulatory regions (21).

## 3. Results and Discussion

### 3.1 Data collection, filtering, and annotation

The initial dataset, collected from the study by Foulds et al. (13), contained 1823 lncRNA IDs, with sequences for 1788 lncRNAs successfully identified and retrieved. After the removal of the duplicate sequences, the final dataset consisted of 1753 lncRNAs. These were then classified into two subgroups: the first subgroup contained 778 upregulated lncRNAs and the second subgroup contained 975 downregulated lncRNAs (**Table S1**).

### 3.2 Dataset classification and Motif identification

The MEME algorithm was employed to identify potential motifs within the lncRNA sequences of each subgroup. The results of this analysis led to the identification of six motifs-three motifs in the upregulated lncRNA group and three in the downregulated lncRNA group.

In the upregulated lncRNAs group following NR activation, the first identified motif is GATCCGCCCGCCTCGGCCTCCCAAAGTGCTGGGATTACAGG, found in 140 lncRNAs from the total 778 (**Figure S1**). The second motif discovered in this group is TGGCCAGGCTGGTCTCGAACTCCTGACCT, present in 127 lncRNAs (**Figure S2**), while the third motif is TAATCCCAGCACTTTGGGAGGCCGAGG, which occurs in 115 lncRNAs (**Figure S3**).

For the downregulated lncRNAs group after NR activation, the first motif discovered is GTGGCTCACGCCTGTAATCCCAGCACTTTGGGAGGCCGAGG, found in 145 lncRNA out of the total of 975 (**Figure S4**). The second motif discovered is AGGTCAGGAGTTCGAGACCAGCCTGGCCAACATGGTGAAAC, which is present in 128 lncRNAs (**Figure S5**), and the third motif is CCTCAGCCTCCCGAGTAGCTGGGACTACAGGCGC, found in 129 lncRNAs from this group (**Figure S6**).

### 3.3 Statistical evaluation

To calculate the probability of each motif appearing randomly at a position within the 3,2 billion base pairs of human DNA, the following steps were taken:

The first motif from the upregulated lncRNAs subgroup consists of 41 nucleotide bases. Since each base (A, C, G, or T) has a ¼ probability of being in a specific position in the sequence, the probability of finding the full sequence of these 41 nucleotide bases is:

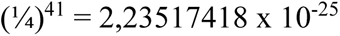

Taking into consideration the total number of humans DNA base pairs (3,2 billion or 3,2 x 10^9^), the probability of this sequence occurring randomly at any position in the genome is:

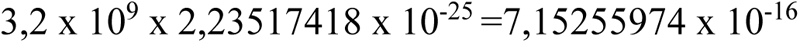

Accordingly, for the second motif in the upregulated lncRNAs subgroup, consisting of 29 nucleotides, the probability calculation is as follows:

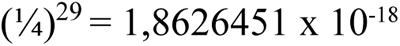

The probability of finding this specific sequence randomly in human genome is:

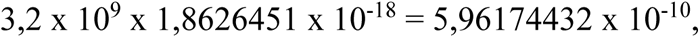

For the third motif in the upregulated lncRNAs subgroup, which consists of 27 nucleotides, the results are as follows:

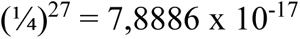

Thus, the probability of its random occurrence is:

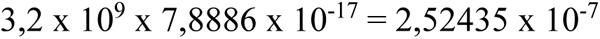

In the downregulated lncRNAs subgroup, both first and second motifs consist of 41 nucleotides, so, the same probability calculations apply:

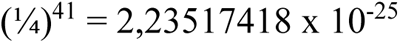

Thus, the probability of their random occurrence is:

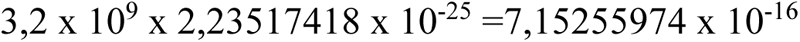

For the third motif in the downregulated lncRNAs subgroup, consists of 34 nucleotides:

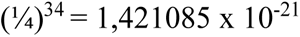

Thus, the probability of its random occurrence is:

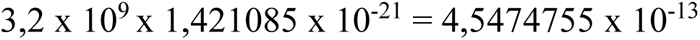

The calculated probabilities are extraordinarily low compared to common biological standards and are consistent with the idea that these motifs are not randomly occurring in the genome (22).

The above-calculated probabilities suggest that the motifs discovered are non-random occurrences in the human genome, even when assuming independent events. However, the nucleotide sequence of DNA is linked to biological functions. The order of the bases is strictly determined by the way nucleotides bind each other in the double helix structure. The complementary pairing of nucleotides restricts the possible sequences that DNA can form, meaning that the probability of these motifs occurring randomly in the genome would likely be even lower than the values calculated above.

### 3.4 Exploration of the potential role of the identified motifs

#### 3.4.1 Exploration of the miRNA-target genes’ interactions

As previously mentioned, lncRNA can function as miRNA sponges, regulating the expression of mRNA targets. Considering this regulatory role, we searched for potential miRNAs complementary to the six identified motifs in miRBase. Our search revealed that the motif GATCCGCCCGCCTCGGCCTCCCAAAGTGCTGGGATTACAGG is complementary to hsa-miR-1908-3p.

ΜiRNAs typically bind to complementary sequences in mRNAs, mainly targeting the 3’ untranslated regions (UTRs) (23), while there is evidence that they can also target the 5’ UTRs and coding regions, thus inhibiting mRNA expression (24). To predict potential targets of miRNAs, several computational approaches have been developed, such as miRanda (25), mirSVR (26), PicTar (27), TargetScan (28), TargetScanS (29), RNA22 (30), PITA (31), RNAhybird (32) and DIANA-microT (33). In addition, there are miRNA target prediction databases such as TarBase (34), PicTar (27), and TargetScan (28).

Thus, to identify mRNAs targeted by hsa-miR-1908-3p, whose expression may be affected by the interaction between lncRNAs and this miRNA, we used the miRDB database. This database, through its MirTarget bioinformatics tool, predicts the mRNA based on high-throughput sequencing experiments. By this analysis, 11 targets were identified, including *METTL21A, CHFR, MTA1, CAVIN4, IGFBP3, GRB2, DYNLL2, H2AFX, TNRC6C, C1orf229*, and *ELOVL2* (**Table I**). We also searched the TargetScanS (Release 8.0) database (17–19, 35), and found 487 predicted targets (**Table S2**), including the aforementioned 11 targets.

**Table I.** The target genes of mir-1908-3p were predicted by the MirTarget tool of miRDB.

| Gene Symbol | Gene Description | miRNA Name | Target Score |
| --- | --- | --- | --- |
| **METTL21A** | methyltransferase like 21A | hsa-miR-1908-3p | 80 |
| **CHFR** | checkpoint with forkhead and ring finger domains | hsa-miR-1908-3p | 62 |
| **MTA1** | metastasis associated 1 | hsa-miR-1908-3p | 62 |
| **CAVIN4** | caveolae associated protein 4 | hsa-miR-1908-3p | 61 |
| **IGFBP3** | insulin like growth factor binding protein 3 | hsa-miR-1908-3p | 61 |
| **GRB2** | growth factor receptor bound protein 2 | hsa-miR-1908-3p | 56 |
| **DYNLL2** | dynein light chain LC8-type 2 | hsa-miR-1908-3p | 55 |
| **H2AFX** | H2A histone family member X | hsa-miR-1908-3p | 53 |
| **TNRC6C** | trinucleotide repeat containing 6C | hsa-miR-1908-3p | 53 |
| **C1orf229** | chromosome 1 open reading frame 229 | hsa-miR-1908-3p | 51 |
| **ELOVL2** | ELOVL fatty acid elongase 2 | hsa-miR-1908-3p | 50 |

Mir-1908 is encoded by the first intron of the *FADS1* gene and is highly expressed in mature human adipocytes, where it is likely involved in regulating the preadipocyte differentiation (36). The promoter region of miR-1908 contains two binding sites of NF-κB, suggesting its transcription is regulated by this transcription factor (37). MiR-1908 was first discovered in human embryonic stem cells in 2008 (38), and its functional role in melanoma metastasis and angiogenesis was explored in 2012. More specifically, miR-1908, in conjunction with miR-199a-3p and miR-199a-5p, inhibits APOE and DNAJ Heat Shock Protein Family 23 Member A4 (DNAJA4), which reduces the interaction of Apo-E with LRP-1 and LRP-8. This interaction promotes melanoma cell invasion and endothelial cell recruitment, resulting in poor prognosis (39). Moreover, miR-1908 dysregulation has been associated with various types of cancer, where its dysregulation tends to enhance cell proliferation, invasion, migration, and angiogenesis, reducing survival rate (40, 41). Finally, free fatty acids and adipokines, like resistin and leptin, are believed to downregulate miR-1908 expression, suggesting a role of this miRNA in regulating obesity-related insulin resistance (42).

#### 3.4.2 Exploration of motif sequences in whole human DNA

For the analysis of motif sequences across the whole human genome using the FIMO tool, the following results were obtained for each of the six motifs. In the upregulated lncRNA subgroup, the first motif appeared 53524 times, the second motif was detected 51673 times and the third motif was found 53936 times. In the downregulated lncRNA subgroup, the first motif (fourth in total) was found 63126 times, the second motif (fifth in total) was detected 58442 times and the third motif (sixth in total) was found 57943 times. Using these results, we create karyotypes for each motif, and calculated the percentage of the occurrences of each motif across all chromosomes. The karyotypes are presented in **Figures S7-12**.

Afterward, we focused on the 1000 highest-scoring loci occurred by the FIMO tool for each motif across the human genome. Using these 1000 loci, karyotypes for each motif were also created, and the percentage of occurrences of each motif was also calculated for each chromosome (**Figures 1-6**).

**Figure 1.**
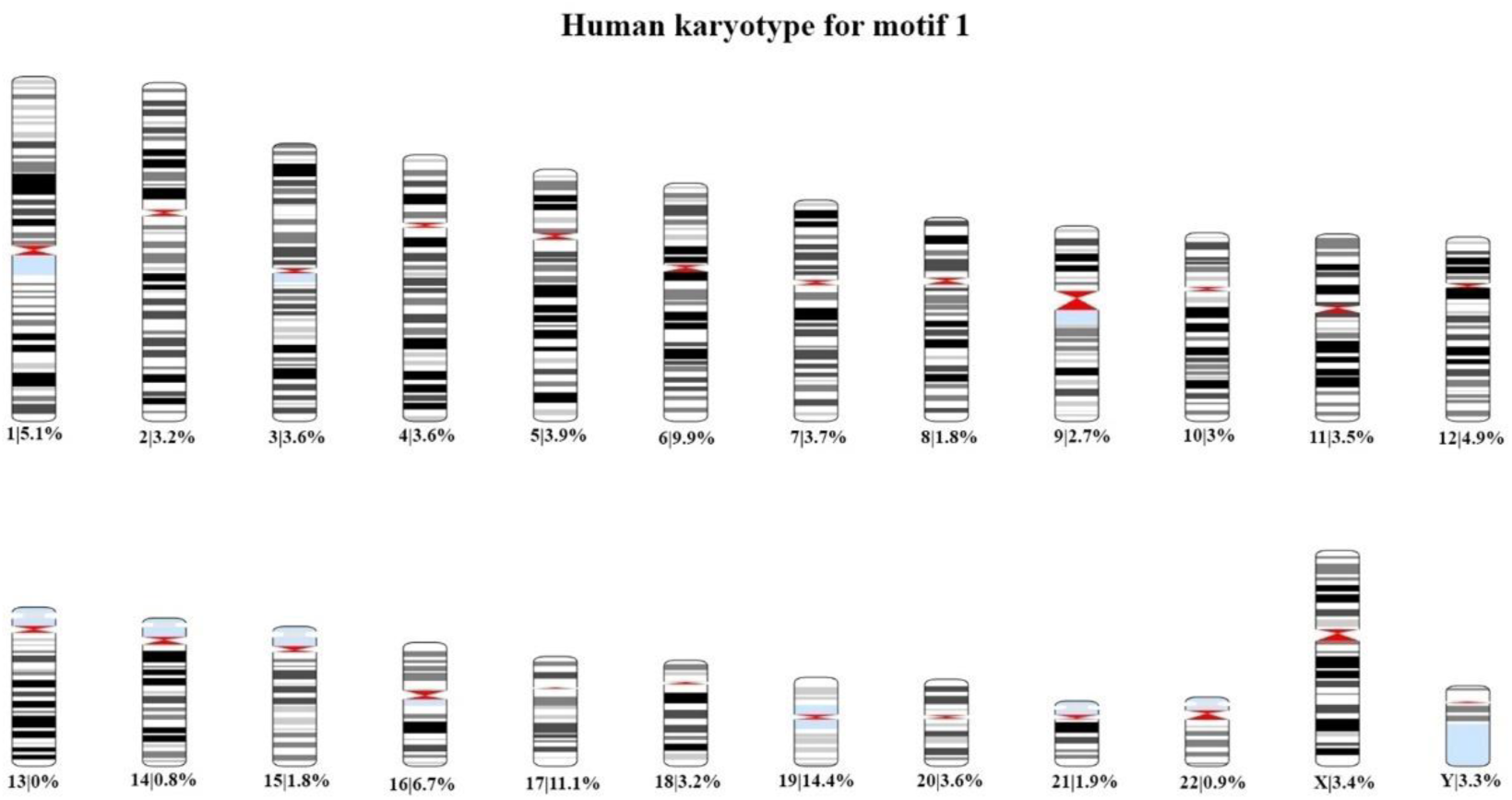
Human karyotype for the 1000 higher-scored loci that presents the percentage of the occurrence of motif 1 in each chromosome.

**Figure 2.**
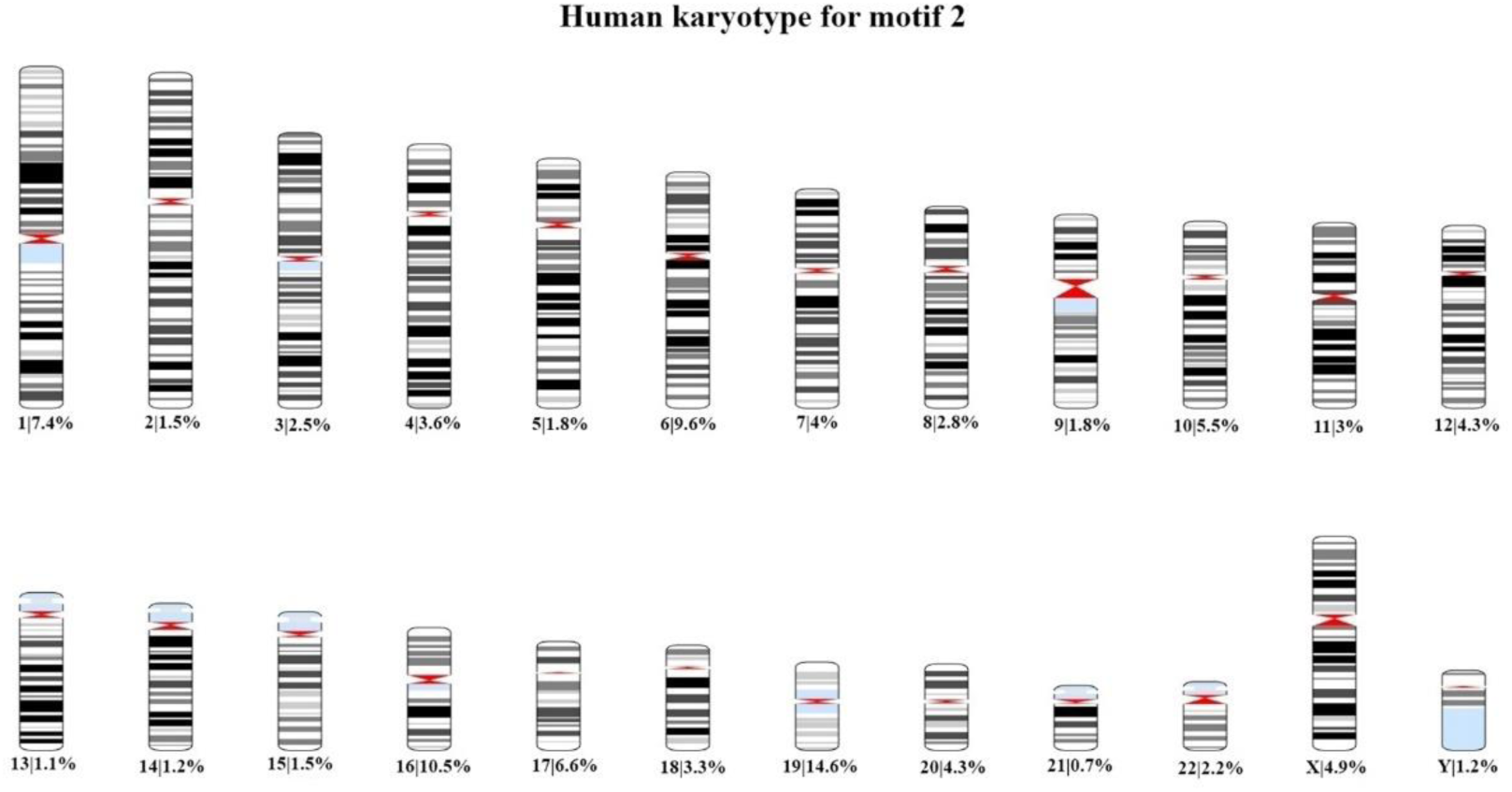
Human karyotype for the 1000 higher-scored loci that presents the percentage of the occurrence of motif 2 in each chromosome.

**Figure 3.**
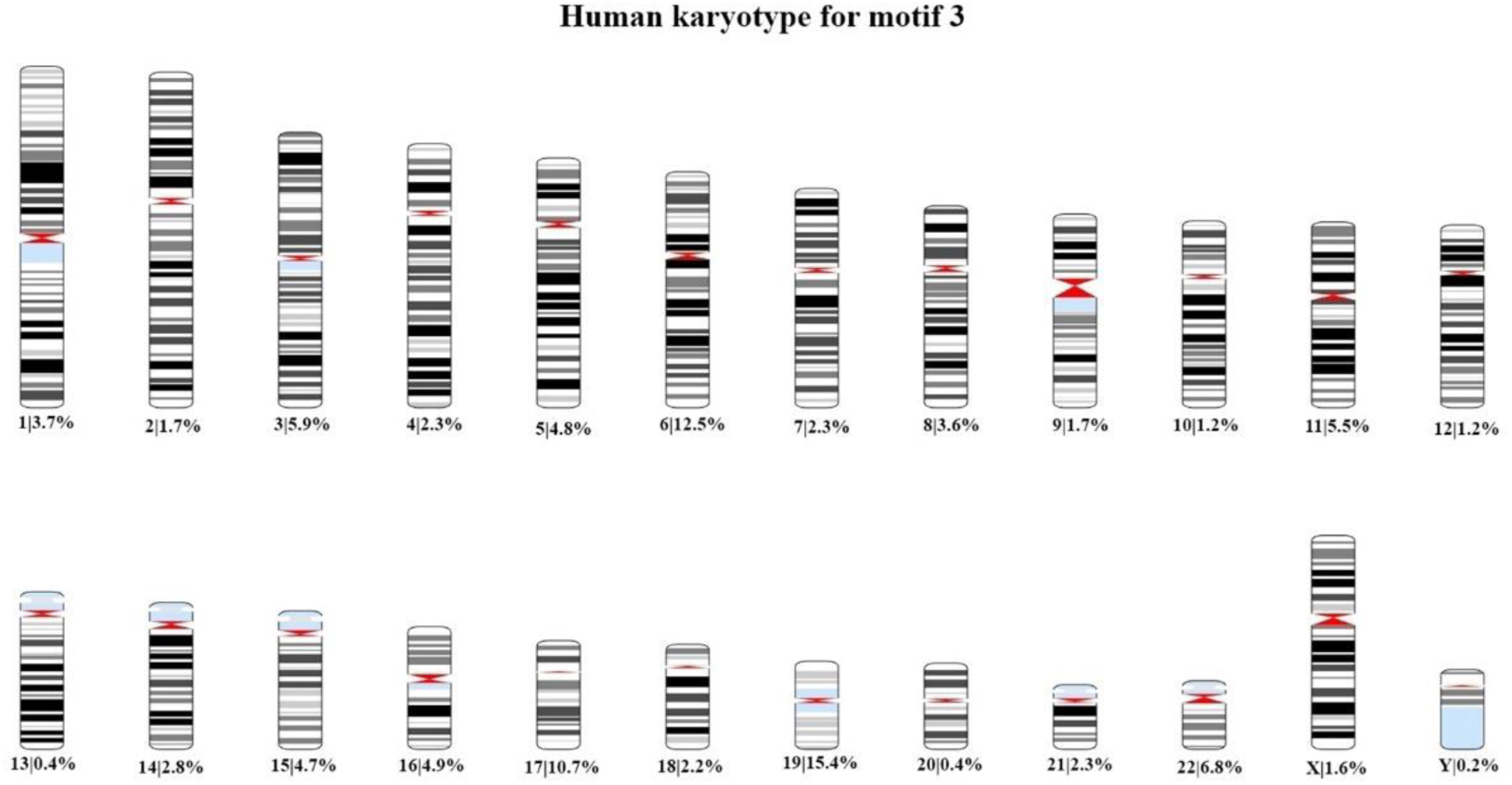
Human karyotype for the 1000 higher-scored loci that presents the percentage of the occurrence of motif 3 in each chromosome.

**Figure 4.**
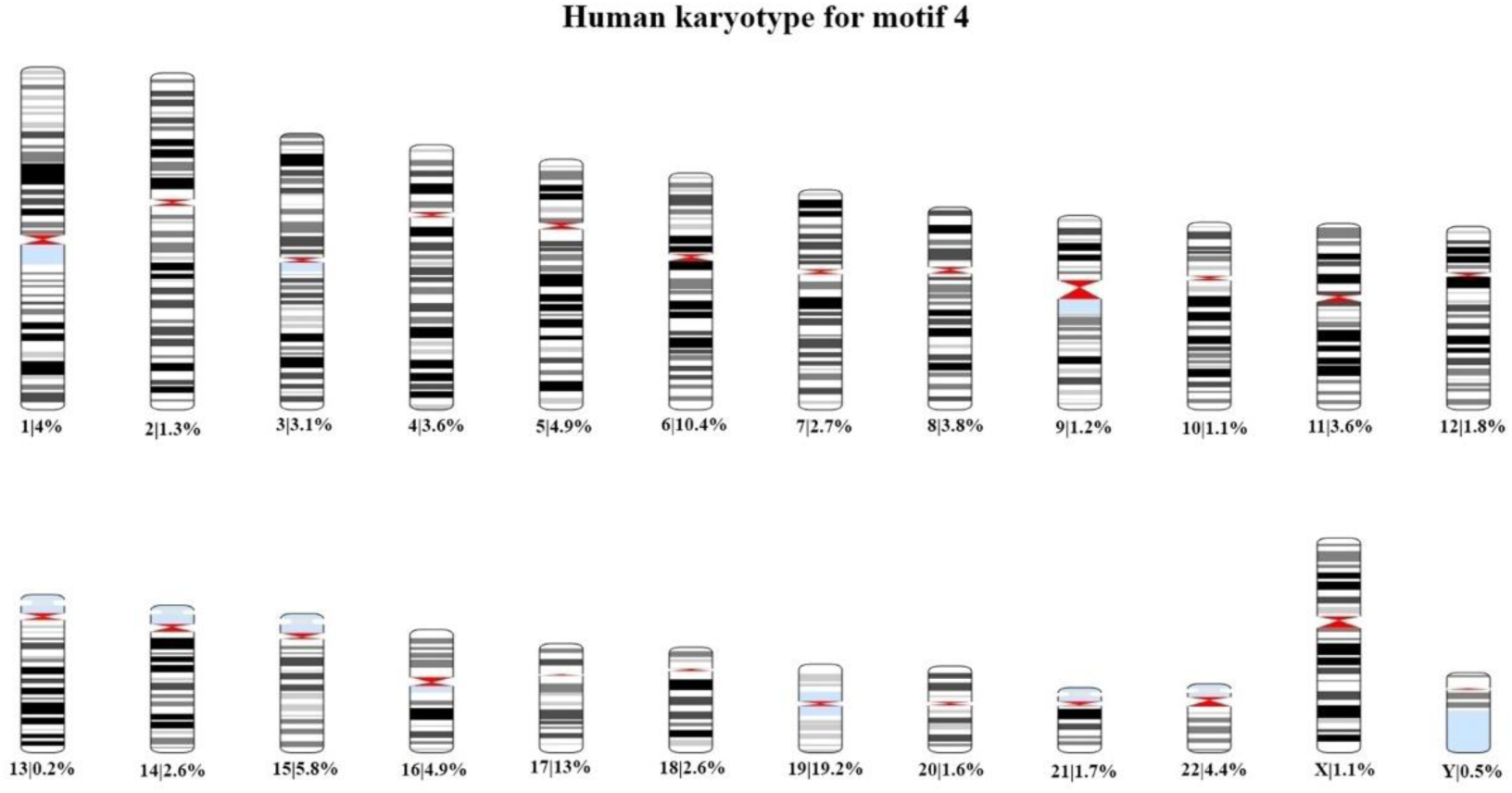
Human karyotype for the 1000 higher-scored loci that presents the percentage of the occurrence of motif 4 in each chromosome.

**Figure 5.**
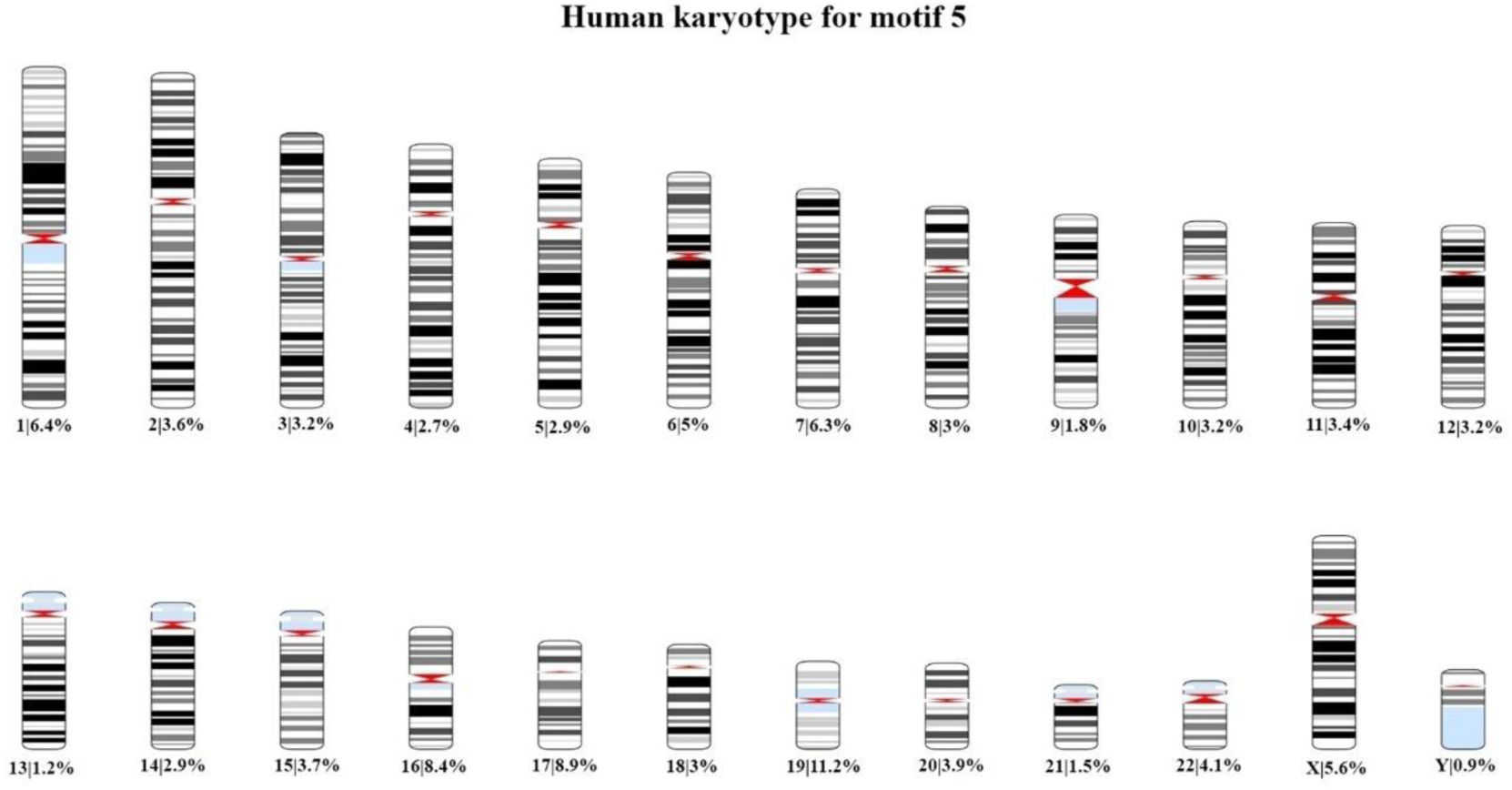
Human karyotype for the 1000 higher-scored loci that presents the percentage of the occurrence of motif 5 in each chromosome.

**Figure 6.**
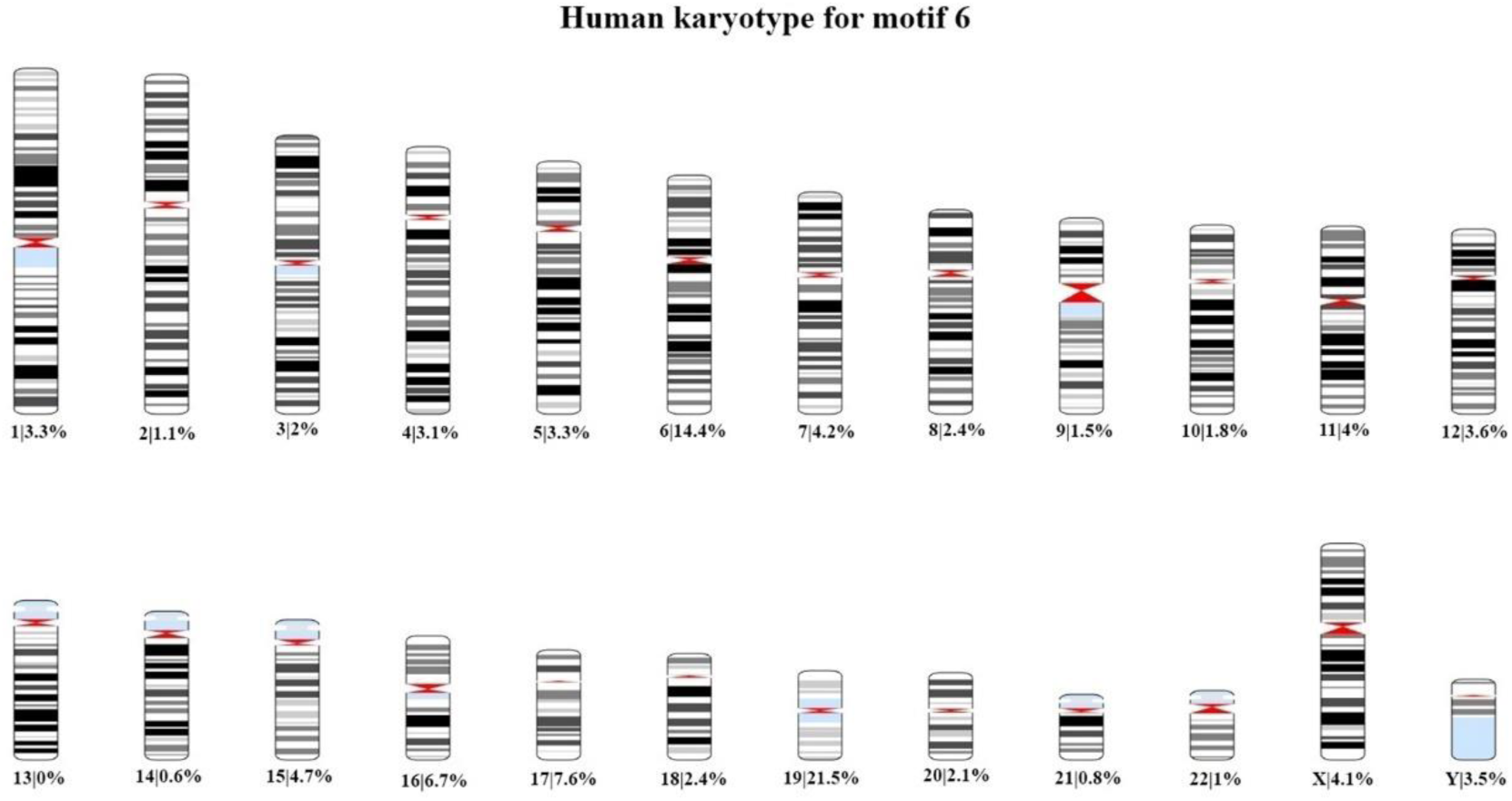
Human karyotype for the 1000 higher-scored loci that presents the percentage of the occurrence of motif 6 in each chromosome.

In addition, the chromosome plots for each motif were created, where the 1000 loci are represented as red dots beside the corresponding region. These chromosome plots are presented in **Figures S13-18** .

Based on the these karyotypes and chromosome plots, we observed that chromosome 19 exhibits a fairly high percentage of motif occurrence relative to its size. In fact, for all six motifs, the percentage of occurrence in chromosome 19 is higher than that of longer chromosomes.

According to the literature, human chromosome 19 is recognized for its unusual nature, which was noted even before the initial publication of its DNA sequence (43). One of its unusual features is the considerably high gene density, which exceeds the genome-wide average by more than twofold and includes 20 large, tandemly clustered gene families (43). In addition, chromosome 19 contains a considerable number of segmental duplications, with 6.2% of its sequence lying within intrachromosomal segmental duplications (43). The sequence divergence within these duplications implies that these events occurred between 30 and 40 million years ago (MYA) (44). These duplications likely played a pivotal role in the evolution of phenotypic characteristics that occurred by genes on chromosome 19 across primates, including humans. Moreover, chromosome 19 presents an unusually high repeat content, comprising 55%, with Alu repeats comprise 26% of the chromosome’s sequence (44). Another remarkable feature is the GC content (48%) of chromosome 19, the highest of any human chromosome, compared to the GC content of the whole human genome of 41%. This elevated GC content enhances the potential for wide gene regulation via DNA methylation, particularly at CpG islands, as well as in CpG regions located within promoters and enhancers (45).

According to the chromosome plots, the six identified motifs have been detected in various regions of chromosome 19. Between those regions, all six motifs aligned with a specific locus, the Adeno-associated virus integration site 1 (AAVS1), located on chromosome 19q13.42. This site was first mapped in 1991 Kotin et al. (46) and Samulski et al. (47), who were investigating the integration mechanism of the adeno-associated virus (AAV) into the host genome. According to these studies, the AAV preferentially intergrades into the AAVS1 region of human chromosome 19q13.42, a unique phenomenon among eukaryotic DNA viruses (48).

Several features make the AAVS1 region particularly interesting. Notably, the locus for myotonic dystrophy (49), as well as a common breakpoint related to chronic B-cell leukemia (BCL-3) (47, 48), are located at this region. In addition, AAVS1 undergoes frequent sister chromatid exchanges (50). The alignments of motifs with the AAVS1 region were developed using the Clustal Omega program (51) and visualized with Jalview version 2.11.2.6 program (52) (**Figures S19-24**).

The AAVS1 region is located within the first intron of the phosphatase 1 regulatory subunit 12C (*PPP1R12C*) gene, which encodes a protein with a poorly defined function. This region exhibits several unique features that contribute to its unique nature. First, it includes a tetrad GACT repeat, which functions as an AAV Rep-binding element, also detected in the AAV2 inverted terminal repeats (ITR), facilitating the exclusive integration of the AAV genome in the presence of AAV replication protein. Second, chromosome 19 is GC-rich, with a higher number of CpG islands relative to GpC islands, a characteristic linked to regions of transcriptional activity in vertebrates, which may be relevant to gene expression (53–55). Third, several putative transcription factor-binding sites are present in AAVS1 including CREB, AP-1, AP-2, and Sp1 (48, 56). Fourth, topoisomerase I is predicted to cleave at least two sites within AAVS1 (48). Fifth, Lamartina, et al. (56) have proved an open chromatin conformation of AAVS1, accompanied by promoter activity on chromosome 19 in cultured cells. All these features position AAVS1 as an approachable region, subject to regulation by both host and viral regulatory elements.

The potential of the AAVS1 site as a target for gene therapy applications has been investigated and well-characterized. This site is known for its ability to integrate exogenous DNA with limited disruption to endogenous genes, providing a preferred site for stable transgene expression (57). An important feature that facilitates the usage of the AAVS1 site in gene therapy is its open chromatin structure, which is accompanied by endogenous insulator elements that protect the integrated DNA sequences from trans-activation or repression (58).

The insertion of exogenous genes into the human genome is conducted mainly using lentiviral and gamma-retroviral vectors. However, these vectors tend to integrate randomly within the genome (59–61), which can lead to unpredictable interactions between the transgene and the host genome. Such random integration may result in attenuation or complete silencing of the transgene (62–64) or, more critically, dysregulation of the host gene expression (64, 65), which poses significant risks in gene therapy application. One promising strategy to mitigate these risks involves the use of gene-editing tools that enable site-specific insertion of transgenes into genomic safe harbor (GSH) (64, 66, 67). GSHs are genomic locus that support a stable environment for transgene expression without interfering the integrity of endogenous genes (64, 68, 69). The AAVS1 site is considered a GSH that provides a stable transgene expression (70) due to the presence of flanking insulator regions (58). According to the study of Lombardo et al.(66), the GSH AAVS1 can support transgene expression across different human cell types, facilitated by the active and open chromatin structure of the PPP1R12C gene located in this region (66).

The regulatory role of lncRNAs is well-characterized. Numerous studies have detected that lncRNAs function as crucial regulators of gene expression, participating in various pathways by acting as decoys, guides, and scaffolds for molecular interactions. In this study, we hypothesized that the regulatory roles of lncRNAs are indisputable, and to investigate this hypothesis, we developed an in-house pipeline to analyze a collection of lncRNAs. This pipeline enabled the identification of conserved motifs within their sequences that may contribute to their regulatory functions. Our analysis that one of the motifs may function as a sponge for a specific miRNA, thereby suppressing the miRNA’s function and leading to the upregulation of its mRNA targets. The function of miRNAs in the suppression of the translation of mRNA targets is well known, with implications for numerous pathological conditions, such as neurodegeneration, cancer, and cardiovascular diseases. Consequently, the interaction between lncRNAs and miRNAs could have significant consequences, either promoting disease prevention and treatment or exacerbating disease progression and severity.

Furthermore, all six motifs identified in this study were detected across numerous regions of the human genome, with chromosome 19 presenting a significantly high frequency of motif occurrence relative to other chromosomes. Focusing on chromosome 19, we found that these motifs were aligned with the AAVS1 regulatory region, a genomic locus known for its unique features, including the specific integration of the AAV, the presence of transcription factor binding sites, an AAV Rep-binding element, high GC-content, open chromatin, and two cleavage sites for topoisomerase I. It is noteworthy that the identified motifs demonstrate a pronounced GC content, a feature commonly associated with key regulatory regions, including transcription factor binding sites, polymerase binding sites, and loci involved in chromatin modification and DNA methylation. GC-rich sequences are often enriched at genomic sites that regulate transcription and chromatin remodeling due to their structural properties, such as enhanced stability and the presence of CpG sites that are key for epigenetic regulation (71–74). The existence of these motifs in regions integral to the regulation of the human genome, combined with their GC-rich nature, suggests their role in regulating gene expression through DNA methylation processes. Consequently, these motifs and the lncRNAs containing them, likely function as pivotal regulatory elements in gene expression.

In addition to the potential functions of lncRNAs that contain the identified motifs-based on the presence of these motif sequences in well-characterized regulatory regions of DNA and their interactions with other molecules, including miRNAs- we further investigated the documented roles of the lncRNAs containing at least one of these motifs, in the literature (75–117) and relevant databases such as NCBI (ncbi.nlm.nih.gov) and GeneCards (118). Despite the incomplete annotation and characterization of these lncRNAs, most of the examined lncRNAs are located either within intergenic space, introns of coding genes, or are antisense to coding genes. These lncRNAs are believed to influence the functions of their associated genes, mainly by suppressing gene functions. The lncRNAs examined are involved in various biological processes, such as development, cellular proliferation, immune responses, inflammation, transmembrane transport, gene expression regulation, chromatin, and chromosome structure modulation, differentiation, cell cycle control, spermatogenesis, metabolic regulation, and several functions related to nervous and cardiovascular system. The results from our research, as presented in the diagrams in **Figures S25-30**, illustrate the involvement of lncRNAs with each specific motif in these diverse biological pathways.

The emergence of nuclear receptors dates back over 1.5 billion years, originating in early eukaryotic organisms as ligand-dependent transcription factors. These receptors were responsive to environmental cues such as hormones, light, and metabolic shifts. The earliest nuclear receptors likely played a critical role in regulating gene expression, helping primitive eukaryotes adapt to dynamic external conditions, thus aiding their survival and proliferation. Remarkably, unicellular organisms like *Euglena gracilis*, a photosynthetic protist, contain an ancestral version of the nuclear receptor, providing evidence that these mechanisms were present well before the evolution of multicellular organisms. This discovery enhances our understanding of the origins of nuclear receptors and their evolutionary significance (119).

As evolution progressed, nuclear receptors diversified into a complex protein family with distinct functions. Different subfamilies evolved to regulate a wide variety of biological processes across animals, plants, fungi, and even bacteria. In modern organisms, nuclear receptors govern critical processes such as metabolism, immune response, reproduction, and development. The diversity of these receptors is reflected in the large number of subfamilies, including steroid hormone receptors, thyroid hormone receptors, peroxisome proliferator-activated receptors (PPARs), and orphan nuclear receptors. Despite their varied roles, these receptors retain a conserved ligand-binding domain that allows them to interact with specific signaling molecules to regulate gene expression (119).

In our study, we discovered six novel sequence motifs within the lncRNA sequences under investigation. These lncRNAs showed altered expression levels upon activation of nuclear receptors, either increasing or decreasing in abundance. Notably, several of these lncRNAs, which contain at least one of the identified motifs, are regulated by multiple NRs. This suggests that these motifs may be associated with NR function but do not show specificity to any single receptor, underscoring the complexity of the regulatory network between NRs and ncRNAs. This finding highlights the importance of further research to explore their broader biological implications.

To investigate the evolutionary conservation of these motifs, we applied the FIMO tool to examine their presence in various organisms. We analyzed species such as *Mus musculus, Squamata (lizard), Danio rerio (zebrafish), Drosophila melanogaster, Caenorhabditis elegans, Escherichia coli (K12 MG1655), Saccharomyces cerevisiae, and Bacillus subtilis (PY79)*. Additionally, we explored the presence of these motifs in the genome of *Euglena gracilis* using the “Matcher” tool on the Galaxy platform. A phylogenetic tree was subsequently constructed using the iTOL tool (Interactive Tree Of Life), which enabled us to visualize and annotate the evolutionary relationships among the organisms and the conservation of the motifs across species (120). The resulting phylogenetic tree (**Figure 7**) clearly illustrates the evolutionary conservation of these motifs, which were identified repeatedly in the studied organisms (**Table II**).

**Figure 7.**
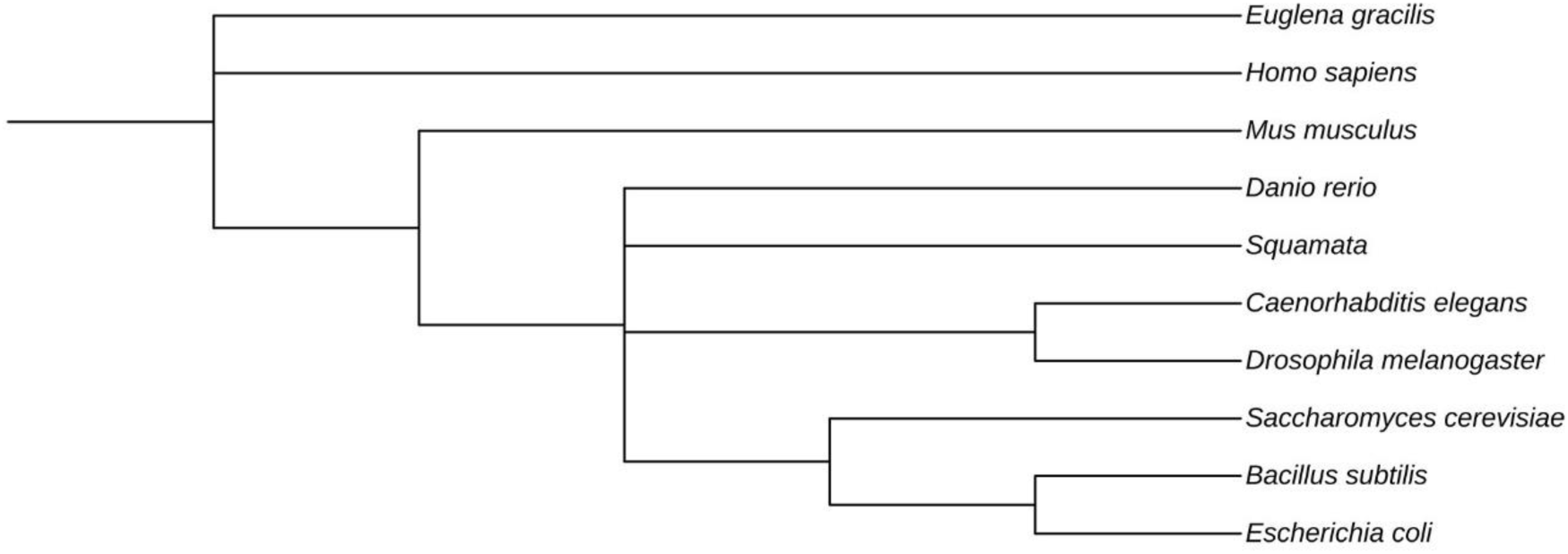
Phylogenetic tree representing the species: Homo sapiens, Mus musculus, Squamata (lizard), Danio rerio (zebrafish), Drosophila melanogaster, Caenorhabditis elegans, Escherichia coli (K12 MG1655), Saccharomyces cerevisiae, Bacillus subtilis (PY79), and Euglena gracilis.

**Table II.**
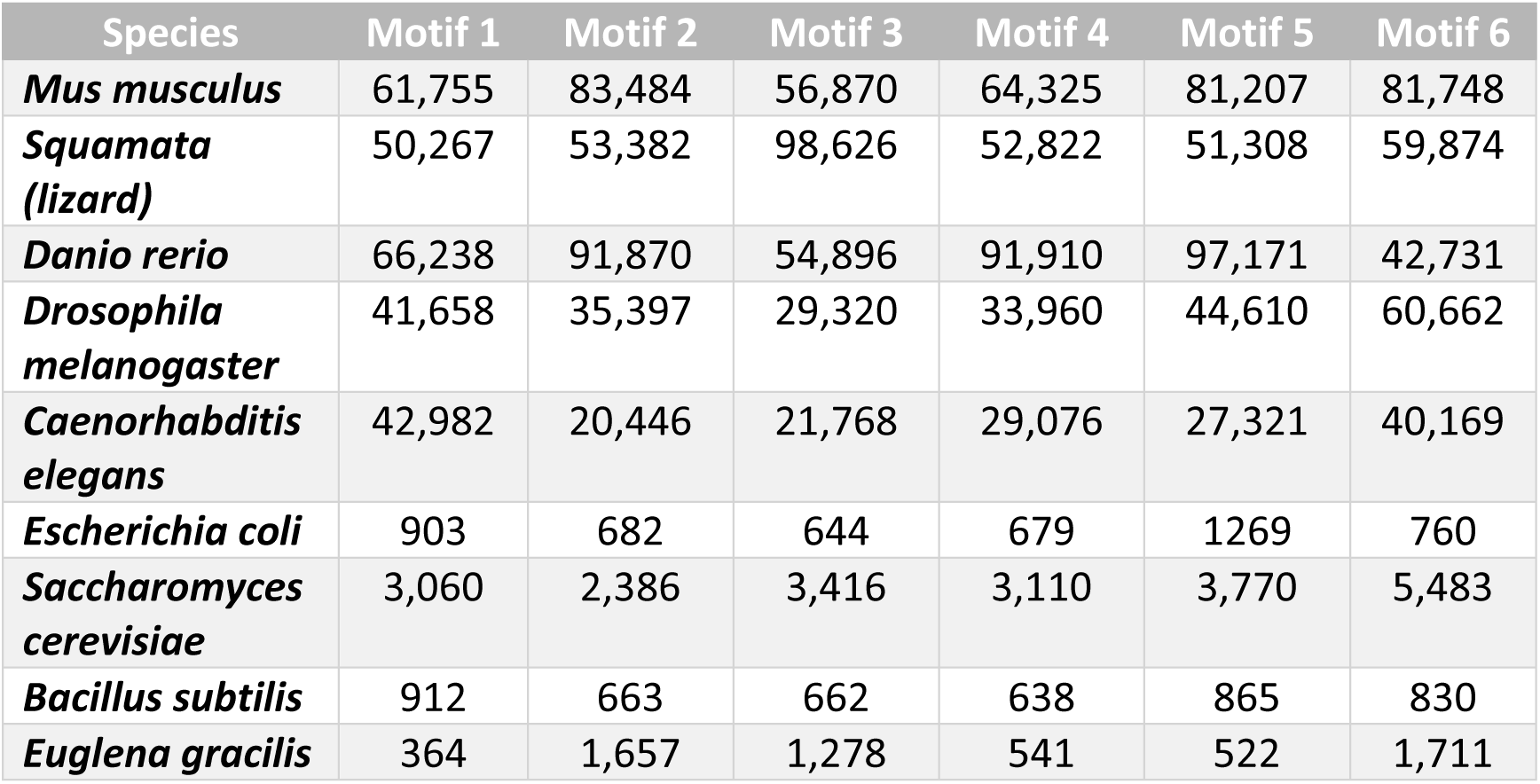
Occurrence frequency of the motifs in the genome sequences of the species Mus musculus, Squamata (lizard), Danio rerio (zebrafish), Drosophila melanogaster, Caenorhabditis elegans, Escherichia coli, Saccharomyces cerevisiae, Bacillus subtilis (PY79), and Euglena gracilis.

The genetic similarities between humans and the other organisms examined in our study show significant variation. For instance, humans and *Mus musculus* (mouse) share approximately 85% genetic similarity, reflecting a divergence of about 70 million years ago, making the mouse an essential model for human genetic research, particularly in disease mechanisms and therapeutic development (121). *Squamata*, a diverse group of reptiles, includes species that serve as critical models for studying thermoregulation, behavior, and reproductive strategies. Research on lizards and snakes has provided important insights into immunity, skin regeneration, and other physiological processes crucial for survival in varied environments (122). The genetic similarity between humans and different *Squamata* species varies considerably, with lizards sharing around 70% genetic similarity with humans, diverging approximately 250 million years ago. The six motifs identified in our study were also present in *Squamata*, suggesting their ancient evolutionary origins. *Danio rerio (zebrafish)*, with a genetic similarity to humans of about 70%, diverged around 400 million years ago and is an important model organism for developmental biology and gene function studies (123). *Drosophila melanogaster* (fruit fly), sharing roughly 60% of its genes with humans and diverging around 600 million years ago, is extensively used in genetic regulation, neurobiology, and cellular research (124). *Caenorhabditis elegans* (nematode), with around 40% genetic similarity to humans, has been instrumental in research on aging, developmental biology, and cellular processes (125). *Escherichia coli K12 MG1655*, although lacking direct gene homologs in humans, plays a crucial role in understanding fundamental gene regulation and bacterial genetics (126). *Saccharomyces cerevisiae* (yeast) shares approximately 23% of its genes with humans and diverged from humans over 1 billion years ago, making it a key organism for cell biology, genomics, and metabolism research (127). *Bacillus subtilis PY79*, a soil bacterium, has contributed significantly to our understanding of bacterial gene regulation and stress responses (128). Finally, *Euglena gracilis*, a single-celled eukaryote, is believed to share ancient nuclear receptor mechanisms with humans, offering insights into the early evolution of eukaryotic gene regulation (129).

The presence of the discovered motifs in species that diverged long before humans, including organisms with distinct genomes such as *Euglena gracilis* and *Saccharomyces cerevisiae*, combined with the lack of specificity in these motifs, suggests they may represent ancestral binding sites for the precursors of nuclear receptors. The conservation of these motifs across a broad evolutionary spectrum emphasizes their potential role in regulating gene expression via NR binding. Given that these motifs do not exhibit specificity for any particular NR, we propose that they may have functioned as general binding sites for ancestral NR precursors (130). Moreover, these motifs have a crucial role in the evolution of NR function by enabling NRs to interact with a broader range of binding sites, facilitating gene regulation in response to environmental and internal cues (119, 131).

The widespread presence of these motifs across different evolutionary lineages points to their ancient origins, likely predating the highly specific NRs found in modern organisms. In early eukaryotes, these motifs could have functioned as genomic binding sites for precursor forms of NRs, which were less specific in their binding preferences compared to the more specialized NRs we observe today (132). This generalization would have been essential in the early stages of NR evolution, allowing for flexible and robust gene regulation. These early, versatile genomic binding sites could have allowed primordial NR-like proteins to modulate gene expression effectively, even in the absence of the highly refined receptor-genome interactions seen in contemporary systems.

The conservation of these motifs across such diverse species also suggests that they may have acted as backup mechanisms, ensuring the continued regulation of essential genes when canonical NR binding sites, such as the glucocorticoid response element (GRE) for GR, were unavailable or dysfunctional (132). In this sense, these motifs may have served as a safety net, allowing the NR system to maintain gene expression and cellular functions even in the absence of optimal binding site availability. This backup role would have been particularly important in the early evolutionary stages, when the molecular machinery for precise gene regulation was still evolving (133).

As NRs became more specialized through evolution, the ancestral motifs likely persisted due to their role in maintaining regulatory flexibility. In modern organisms, these motifs may still contribute to the robustness of gene regulation, particularly in cases where the canonical NR-binding sites are compromised or unable to function. Their ability to interact with multiple NRs enhances the adaptability of the organism to different signaling environments, providing a more versatile and resilient regulatory system (133).

The preservation of these motifs across such a broad evolutionary spectrum underscores their importance in the evolutionary trajectory of gene regulation. These motifs may represent a foundational aspect of NR functionality, offering a mechanism that allowed early organisms to adapt to changing environments by ensuring continued gene regulation, even when specific receptor-binding sites were not fully evolved. The fact that these motifs can still be found in modern organisms suggests that they continue to play a role in gene regulation, particularly in situations where canonical NR interactions are suboptimal or unavailable (119).

This broader, more flexible role in gene regulation may have allowed organisms to thrive in variable environments and adapt to new challenges. The motifs’ conservation across both ancient and modern species highlights their functional importance and their potential as key elements in maintaining the adaptability of the NR signaling system throughout evolutionary history.

Today, numerous studies examine and describe the potential function of lncRNAs in the regulation of gene expression and their abilities to act as biomarkers or as pharmaceutical targets, or even therapeutic molecules. In this study, six conserved motifs were detected for the first time in the sequences of the lncRNAs studied which are suggested to participate in the regulatory function of lncRNAs that contain them. However, to fully understand the role of these motifs-and the lncRNAs containing them-in gene regulation, future research will need to explore their functional significance in greater detail. Experimental studies could confirm whether these motifs act as alternative binding sites for NRs or other transcription factors. Additionally, further investigation into how these motifs contribute to the regulation of gene expression in both ancient and modern species will provide valuable insights into the evolution of NR-mediated gene regulation and its ongoing role in cellular adaptability.

## Acknowledgments

Not applicable.

## Data Availability Statement

Not applicable.

## Authors’ contributions

DV conceived the current study. KP, LP, GPC, EE, and DV wrote, drafted, revised, edited, and reviewed the manuscript. All authors have read and approved the final manuscript. Data sharing is not applicable.

## Ethics approval and consent to participate

Not applicable.

## Patient consent for publication

Not applicable.

