## Supplementary material for "Role of lncRNAs related to NRs in the regulation of gene expression": Data S1

1. **Algorithm for the first ID type:**

function [S] = ID (nickname)

S='';

try

S = getgenbank(nickname');

catch

url1=['https://www.ncbi.nlm.nih.gov/search/all/?term=' nickname];

data = webread(url1);

name=data(strfind(data,'pageData[''itemUid''] = "'):strfind(data,'pageData[''itemUid''] = "')+40);

pointer1=strfind(name,'"');

if isempty(pointer1)==0

name=name(pointer1(1)+1:pointer1(2)-1);

if isequal(name,'None')==0 && isempty(name)==0

url1=['https://www.ncbi.nlm.nih.gov/gene/' name];

data = webread(url1);

name=data(strfind(data,'<p><a href="/nuccore/'):strfind(data,'<p><a href="/nuccore/')+40);

pointer1=strfind(name,'/');

pointer2=strfind(name,'.');

if isempty(pointer1)==0 && isempty(pointer2)==0

name=name(pointer1(2)+1:pointer2(1)-1);

try

S = getgenbank(name);

catch

end

end

end

end

end

2. **Algorithm for the second ID type:**

function [S]=ID (nickname)

S='';

try

S = getgenbank(nickname');

catch

url1=['https://www.ncbi.nlm.nih.gov/gene/?term=' nickname];

data = webread(url1);

name=data(strfind(data,'<p><a href="/nuccore/'):strfind(data,'<p><a href="/nuccore/')+40);

pointer1=strfind(name,'/');

pointer2=strfind(name,'.');

if isempty(pointer1)==0 && isempty(pointer2)==0

name=name(pointer1(2)+1:pointer2(1)-1);

try

S = getgenbank(name);

catch

end

end

end

3. **Algorithm for the third ID type:**

function [S] = ID (nickname)

S='';

try

S = getgenbank(nickname);

catch

end

4. **Algorithm for the fourth ID type**

function [fasta]=ID (nickname)

try

url1=['https://lncipedia.org/db/search?search_id=' nickname];

data = webread(url1);

pointer1=strfind(data,'<td>');

pointer2=strfind(data,'</td>');

tname=data(pointer1(1)+5:pointer2(1)-2);

tname=tname(1:end-1);

pointer1=strfind( tname,'pt/');

pointer2=strfind( tname,'">');

tname2=tname(pointer1(1)+3:pointer2(1)-1);

url1=['https://lncipedia.org/db/transcript/' tname2];

data = webread( url1);

name=data(strfind(data,'<textarea style="width: 100%; height: 200px;">'):strfind(data,'</textarea>')-1);

pointer1=strfind(name,'">');

fasta.Header=[nickname '|' tname2];

fasta.Sequence=name(pointer1(1)+2:end);

catch

fasta='';

end
