## Supplementary material for "Role of lncRNAs related to NRs in the regulation of gene expression": Data S2

countera=1;

counterb=1;

for i=166:length(data)

i

clear fasta

try

[fasta]=4th ID type(data{i,1});

catch

end

if isempty(fasta)==0

lnc(countera)=fasta;

countera=countera+1;

elseif isempty(fasta)==1

try

clear S

[S]=3rd ID type(data{i,1});

if isempty(S)==1

[S]=1st ID type(data{i,1});

elseif isempty(S)==1

[S]=2nd ID type(data{i,1});

end

catch

end

if isempty(S)==1

n_lnc(counterb).name=data{i,1};

n_lnc(counterb).name

counterb=counterb+1;

elseif isempty(S)==0

try

lnc(countera).Header= [ data{i,1} '|' S(1).Accession];

lnc(countera).Sequence=upper(S(1).Sequence);

lnc(countera)

countera=countera+1;

catch

end

end

end

end

fastawrite('NR_LNC_MR_GR.fasta',lnc)

fileIDA2 = fopen('NR_LNC_Not_found.txt','w');

for i=1:172

fprintf(fileIDA2,'%s\n',n_lnc(i).name);

end

fclose(fileIDA2);

save('LNC_NR_RESULTS')
