## Supplementary material for "Role of lncRNAs related to NRs in the regulation of gene expression": Data S3

*counter=1;*

*for i=****first result’s row:last result’s row***

*aa=split(data{i,1},'_');*

*s(counter).chromosome=aa{1};*

*s(counter).place=data{i,3};*

*counter=counter+1;*

*end*

*clear A1 A B C D*

*for i=1:length(s)*

*if i==1*

*A{i}=s(i).chromosome(4:end);*

*B=[1];*

*C=str2num(s(i).place);*

*D=str2num(s(i).place)+200000;*

*else*

*A{i}=s(i).chromosome(4:end);*

*B=[B 1];*

*C=[C , str2num(s(i).place)];*

*D=[D , (str2num(s(i).place)+200000)];*

*end*

*end*

*A1=char(A);*

*cna_struct = struct('Chromosome', A1,...*

*'CNVType', B,...*

*'Start', C,...*

*'End', D);*

*figure2=chromosomeplot(hs_cytobands, 'cnv', cna_struct, 'unit', 2);*
