## Supplementary figures and images for "Role of lncRNAs related to NRs in the regulation of gene expression"

### Figure S1

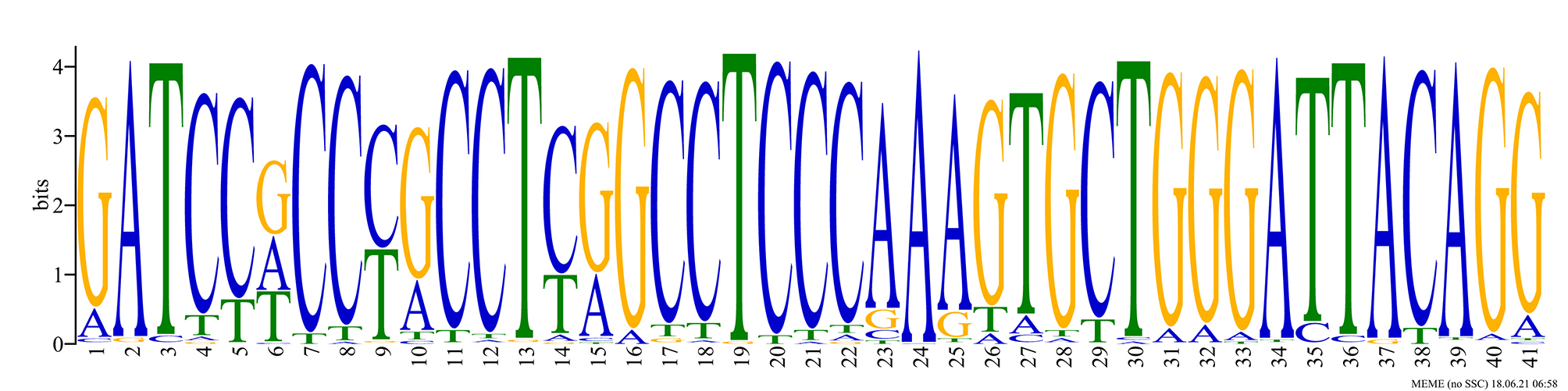

### Figure S2

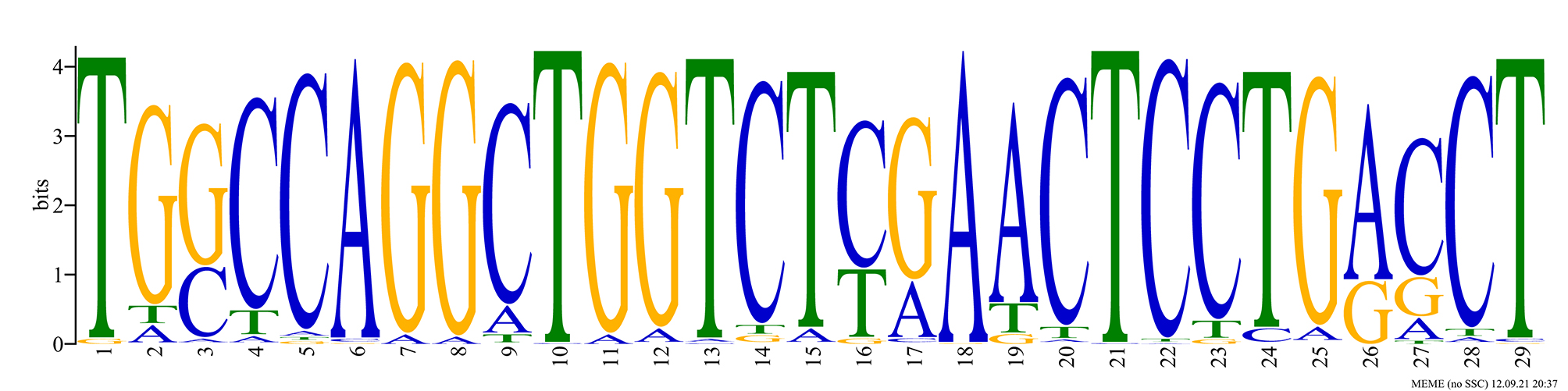

### Figure S3

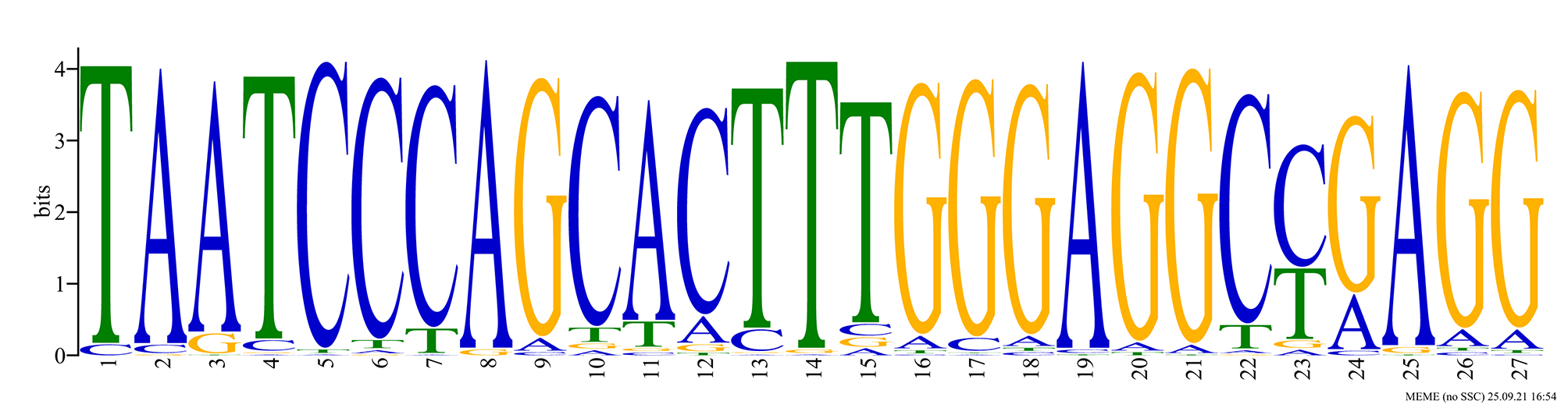

### Figure S4

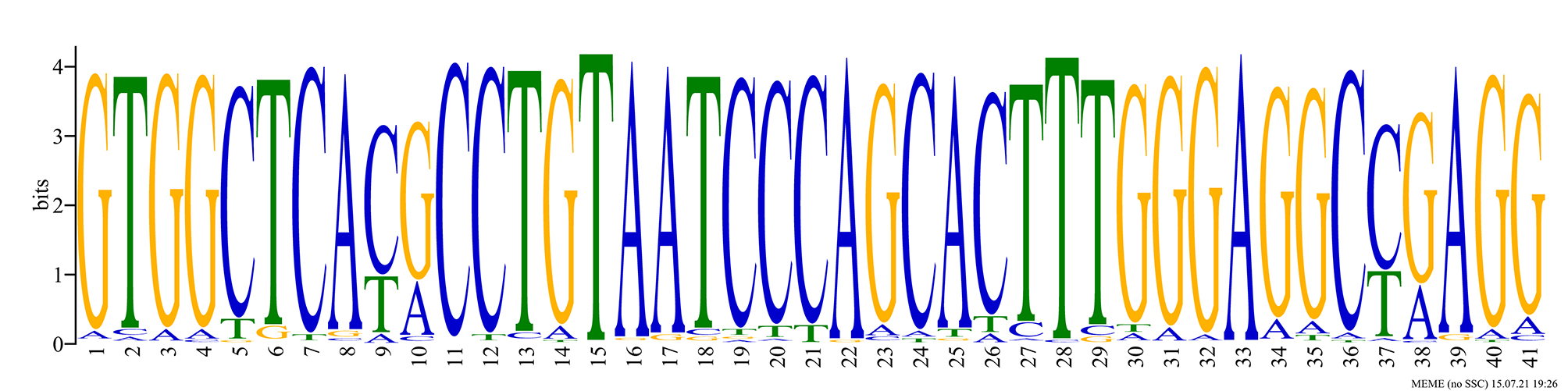

### Figure S5

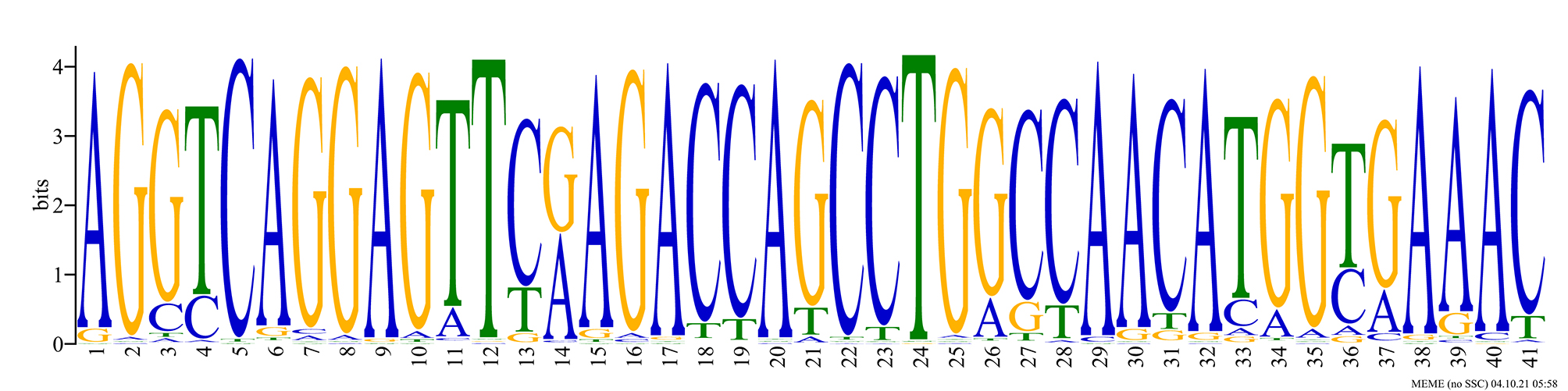

### Figure S6

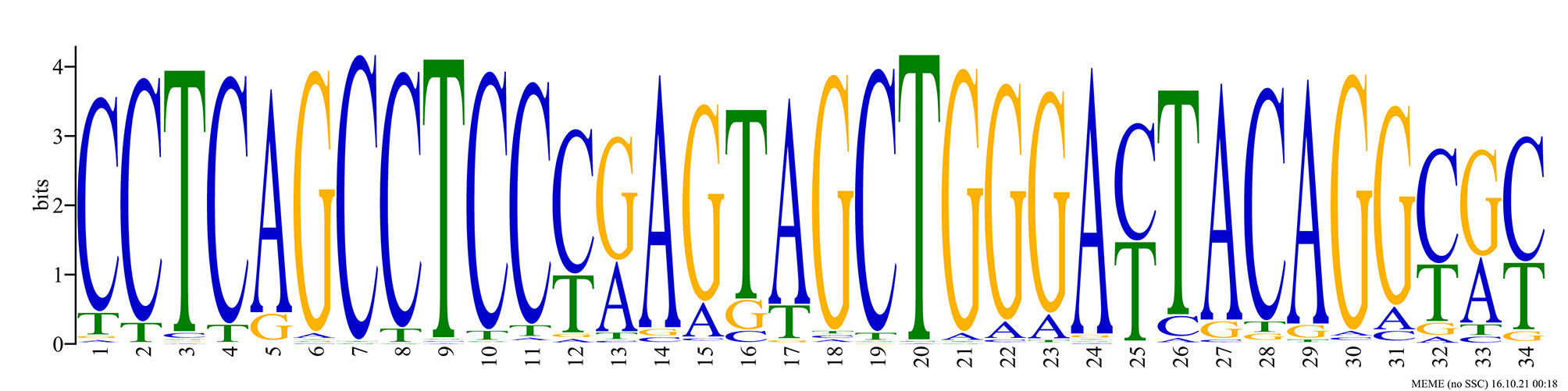

### Figure S7

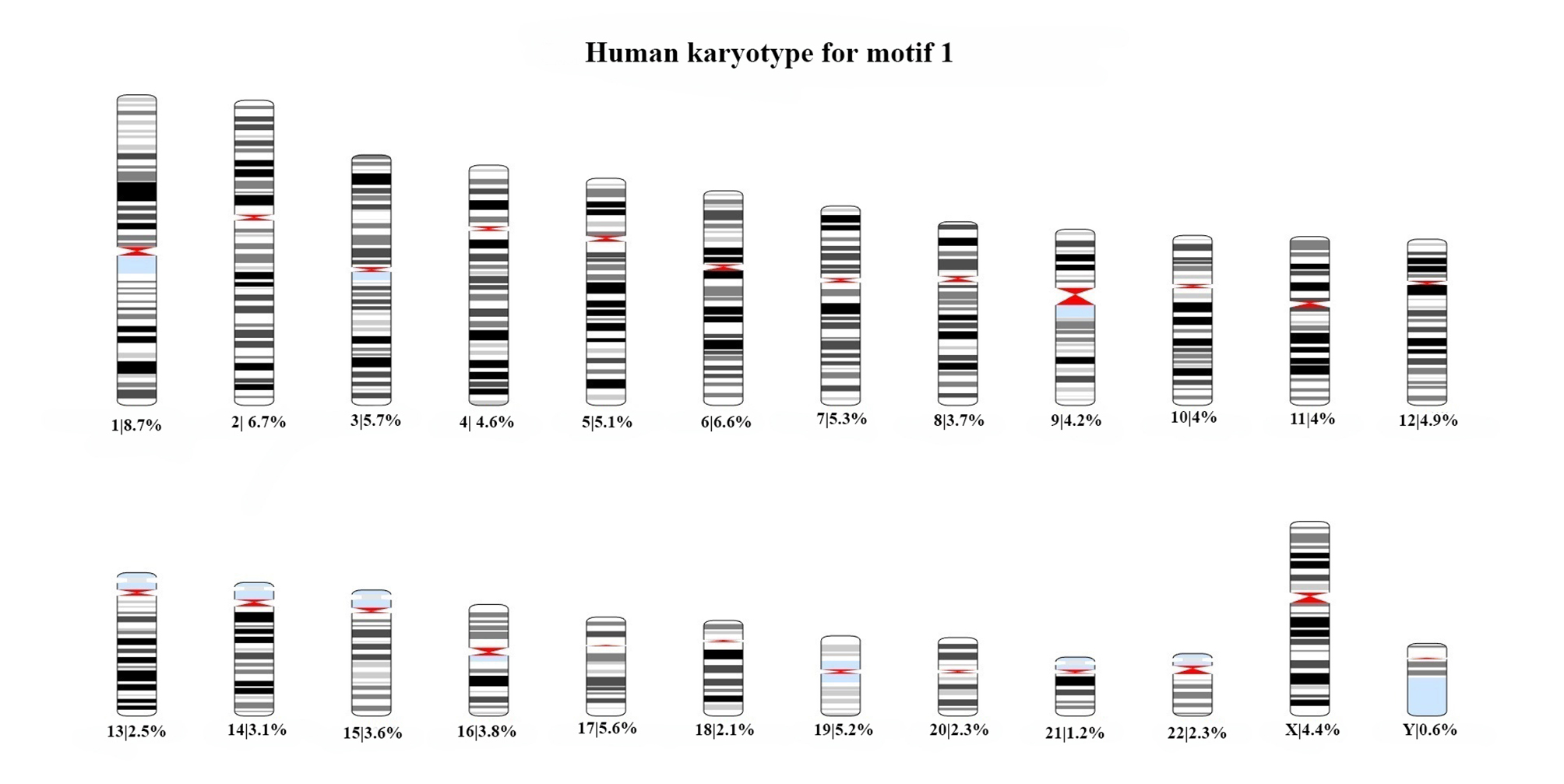

### Figure S8

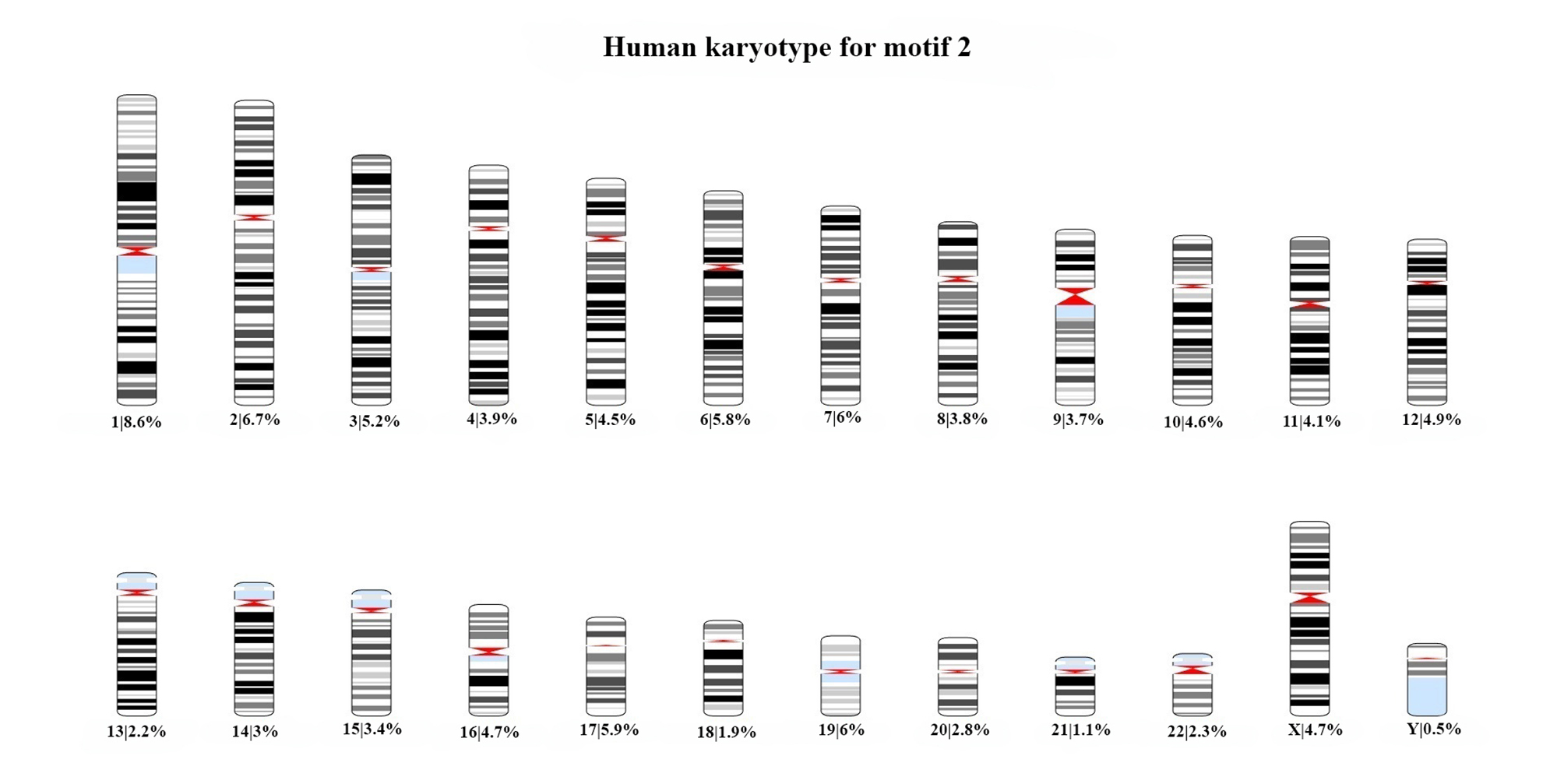

### Figure S9

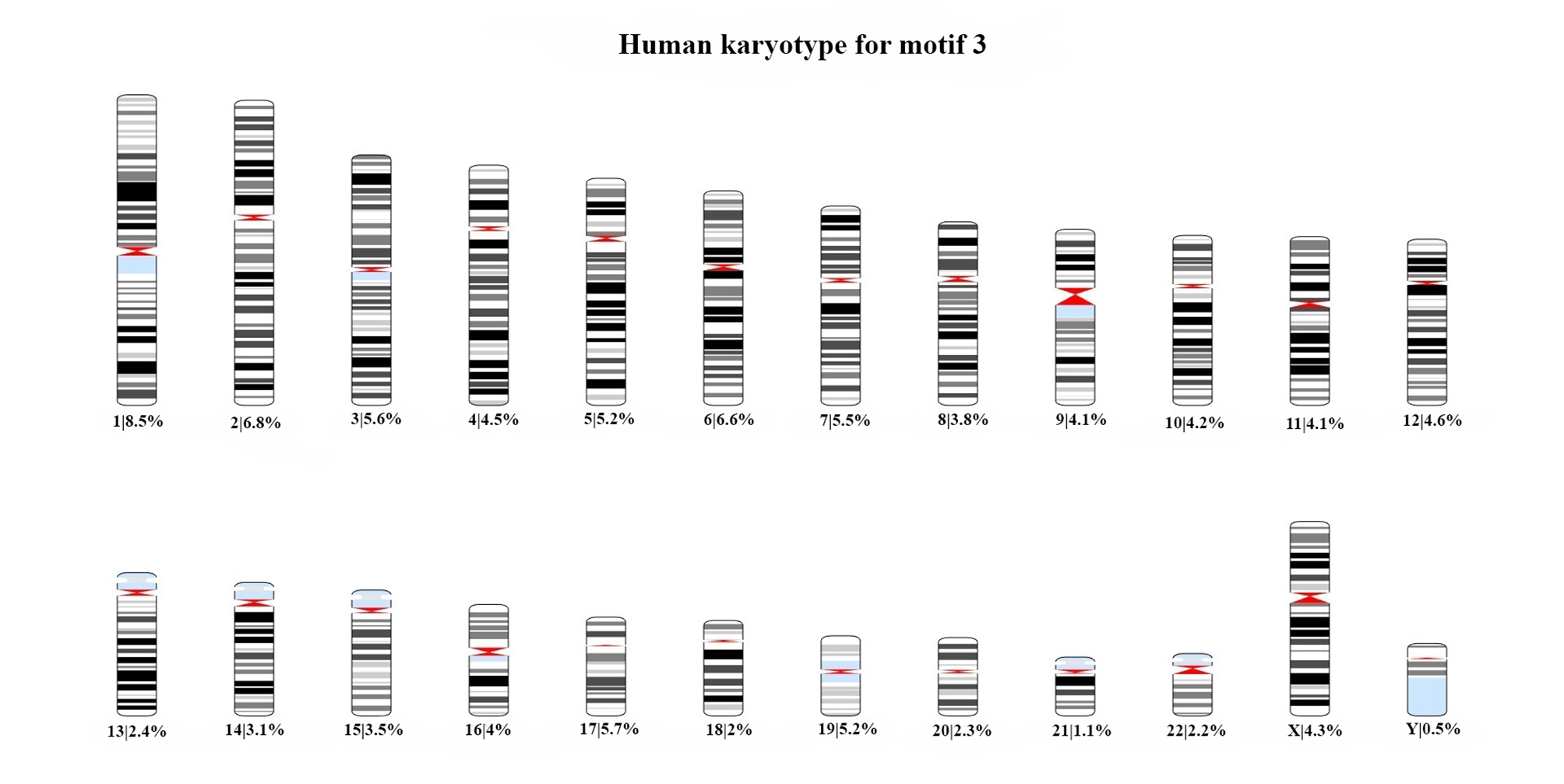

### Figure S10

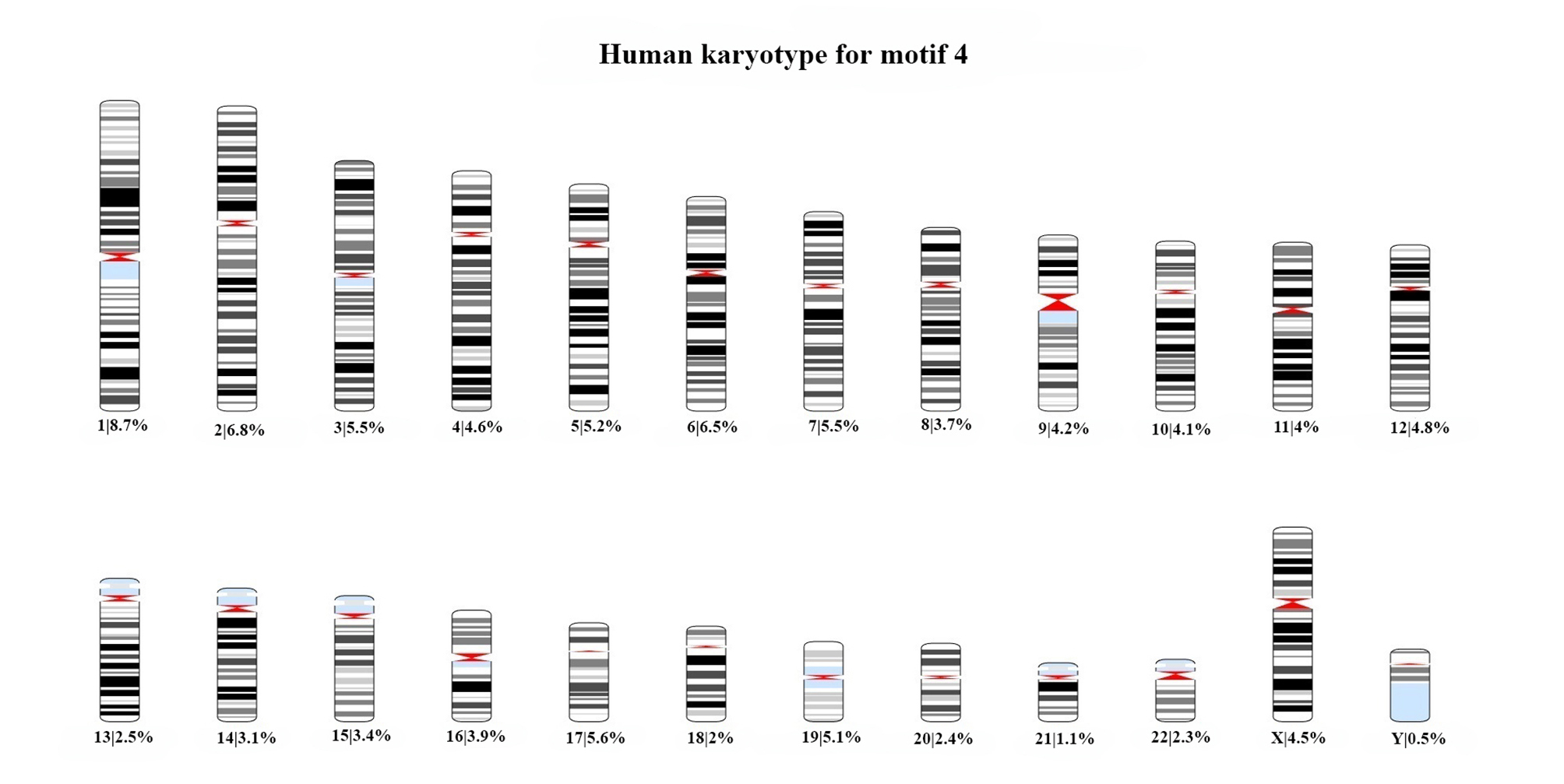

### Figure S11

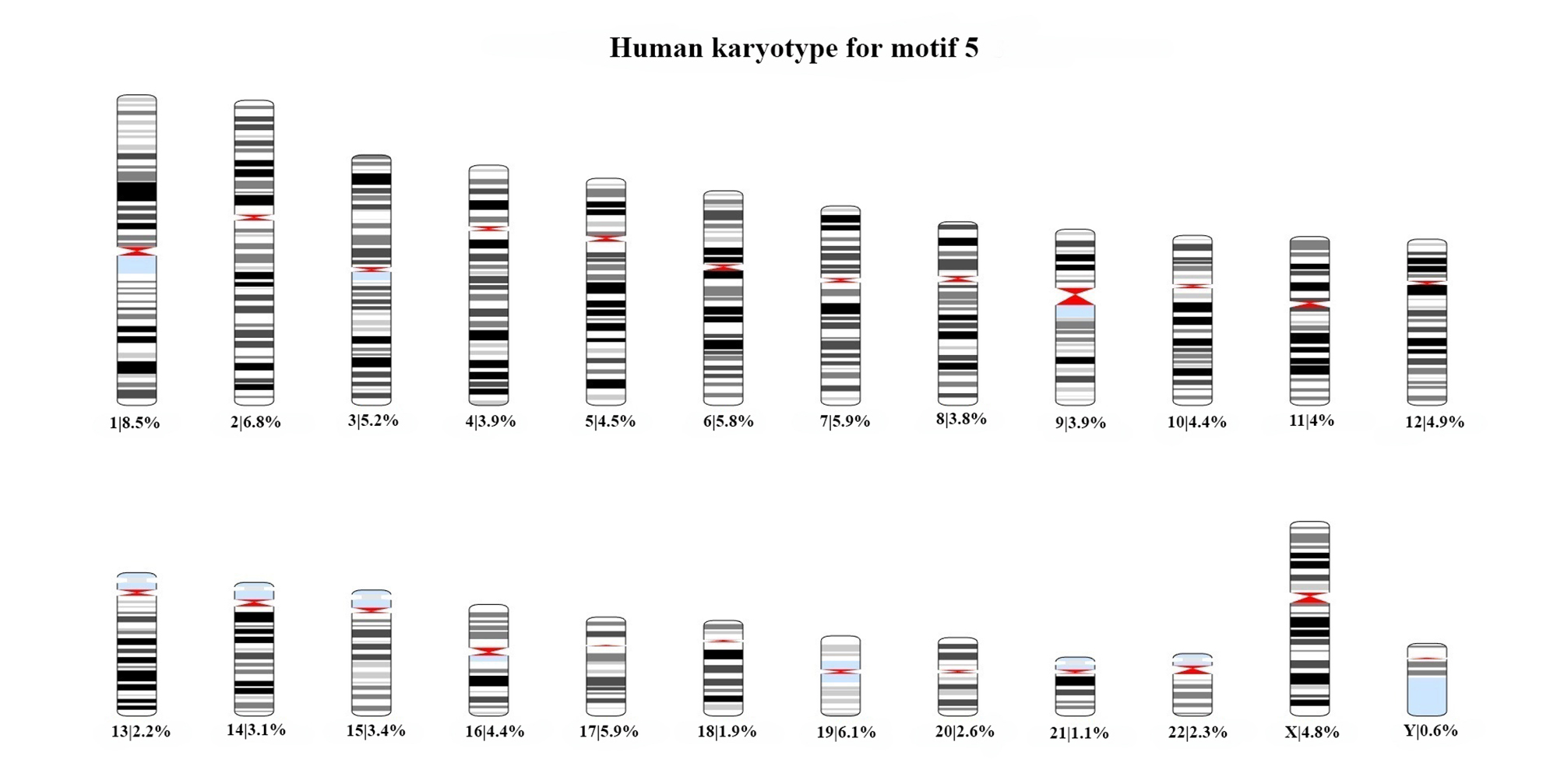

### Figure S12

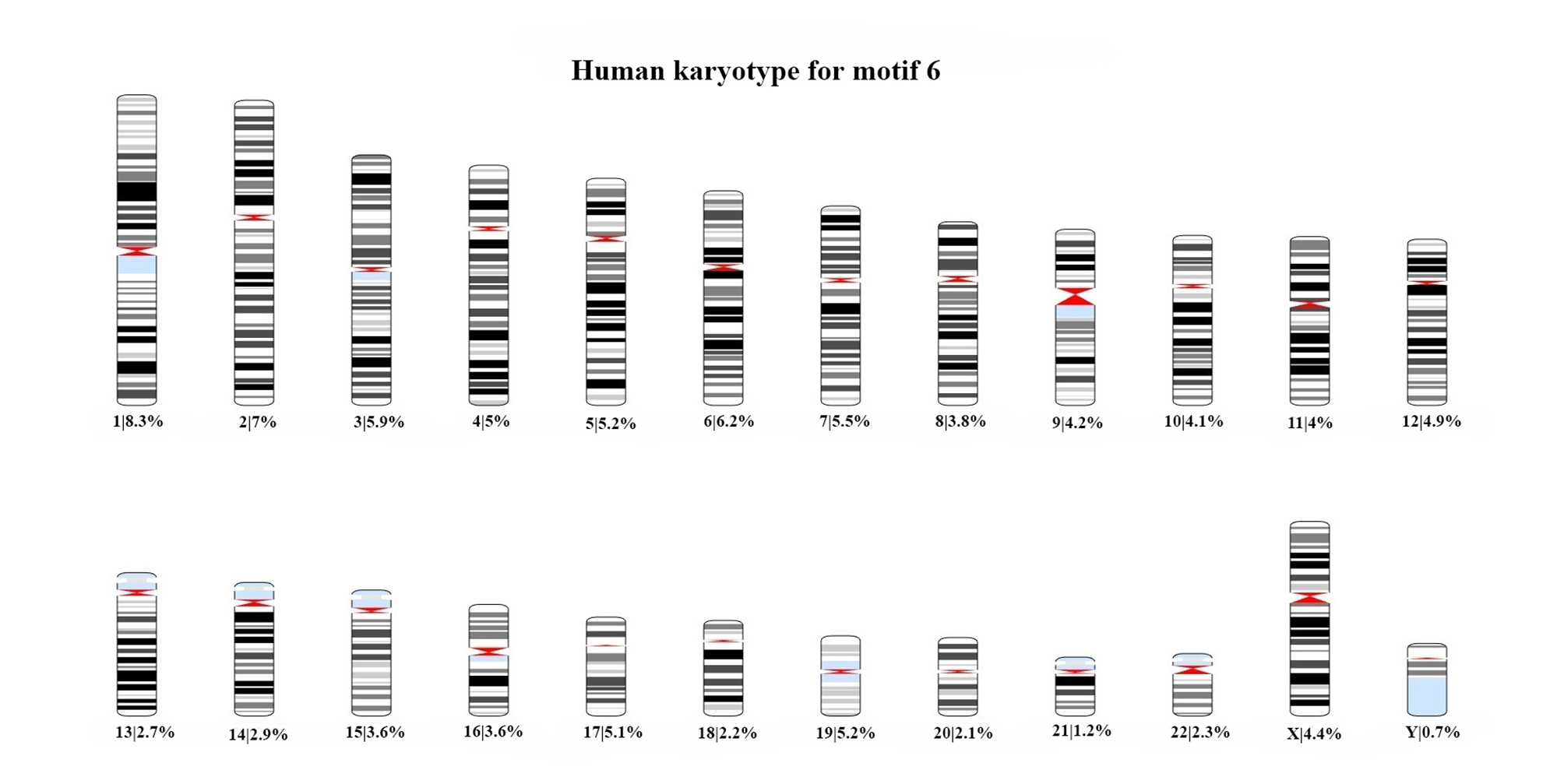

### Figure S13

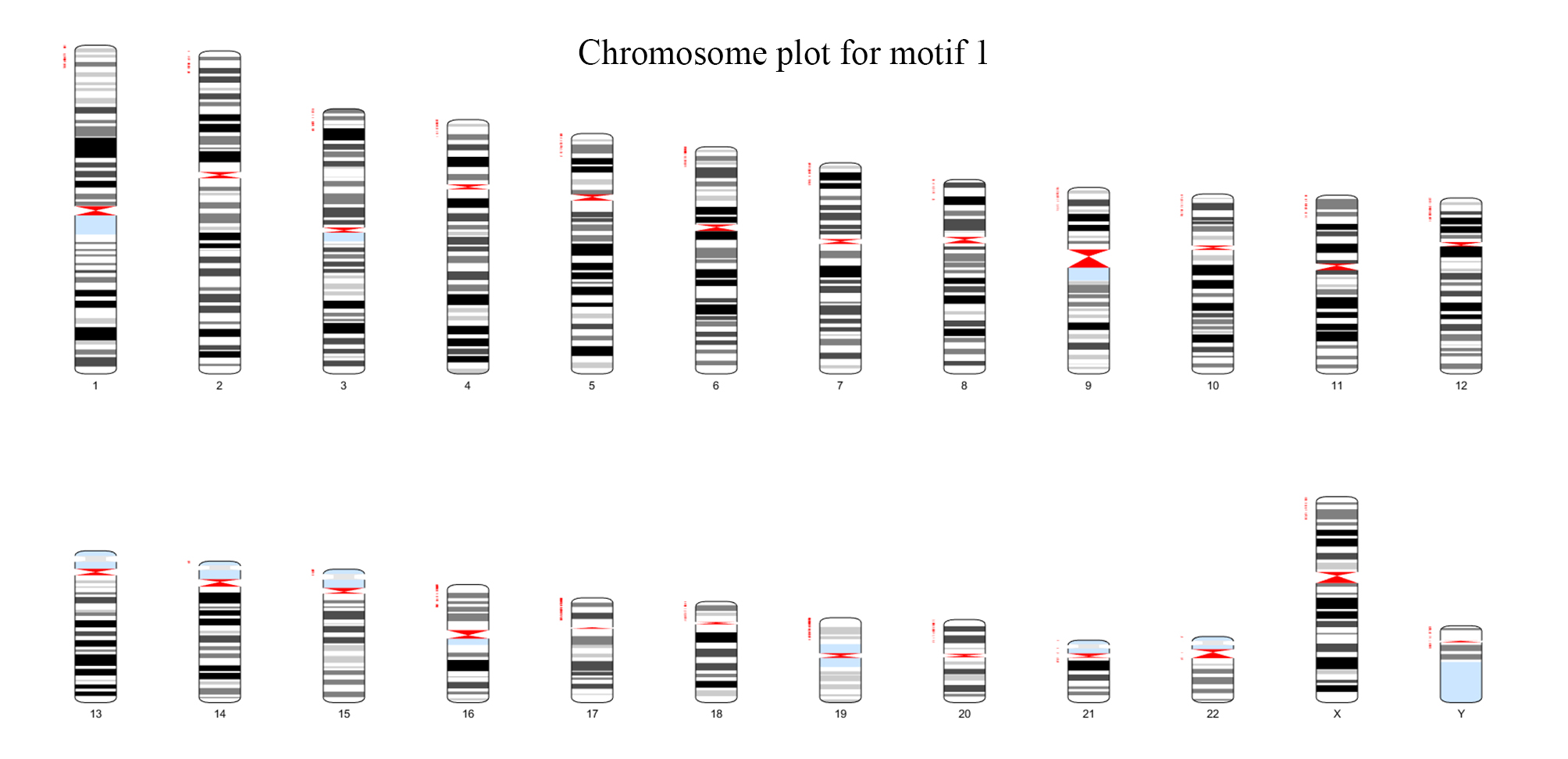

### Figure S14

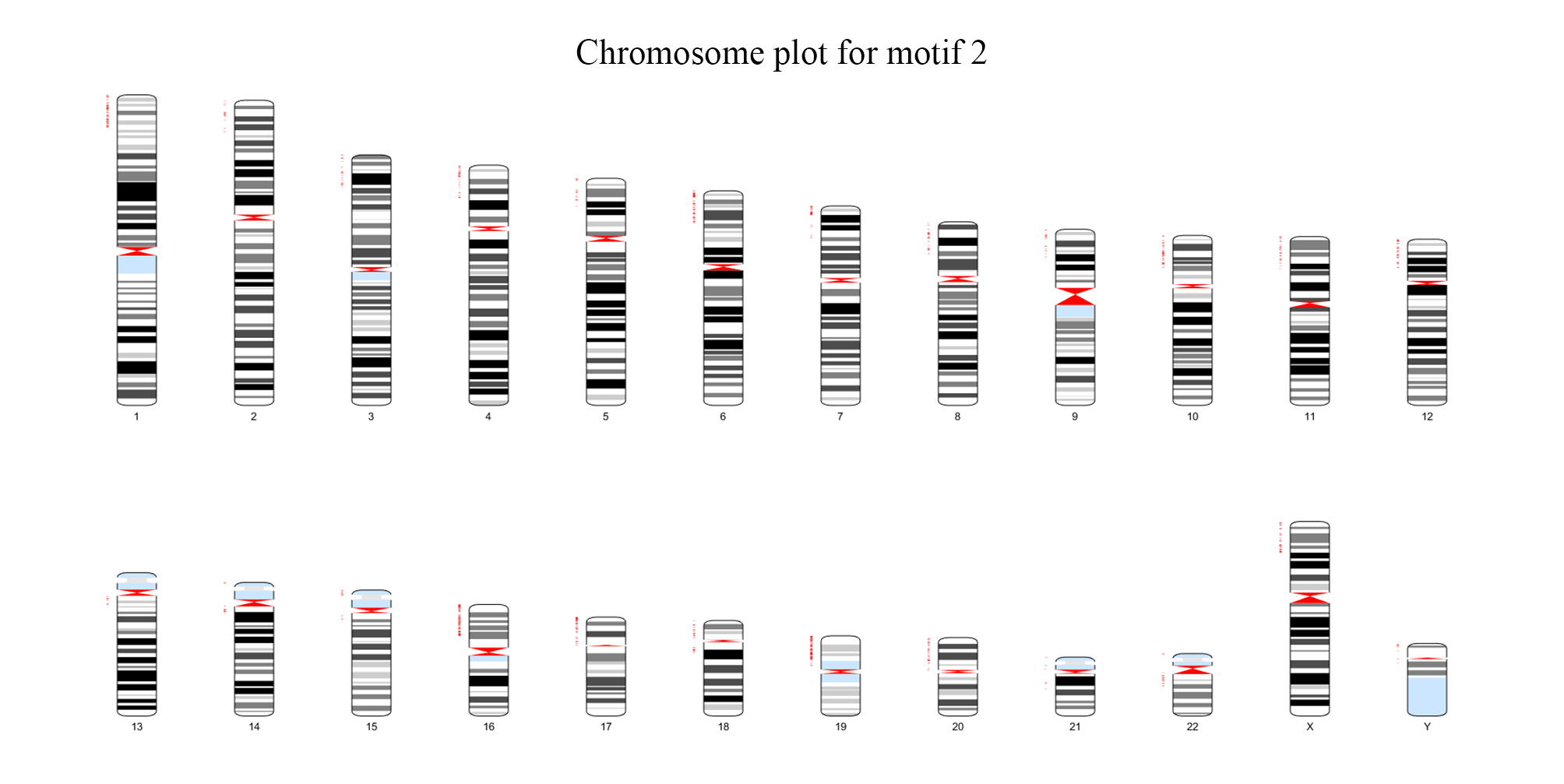

### Figure S15

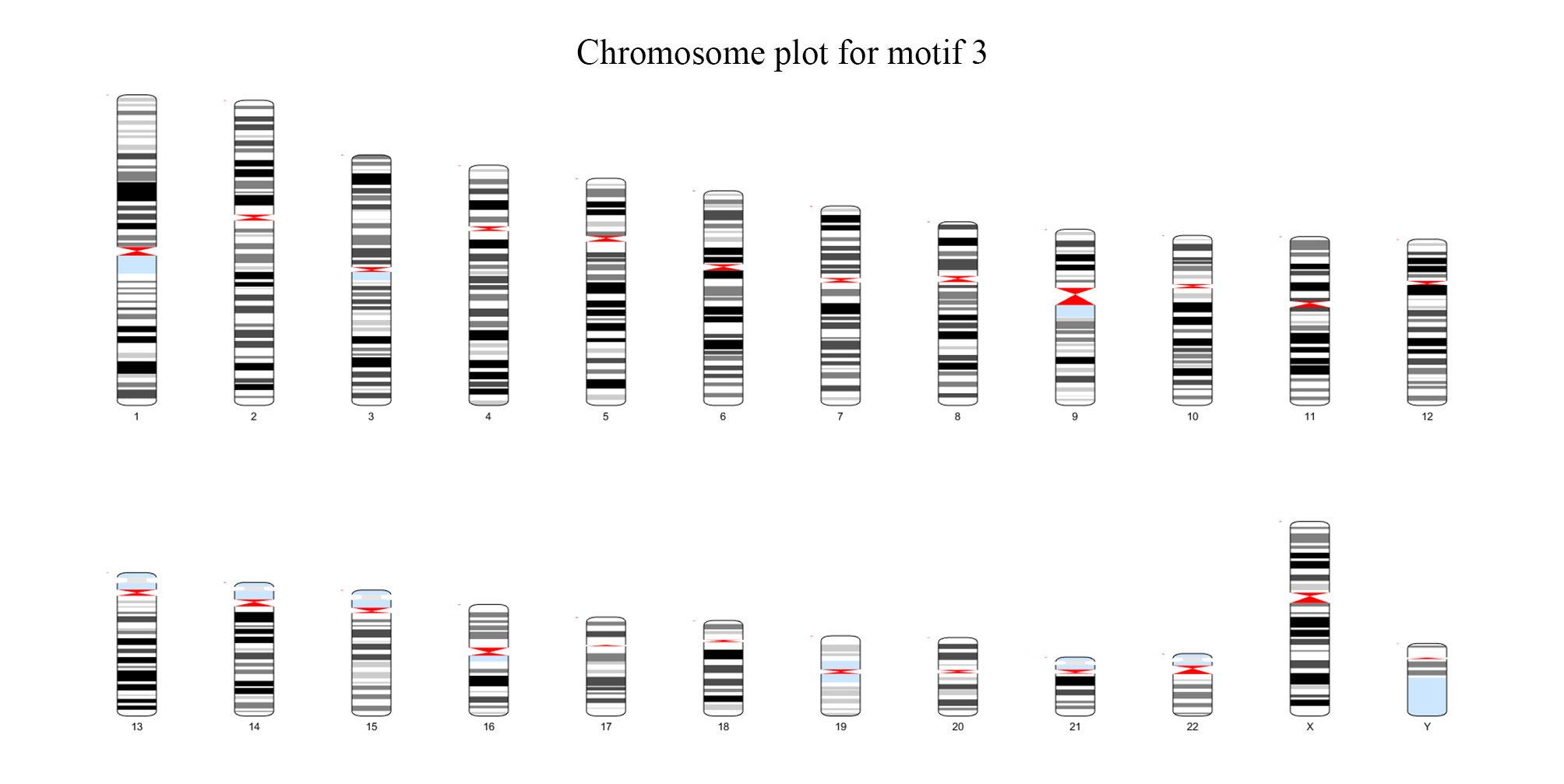

### Figure S16

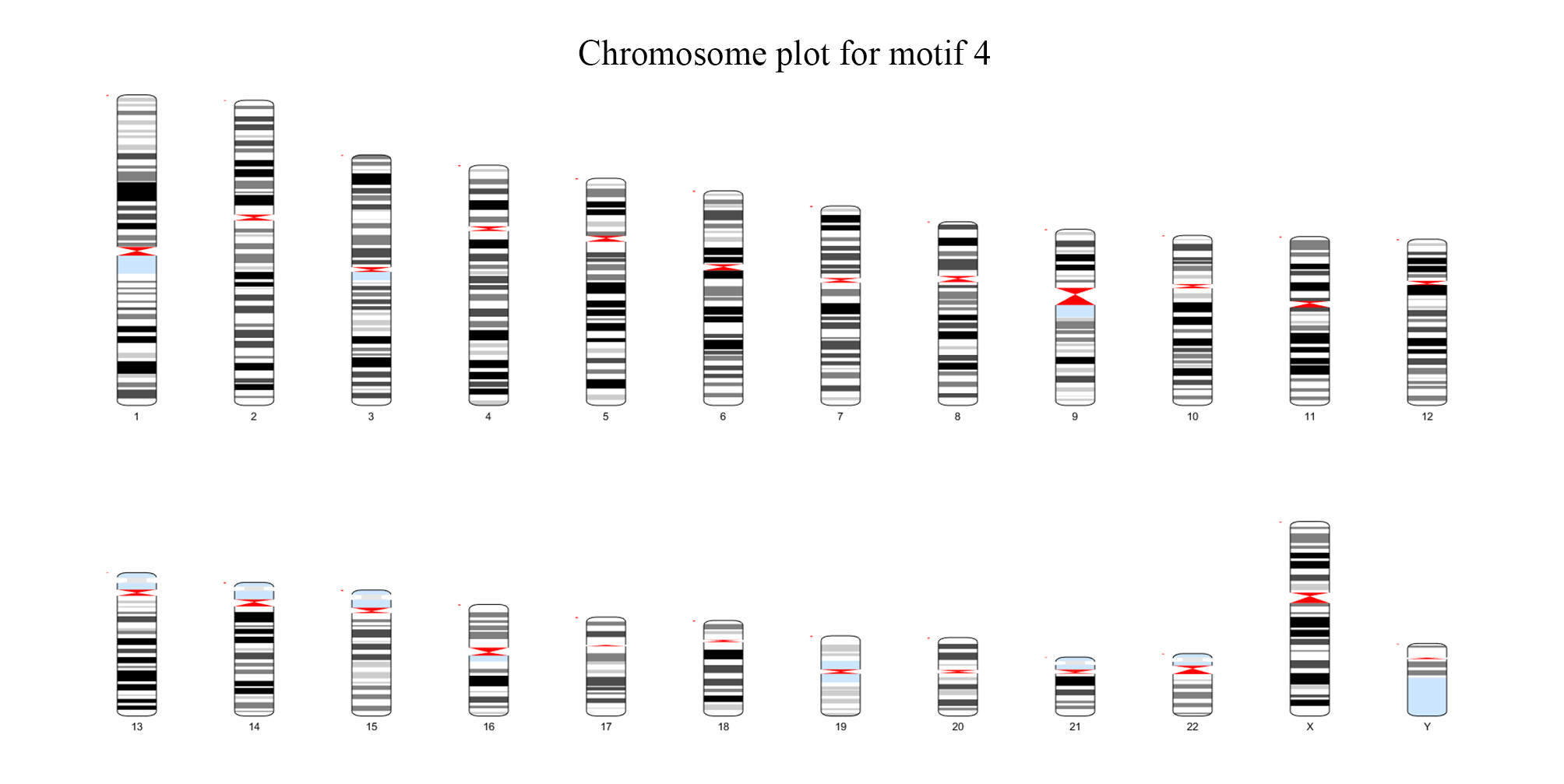

### Figure S17

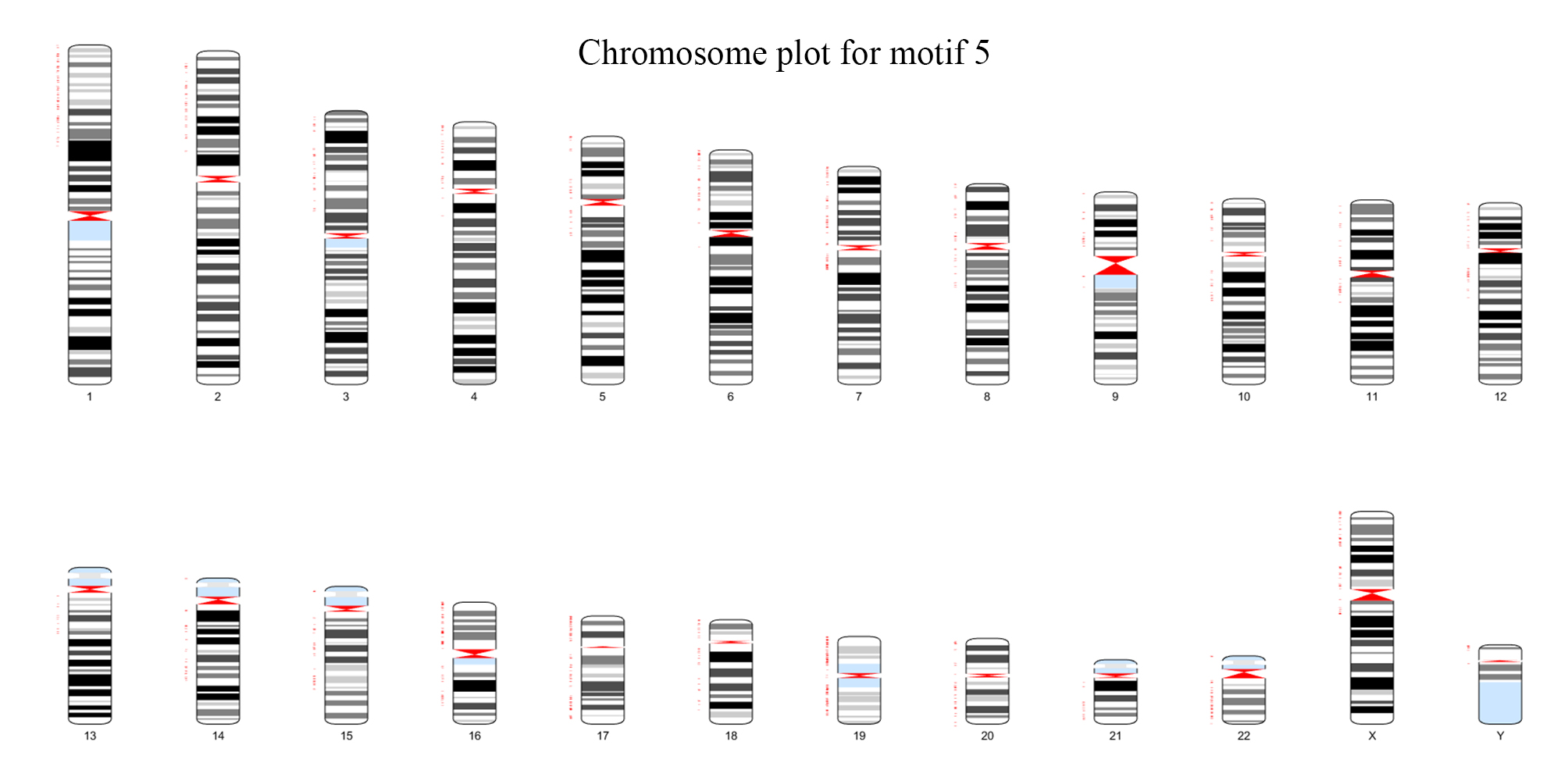

### Figure S18

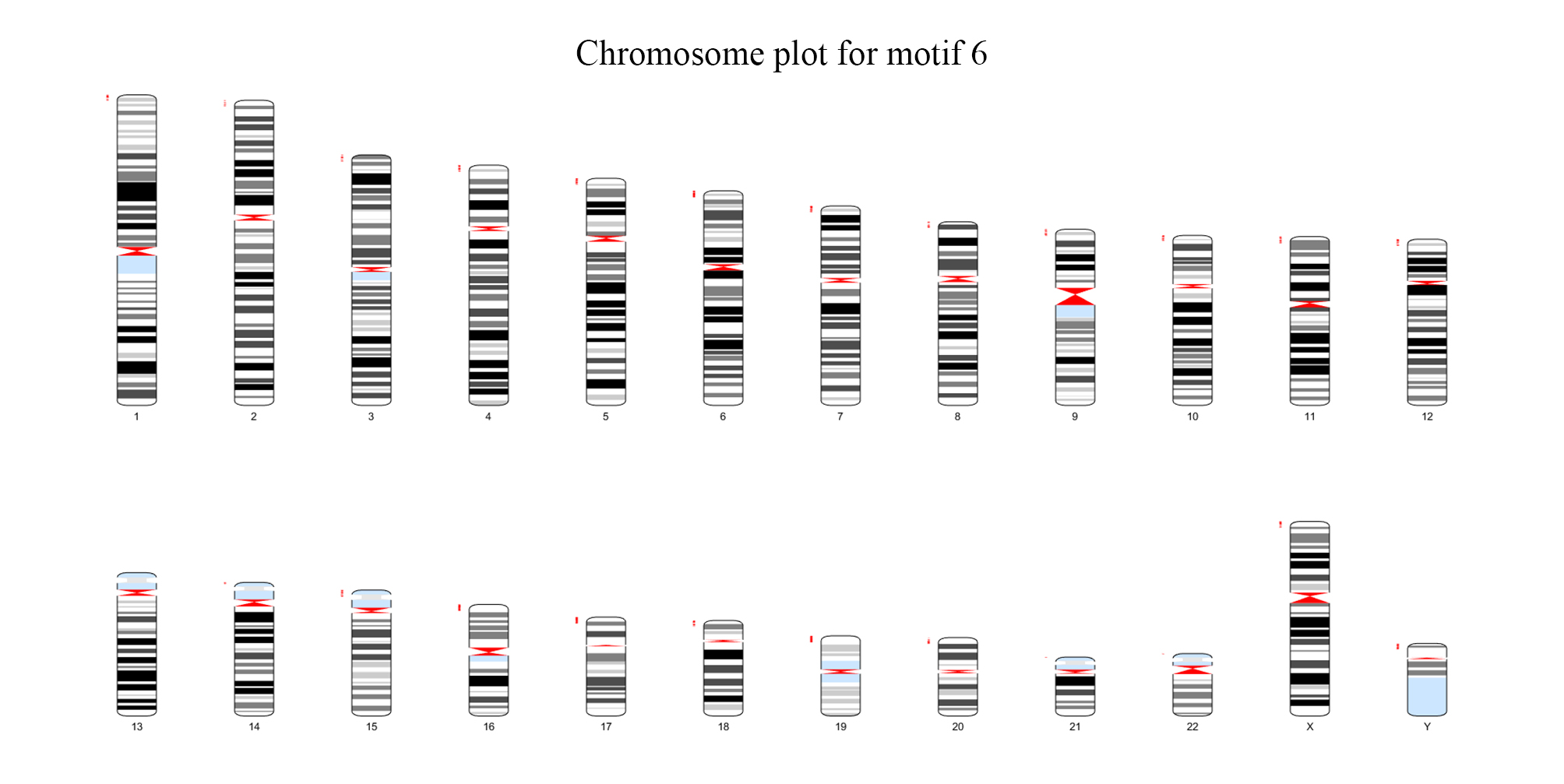

### Figure S19

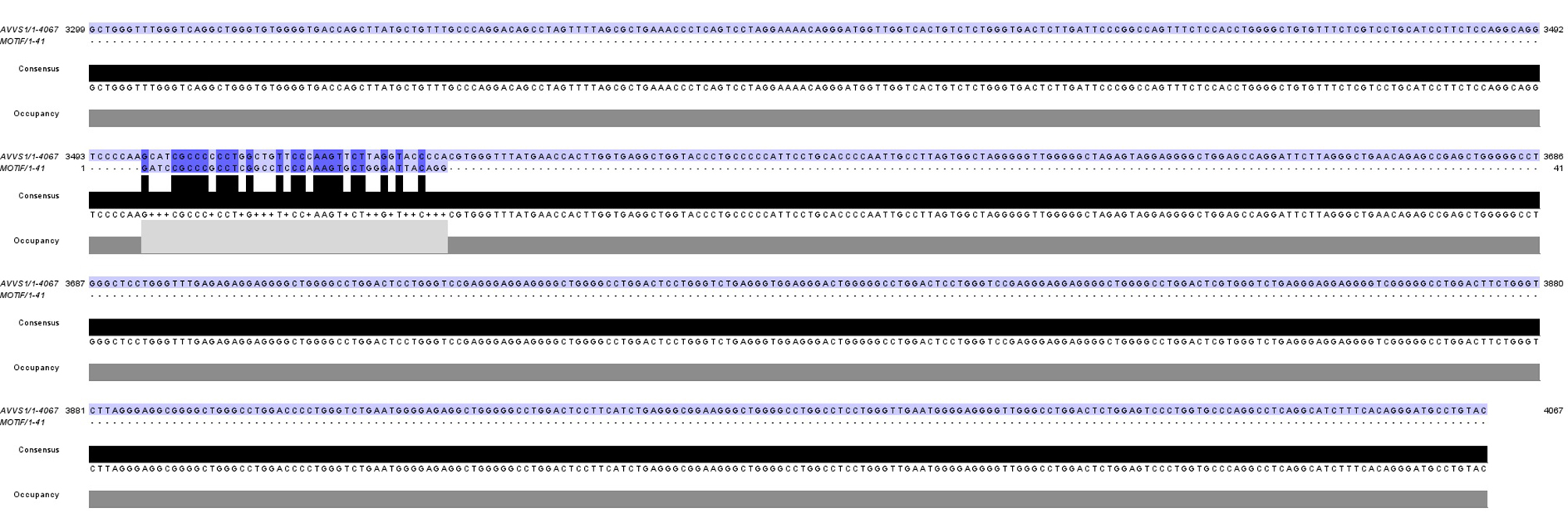

### Figure S20

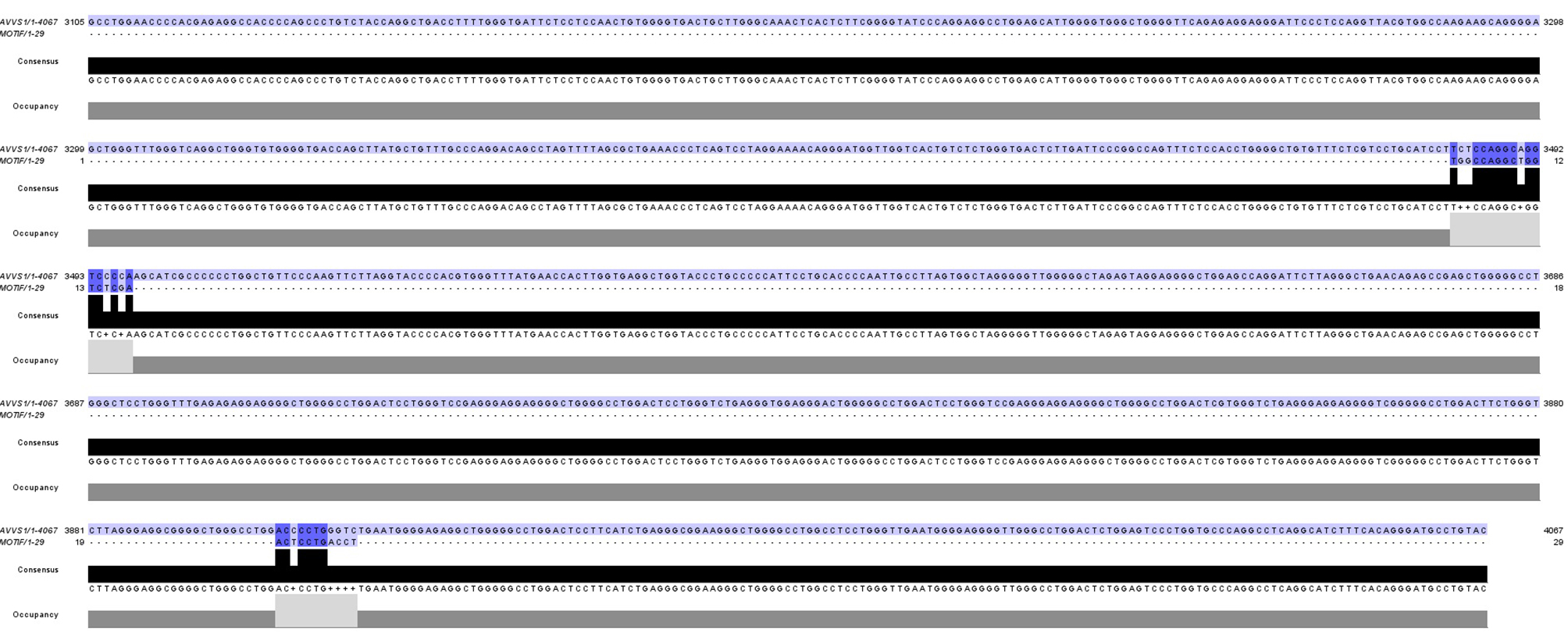

### Figure S21

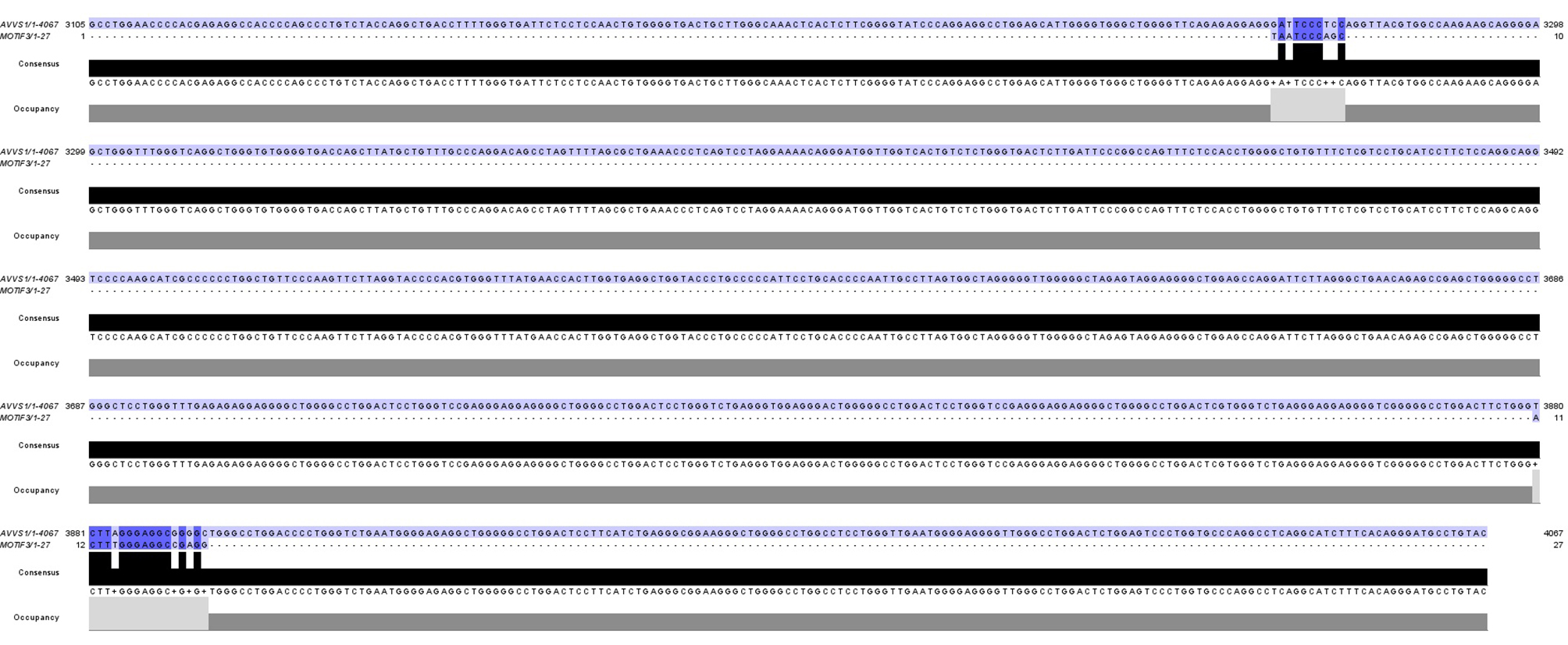

### Figure S22

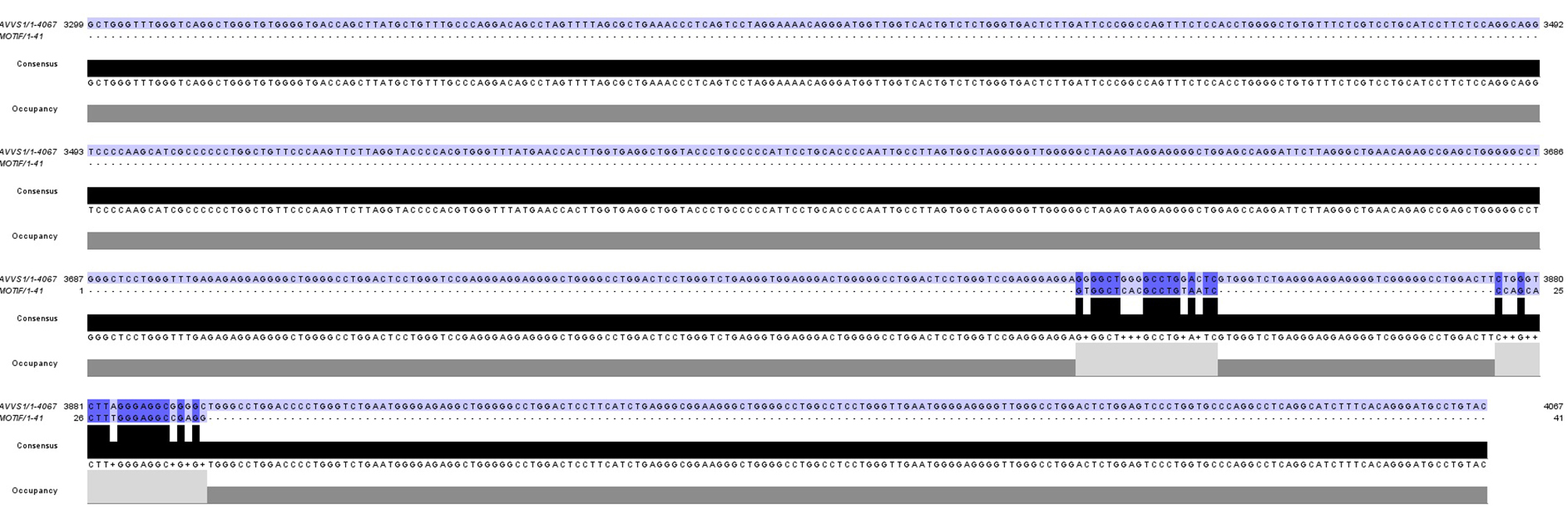

### Figure S23

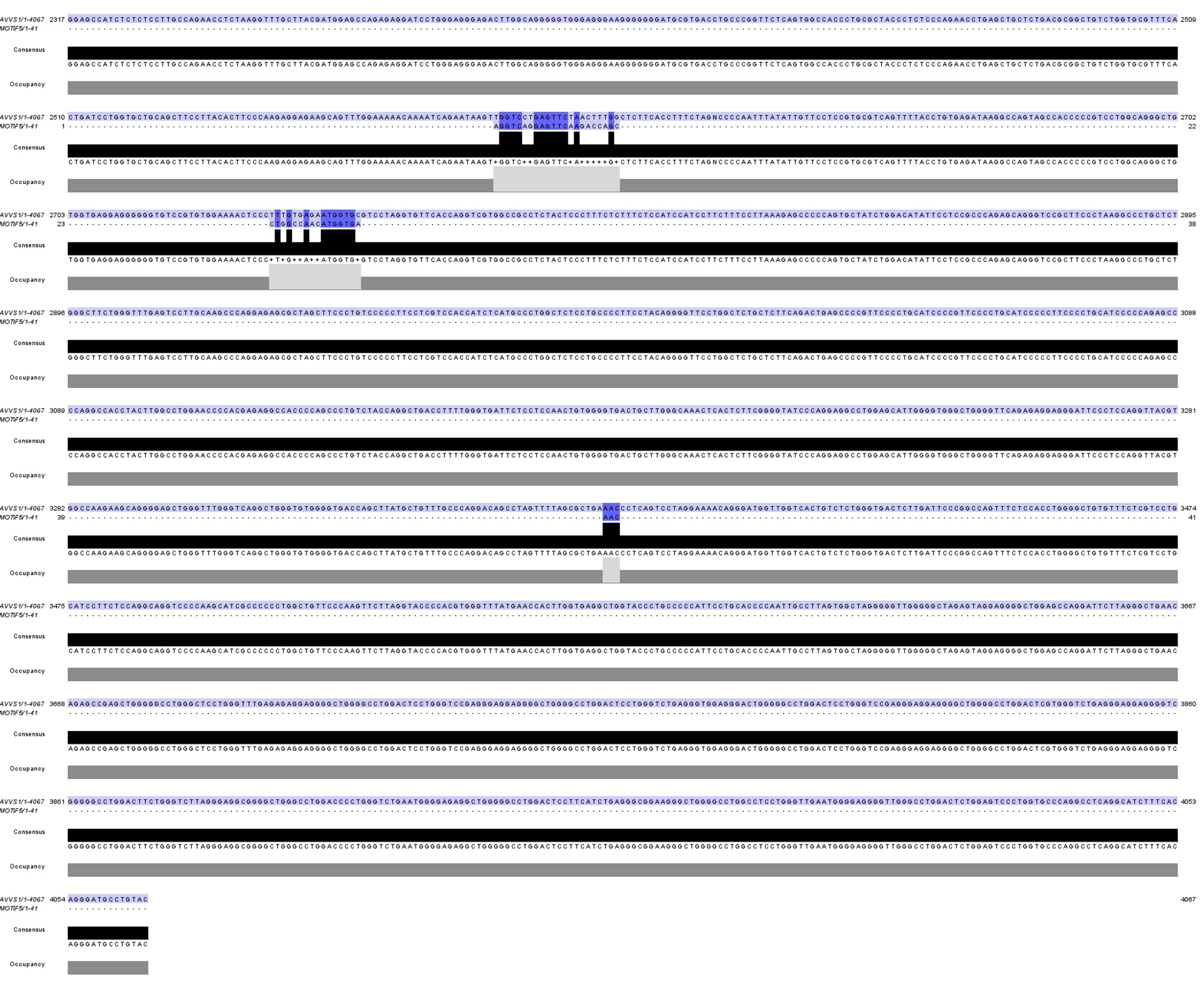

### Figure S24

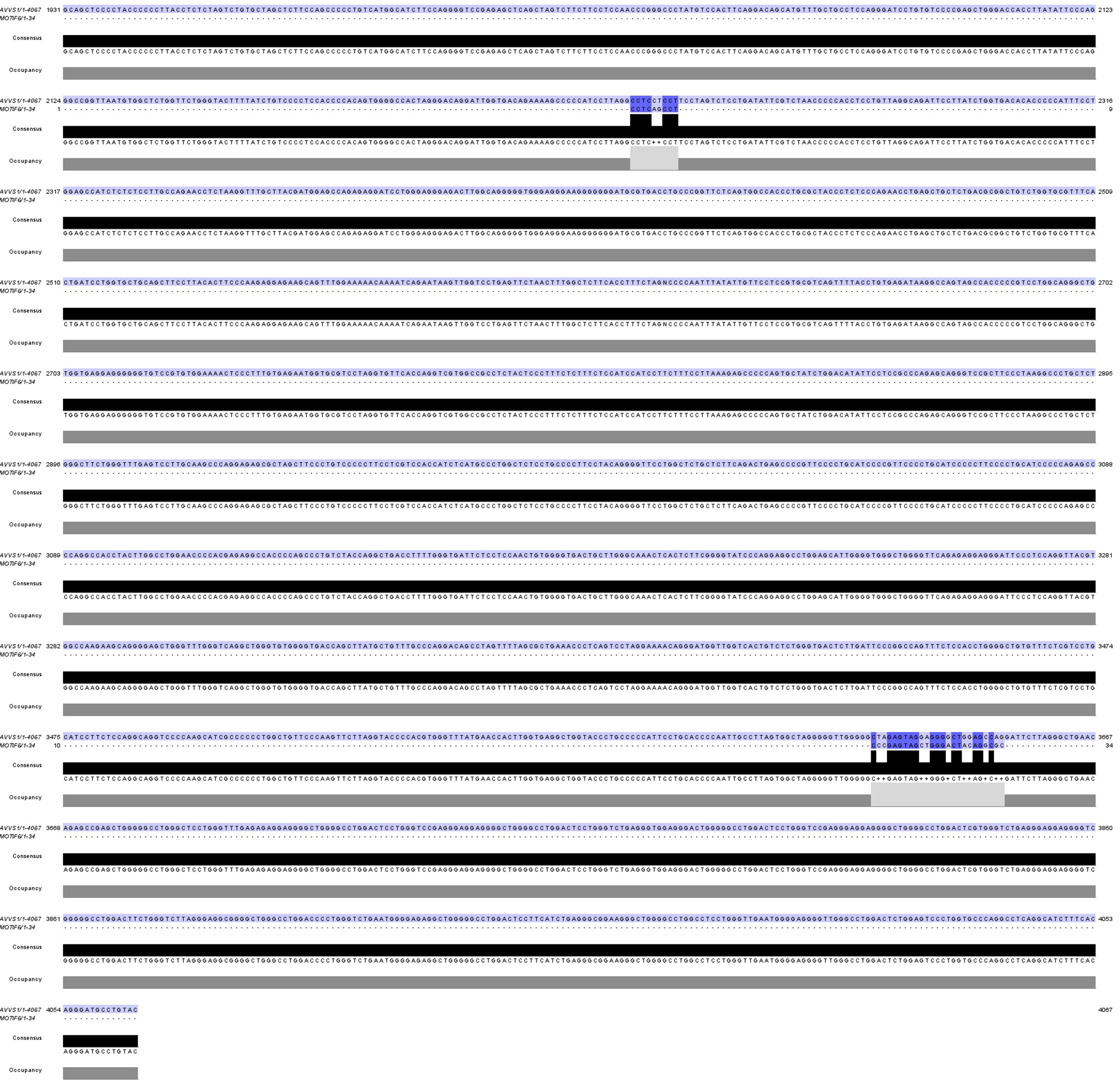

### Figure S25

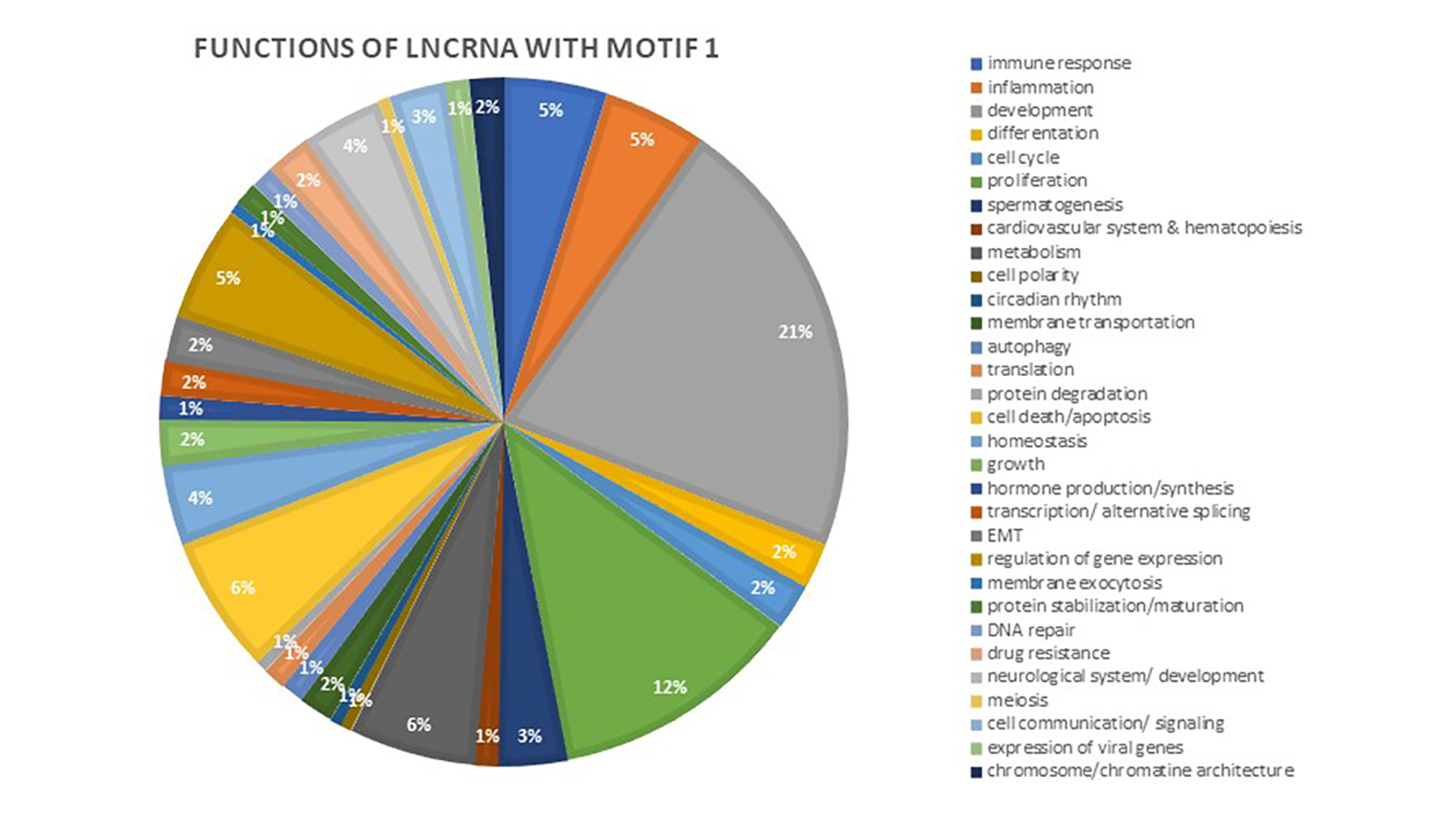

### Figure S26

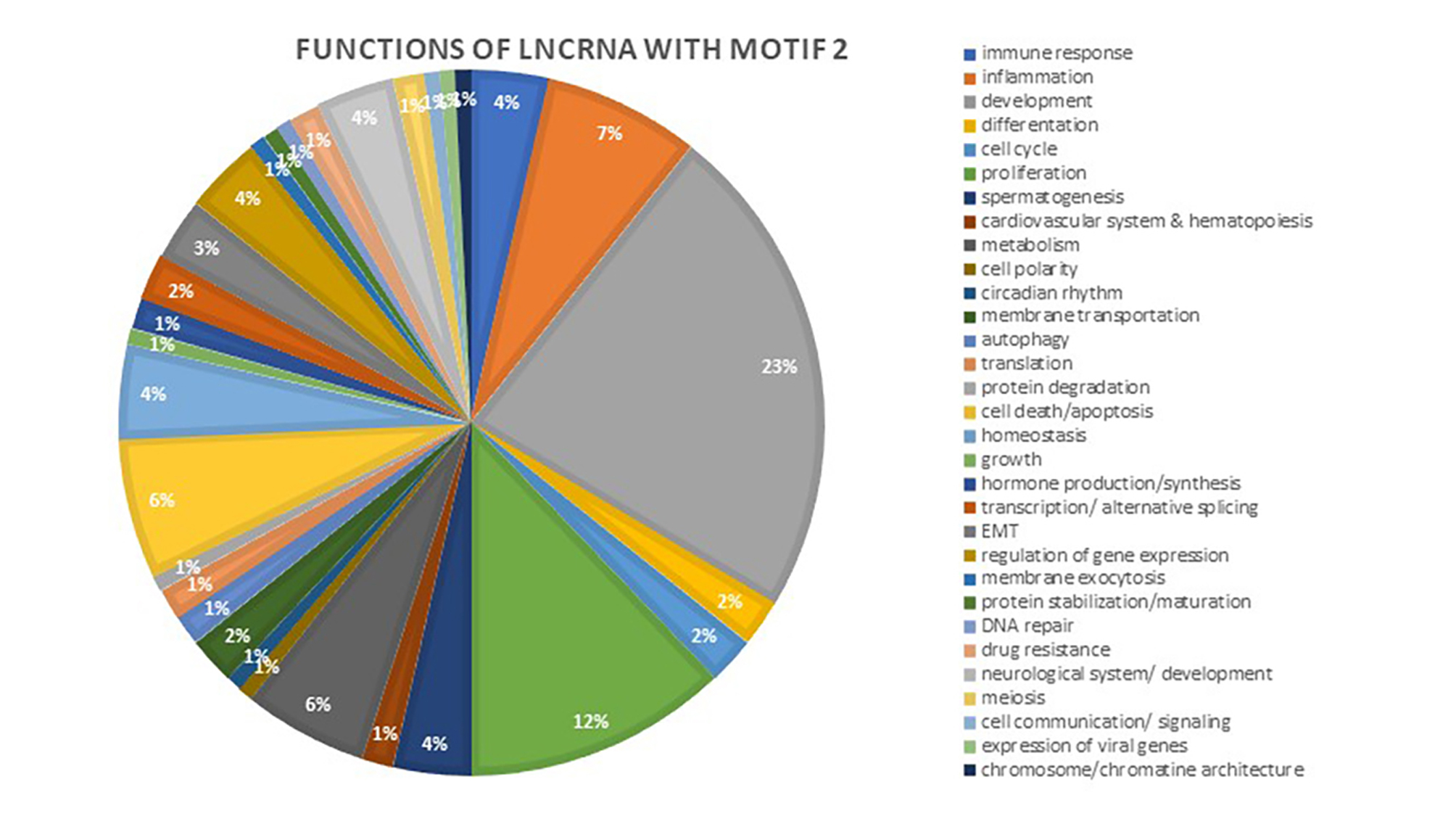

### Figure S27

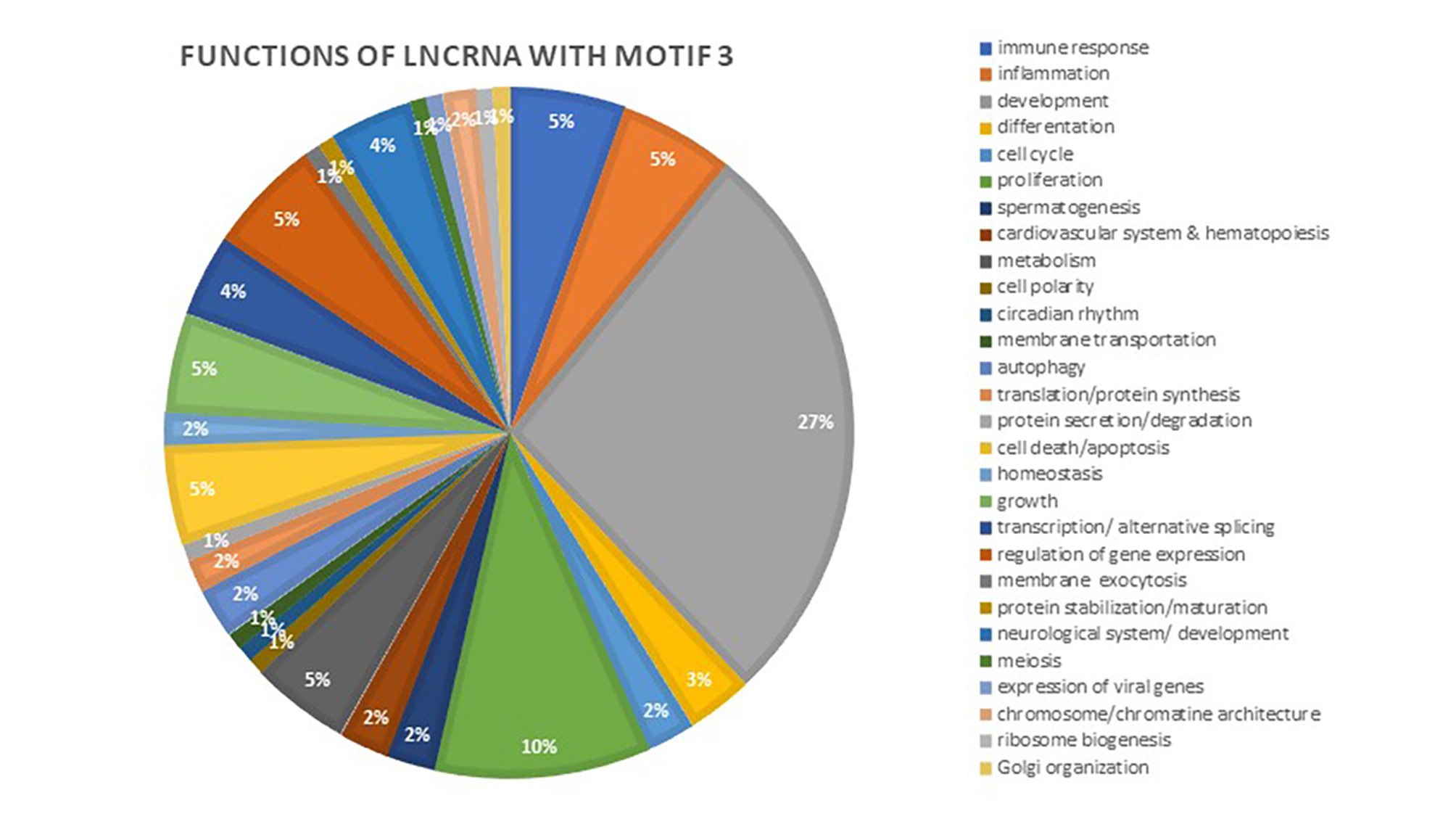

### Figure S28

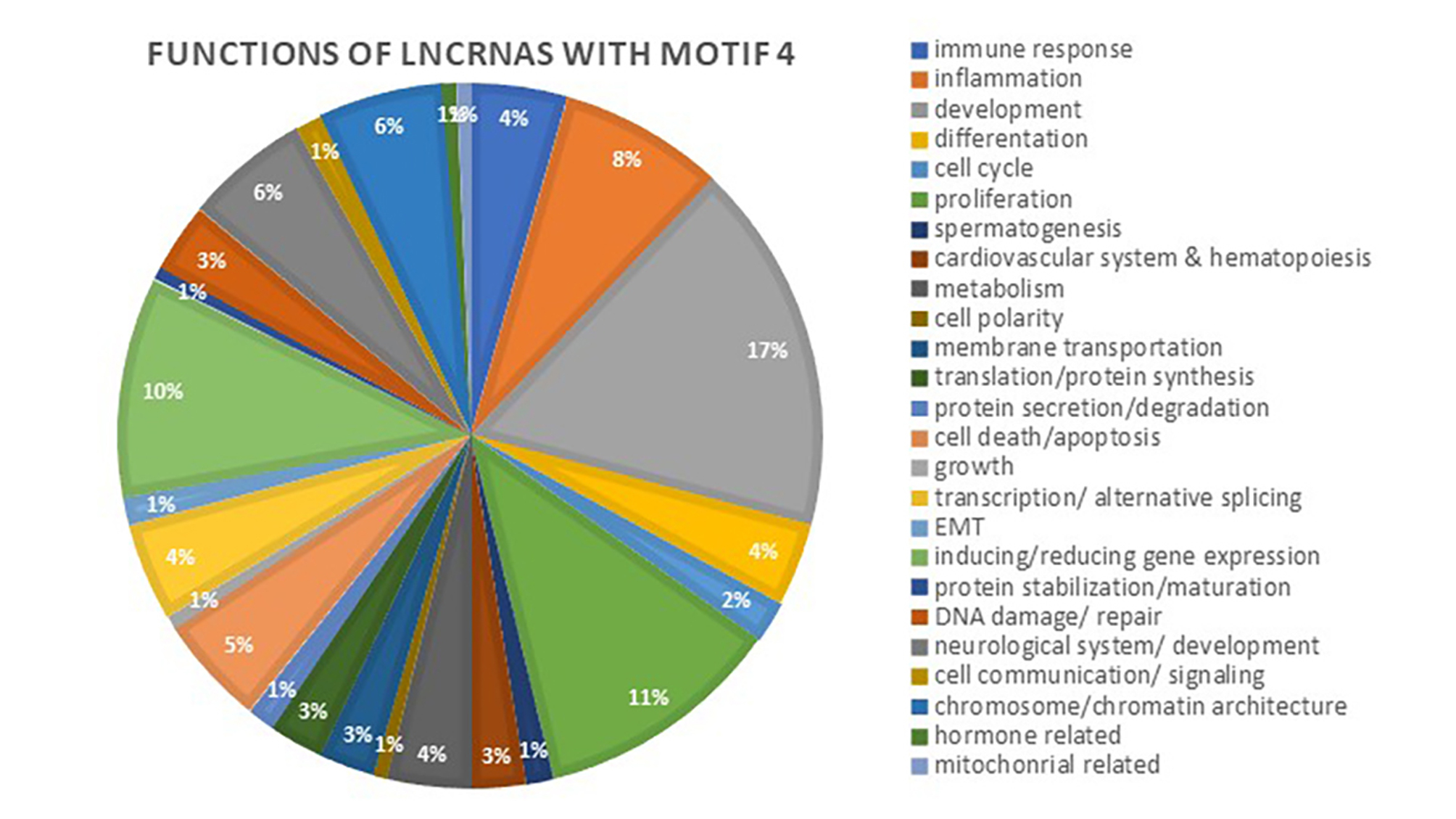

### Figure S29

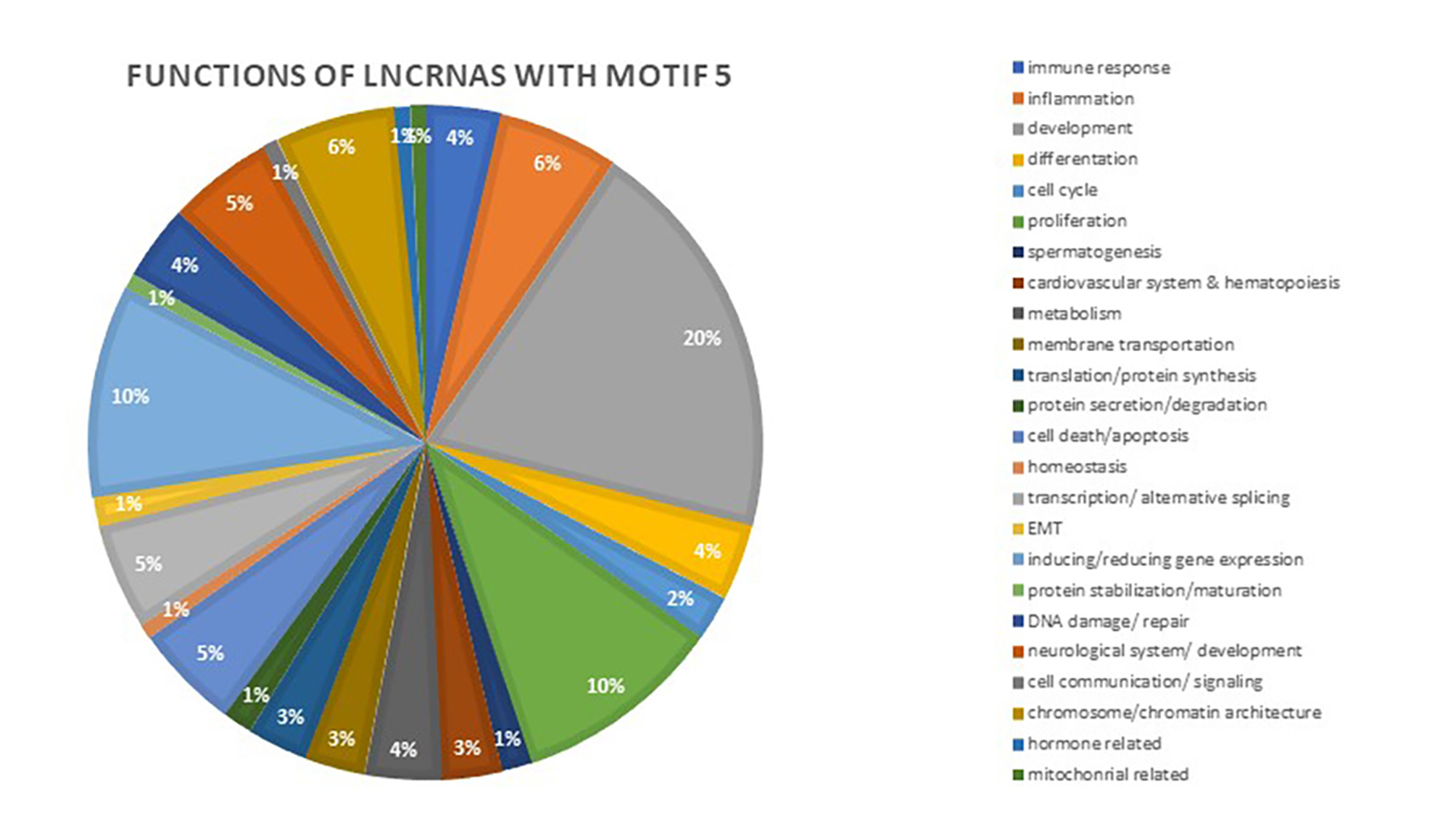

### Figure S30

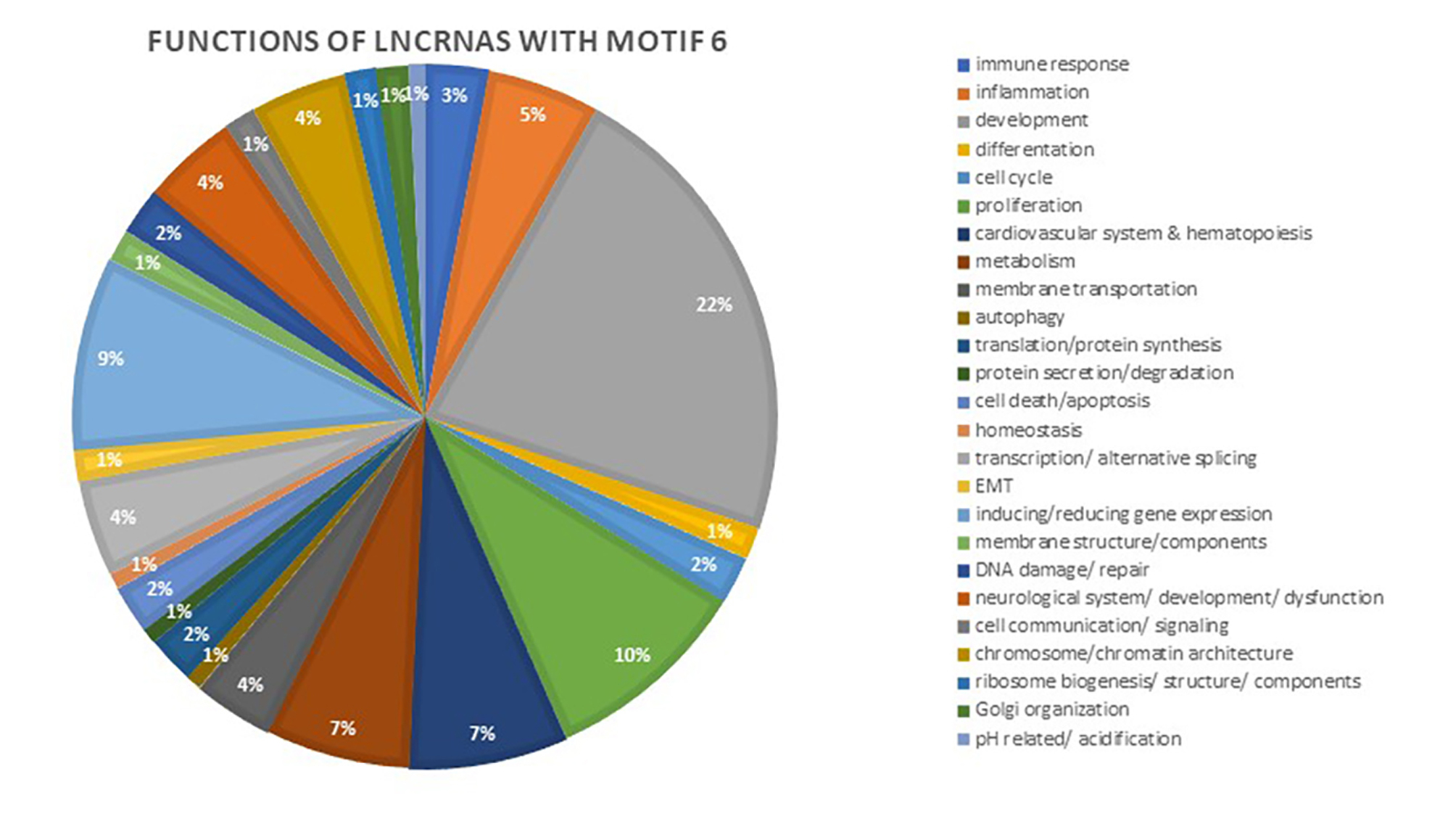
