## Supplementary material for "Role of lncRNAs related to NRs in the regulation of gene expression": Table S1

Table S. 1: The dataset of the lncRNAs that are either upregulated or downregulated after the activation of an NR.

| lncRNAs | Effect |
| --- | --- |
| HOTAIR | Upregulated |
| MALAT-1 | Downregulated |
| NEAT1 | Upregulated |
| TC0101441 | Upregulated |
| TC0101686 | Downregulated |
| FOXC1 eRNA | Upregulated |
| CA12 eRNA | Upregulated |
| SIAH2 eRNA | Upregulated |
| NRIP1 eRNA | Upregulated |
| GREB1 eRNA | Upregulated |
| PGR eRNA | Upregulated |
| SMAD7 eRNA | Upregulated |
| P2RY eRNA | Upregulated |
| TFF1 eRNA | Upregulated |
| KCNK5 eRNA | Upregulated |
| KLK3 eRNA | Upregulated |
| Cx3cr1 eRNA | Downregulated |
| Mmp9 eRNA | Downregulated |
| ADAMTS9-AS2 | Upregulated |
| COL18A1-AS1 | Upregulated |
| CPB2-AS1 | Upregulated |
| FRY-AS1 | Upregulated |
| GNG12-AS1 | Upregulated |
| MLLT4-AS1 | Upregulated |
| MYLK-AS1 | Upregulated |
| RERG-AS1 | Upregulated |
| TPRG1-AS1 | Upregulated |
| TTC28-AS1 | Upregulated |
| ZNF295-AS1 | Upregulated |
| ENOX1-AS1 | Downregulated |
| EXTL3-AS1 | Downregulated |
| GCFC1-AS1 | Downregulated |
| LIFR-AS1 | Downregulated |
| UBAC2-AS1 | Downregulated |
| BCYRN1 | Downregulated |
| DIO3OS | Upregulated |
| DLEU2 | Downregulated |
| FKBP1A-SDCBP2 | Upregulated |
| FTX | Downregulated |
| HAR1B | Upregulated |
| LOC147976 | Downregulated |
| MIR205HG | Downregulated |
| MIR7-3HG | Upregulated |
| MIRLET7DHG | Downregulated |
| NKAPP1 | Downregulated |
| PVT1 | Upregulated |
| H19 | Upregulated (E2;Cort.) |
|  | Downregulated (P4) |
|  | Downregulated |
| GAS6 (GenBank Accession Spotted EST : BF879099) | Downregulated |
| RAB25 (GenBank Accession Spotted EST : BF360792) | Downregulated |
| SPIRE2 (GenBank Accession Spotted EST : CK327123) | Downregulated |
| AW805354 | Downregulated |
| PHC2 (GenBank Accession Spotted EST : BF854186) | Downregulated |
| NIBP (GenBank Accession Spotted EST : BF333281) | Downregulated |
| DST (GenBank Accession Spotted EST : AW860958) | Downregulated |
| ACTN4 (GenBank Accession Spotted EST : BF768459) | Downregulated |
| DNAJC3 (GenBank Accession Spotted EST : BF368584) | Downregulated |
| ACTN4 (GenBank Accession Spotted EST : BF882783) | Downregulated |
| CFLAR (GenBank Accession Spotted EST : AW815632) | Upregulated |
| RSNL2 (GenBank Accession Spotted EST : CK327134) | Upregulated |
| ITGA6 (GenBank Accession Spotted EST : CK327184) | Upregulated |
| PALLD (GenBank Accession Spotted EST : BE156190) | Upregulated |
| SAP18 (GenBank Accession Spotted EST : BF332494) | Upregulated |
| STARD13 (GenBank Accession Spotted EST : BE087892) | Upregulated |
| TBN (GenBank Accession Spotted EST : BE061111) | Upregulated |
| ERG (GenBank Accession Spotted EST : AW805635) | Upregulated |
| NBPF9 (GenBank Accession Spotted EST :BG004239) | Upregulated |
| KDELR2 (GenBank Accession Spotted EST : BF350758) | Upregulated |
| PMF1 (GenBank Accession Spotted EST : BF805741) | Upregulated |
| PPP3CB (GenBank Accession Spotted EST : CK327191) | Upregulated |
| P2RX1 (GenBank Accession Spotted EST : BF894911) | Upregulated |
| TGFBR2 (GenBank Accession Spotted EST : AW880165) | Upregulated |
| AW805625 | Upregulated |
| ZNF644 (GenBank Accession Spotted EST :CK327190) | Upregulated |
| PASK (GenBank Accession Spotted EST :BE168353) | Upregulated |
| MYO5A (GenBank Accession Spotted EST :BF848956) | Upregulated |
| UBE2V1 (GenBank Accession Spotted EST :BE062809) | Upregulated |
| BF768642 | Upregulated |
| P2RY14 (GenBank Accession Spotted EST :CK327189) | Upregulated |
| PPP2R2A (GenBank Accession Spotted EST :AW819948) | Upregulated |
| PECAM1 (GenBank Accession Spotted EST :AW819948) | Upregulated |
| XRCC1 (GenBank Accession Spotted EST : BF926453) | Upregulated |
| ADD3 (GenBank Accession Spotted EST :BF926454) | Upregulated |
| ALMS1 (GenBank Accession Spotted EST :BF881469) | Upregulated |
| C16orf46 (GenBank Accession Spotted EST :BF364131) | Upregulated |
| ATF2 (GenBank Accession Spotted EST : CK327094) | Upregulated |
| MEG3 (GenBank Accession Spotted EST : BF364068) | Upregulated |
| PCA3 | Upregulated |
| PCAT18 | Upregulated |
| CTBP1-AS | Upregulated |
| PCGEM1 | Upregulated |
| PlncRNA-1 | Downregulated |
| Air (antisense Igf2r RNA) | Upregulated |
| AntiPeg11 | Upregulated |
| Dio3os | Upregulated |
| Dlx1as | Upregulated |
| mHOTAIR (Hox antisense intergenic RNA) | Upregulated |
| HOTTIP (HOXA transcript at the distal tip) | Upregulated |
| H19 | Upregulated |
| Nespas (Neuroendocrine secretory protein antisense) | Upregulated |
| Malat1 | Upregulated |
| Mistral | Upregulated |
| Msx1as | Upregulated |
| linc1242-LINC-Enah | Upregulated |
| PINC | Upregulated |
| Recombination hot spot RNA | Upregulated |
| SRA (steroid receptor RNA activator) | Upregulated |
| Zfas1 | Upregulated |
| BGn-As | Downregulated |
| Foxn2-as | Downregulated |
| Gtl2-as | Downregulated |
| H19-as | Downregulated |
| Kcnq1ot1 | Downregulated |
| LincRNA-p21 | Downregulated |
| Linc-MD1 | Downregulated |
| AC002398.5 | Upregulated |
| AC005029.1 | Upregulated |
| AC006076.1 | Upregulated |
| AC007365.1 | Upregulated |
| AC007365.3 | Upregulated |
| AC007405.4 | Upregulated |
| AC009276.4 | Upregulated |
| AC009495.3 | Upregulated |
| AC009495.4 | Upregulated |
| AC009531.2 | Upregulated |
| AC010096.2 | Upregulated |
| AC010733.4 | Upregulated |
| AC011747.3 | Upregulated |
| AC011747.4 | Upregulated |
| AC012370.3 | Upregulated |
| AC016912.3 | Upregulated |
| AC017074.2 | Upregulated |
| AC019117.1 | Upregulated |
| AC019118.2 | Upregulated |
| AC019118.3 | Upregulated |
| AC020571.3 | Upregulated |
| AC025171.1 | Upregulated |
| AC027119.1 | Upregulated |
| AC068483.1 | Upregulated |
| AC068858.1 | Upregulated |
| AC073046.25 | Upregulated |
| AC073283.4 | Upregulated |
| AC084018.1 | Upregulated |
| AC090421.1 | Upregulated |
| AC093063.1 | Upregulated |
| AC093375.1 | Upregulated |
| AC099552.3 | Upregulated |
| AC104655.3 | Upregulated |
| AC116609.2 | Upregulated |
| AC116609.3 | Upregulated |
| AC123768.4 | Upregulated |
| AC131097.3 | Upregulated |
| AC144450.2 | Upregulated |
| AC145110.1 | Upregulated |
| ADAM20P1 | Upregulated |
| AF064858.8 | Upregulated |
| AF241725.6 | Upregulated |
| AP001258.4 | Upregulated |
| AP001505.9 | Upregulated |
| BCYRN1 | Upregulated |
| C17orf69 | Upregulated |
| CTA-126B4.7 | Upregulated |
| CTA-150C2.13 | Upregulated |
| CTA-363E6.2 | Upregulated |
| CTA-392C11.2 | Upregulated |
| CTC-205M6.2 | Upregulated |
| CTC-228N24.3 | Upregulated |
| CTC-497E21.3 | Upregulated |
| CTC-497E21.4 | Upregulated |
| CTD-2139B15.2 | Upregulated |
| CTD-2193G5.1 | Upregulated |
| CTD-2201E18.3 | Upregulated |
| CTD-2210P24.6 | Upregulated |
| CTD-2215E18.3 | Upregulated |
| CTD-2333K2.1 | Upregulated |
| CTD-2552B11.4 | Upregulated |
| CTD-2561B21.11 | Upregulated |
| CTD-2568A17.1 | Upregulated |
| CTD-2650P22.1 | Upregulated |
| CTD-3032H12.2 | Upregulated |
| CTD-3035D6.2 | Upregulated |
| CTD-3051D23.1 | Upregulated |
| CTD-3051D23.3 | Upregulated |
| CTD-3074O7.2 | Upregulated |
| dblinc-ENST00000379093 | Upregulated |
| dblinc-ENST00000498984 | Upregulated |
| dblinc-ENST00000500478 | Upregulated |
| dblinc-ENST00000500836 | Upregulated |
| dblinc-ENST00000500941 | Upregulated |
| dblinc-ENST00000501118 | Upregulated |
| dblinc-ENST00000501254 | Upregulated |
| dblinc-ENST00000501265 | Upregulated |
| dblinc-ENST00000503487 | Upregulated |
| dblinc-ENST00000503629 | Upregulated |
| dblinc-ENST00000504230 | Upregulated |
| dblinc-ENST00000505156 | Upregulated |
| dblinc-ENST00000505360 | Upregulated |
| dblinc-ENST00000505651 | Upregulated |
| dblinc-ENST00000505880 | Upregulated |
| dblinc-ENST00000507426 | Upregulated |
| dblinc-ENST00000507930 | Upregulated |
| dblinc-ENST00000508109 | Upregulated |
| dblinc-ENST00000508332 | Upregulated |
| dblinc-ENST00000509213 | Upregulated |
| dblinc-ENST00000509401 | Upregulated |
| dblinc-ENST00000510288 | Upregulated |
| dblinc-ENST00000510752 | Upregulated |
| dblinc-ENST00000511239 | Upregulated |
| dblinc-ENST00000512200 | Upregulated |
| dblinc-ENST00000512341 | Upregulated |
| dblinc-ENST00000513068 | Upregulated |
| dblinc-ENST00000514792 | Upregulated |
| dblinc-ENST00000515416 | Upregulated |
| dblinc-ENST00000515449 | Upregulated |
| dblinc-NR_001278 | Upregulated |
| dblinc-NR_001590_ | Upregulated |
| dblinc-NR_001590__ | Upregulated |
| dblinc-NR_002799 | Upregulated |
| dblinc-NR_002801 | Upregulated |
| dblinc-NR_002809 | Upregulated |
| dblinc-NR_003263 | Upregulated |
| dblinc-NR_003683 | Upregulated |
| dblinc-NR_003716 | Upregulated |
| dblinc-NR_015360 | Upregulated |
| dblinc-NR_015407 | Upregulated |
| dblinc-NR_015424 | Upregulated |
| dblinc-NR_023382 | Upregulated |
| dblinc-NR_024037 | Upregulated |
| dblinc-NR_024087 | Upregulated |
| dblinc-NR_024103 | Upregulated |
| dblinc-NR_024256 | Upregulated |
| dblinc-NR_024349 | Upregulated |
| dblinc-NR_024355 | Upregulated |
| dblinc-NR_024418 | Upregulated |
| dblinc-NR_024436 | Upregulated |
| dblinc-NR_026681 | Upregulated |
| dblinc-NR_026730 | Upregulated |
| dblinc-NR_026755 | Upregulated |
| dblinc-NR_026762 | Upregulated |
| dblinc-NR_026765 | Upregulated |
| dblinc-NR_026807 | Upregulated |
| dblinc-NR_026864 | Upregulated |
| dblinc-NR_026979 | Upregulated |
| dblinc-NR_027072 | Upregulated |
| dblinc-NR_027131 | Upregulated |
| dblinc-NR_027183 | Upregulated |
| dblinc-NR_027358 | Upregulated |
| dblinc-NR_027706 | Upregulated |
| dblinc-NR_027994 | Upregulated |
| dblinc-NR_028301 | Upregulated |
| dblinc-NR_033920 | Upregulated |
| dblinc-NR_033962 | Upregulated |
| dblinc-NR_033980 | Upregulated |
| dblinc-NR_034023 | Upregulated |
| dblinc-NR_034034 | Upregulated |
| dblinc-NR_034105 | Upregulated |
| dblinc-NR_034130 | Upregulated |
| dblinc-NR_034132 | Upregulated |
| dblinc-NR_034156 | Upregulated |
| dblinc-NR_034158 | Upregulated |
| dblinc-NR_034179 | Upregulated |
| dblinc-NR_036530 | Upregulated |
| dblinc-uc001crj.1 | Upregulated |
| dblinc-uc001cvn.1 | Upregulated |
| dblinc-uc001dmk.2 | Upregulated |
| dblinc-uc001duc.2 | Upregulated |
| dblinc-uc001gsf.2 | Upregulated |
| dblinc-uc001hpx.2 | Upregulated |
| dblinc-uc001hwh.2 | Upregulated |
| dblinc-uc001hwi.1 | Upregulated |
| dblinc-uc001ikv.1 | Upregulated |
| dblinc-uc001jjs.2 | Upregulated |
| dblinc-uc001khy.1 | Upregulated |
| dblinc-uc001lkw.1 | Upregulated |
| dblinc-uc001pof.1 | Upregulated |
| dblinc-uc001uzl.2 | Upregulated |
| dblinc-uc001vca.2 | Upregulated |
| dblinc-uc001wun.2 | Upregulated |
| dblinc-uc001xev.1 | Upregulated |
| dblinc-uc001zjy.2 | Upregulated |
| dblinc-uc002obw.1 | Upregulated |
| dblinc-uc002pnr.1 | Upregulated |
| dblinc-uc002qvl.2 | Upregulated |
| dblinc-uc002qxh.1 | Upregulated |
| dblinc-uc002rvu.1 | Upregulated |
| dblinc-uc002rzk.2 | Upregulated |
| dblinc-uc002skg.2 | Upregulated |
| dblinc-uc002thh.1 | Upregulated |
| dblinc-uc002uaw.1 | Upregulated |
| dblinc-uc002wjr.2 | Upregulated |
| dblinc-uc002yjw.1 | Upregulated |
| dblinc-uc002yqj.1 | Upregulated |
| dblinc-uc003fux.1 | Upregulated |
| dblinc-uc003hdc.1 | Upregulated |
| dblinc-uc003igy.2 | Upregulated |
| dblinc-uc003jfz.1 | Upregulated |
| dblinc-uc003mzy.2 | Upregulated |
| dblinc-uc003nhj.2 | Upregulated |
| dblinc-uc003pas.1 | Upregulated |
| dblinc-uc003pmm.2 | Upregulated |
| dblinc-uc003prp.1 | Upregulated |
| dblinc-uc003qis.2 | Upregulated |
| dblinc-uc003qje.1 | Upregulated |
| dblinc-uc003tbe.1 | Upregulated |
| dblinc-uc003ugk.2 | Upregulated |
| dblinc-uc003wlv.1 | Upregulated |
| dblinc-uc003wlw.1 | Upregulated |
| dblinc-uc003xhp.2 | Upregulated |
| dblinc-uc004auk.2 | Upregulated |
| dblinc-uc004dry.1 | Upregulated |
| dblinc-uc010izt.1 | Upregulated |
| DLEU2 | Upregulated |
| ENOX1-AS1 | Upregulated |
| EXTL3-AS1 | Upregulated |
| FTX | Upregulated |
| GCFC1-AS1 | Upregulated |
| KB-1460A1.5 | Upregulated |
| LIFR-AS1 | Upregulated |
| LINC00152 | Upregulated |
| LINC00263 | Upregulated |
| LINC00362 | Upregulated |
| LINC00441 | Upregulated |
| LINC00472 | Upregulated |
| LINC00536 | Upregulated |
| LINC00558 | Upregulated |
| LINC00575 | Upregulated |
| LINC00589 | Upregulated |
| linc-ABCA1-1 | Upregulated |
| linc-ADCY2-9 | Upregulated |
| linc-AGMAT-3 | Upregulated |
| linc-AIRE-2 | Upregulated |
| linc-AK2-1 | Upregulated |
| linc-AKR1E2-10 | Upregulated |
| linc-ALOX12-1 | Upregulated |
| linc-ANGEL2-1 | Upregulated |
| linc-ANP32A-3 | Upregulated |
| linc-APBB2-3 | Upregulated |
| linc-APOBEC3A-1 | Upregulated |
| linc-APPL2-1 | Upregulated |
| linc-AQP11 | Upregulated |
| linc-ARHGEF38-2 | Upregulated |
| linc-ARL4D-1 | Upregulated |
| linc-ARRDC4-3 | Upregulated |
| linc-ASAP2-4 | Upregulated |
| linc-ASCL1-1 | Upregulated |
| linc-ATAD5-2 | Upregulated |
| linc-ATG5-1 | Upregulated |
| linc-ATIC-3 | Upregulated |
| linc-ATP10A-3 | Upregulated |
| linc-ATP6V1G3-1 | Upregulated |
| linc-B3GAT2-4 | Upregulated |
| linc-BAHD1 | Upregulated |
| linc-BAT2L1 | Upregulated |
| linc-BCL2A1-3 | Upregulated |
| linc-BET3L | Upregulated |
| linc-C10orf11-1 | Upregulated |
| linc-C10orf46 | Upregulated |
| linc-C10orf57-5 | Upregulated |
| linc-C11orf66 | Upregulated |
| linc-C13orf1-3 | Upregulated |
| linc-C13orf23-1 | Upregulated |
| linc-C14orf39 | Upregulated |
| linc-C1orf201-1 | Upregulated |
| linc-C1orf210 | Upregulated |
| linc-C1orf63 | Upregulated |
| linc-C1orf87-1 | Upregulated |
| linc-C20orf29 | Upregulated |
| linc-C5orf38-7 | Upregulated |
| linc-C5orf39 | Upregulated |
| linc-C6orf145-3 | Upregulated |
| linc-C7orf11 | Upregulated |
| linc-C7orf45-2 | Upregulated |
| linc-C8orf86-1 | Upregulated |
| linc-CACNB3-1 | Upregulated |
| linc-CALB2-2 | Upregulated |
| linc-CALD1 | Upregulated |
| linc-CALML5-2 | Upregulated |
| linc-CALML5-3 | Upregulated |
| linc-CCDC122-4 | Upregulated |
| linc-CCDC54 | Upregulated |
| linc-CCDC92-1 | Upregulated |
| linc-CCNI-2 | Upregulated |
| linc-CCNK-1 | Upregulated |
| linc-CCR8-3 | Upregulated |
| linc-CD180-13 | Upregulated |
| linc-CD180-4 | Upregulated |
| linc-CD247 | Upregulated |
| linc-CDH3-4 | Upregulated |
| linc-CELF2-7 | Upregulated |
| linc-CETN1-1 | Upregulated |
| linc-CHAC2-5 | Upregulated |
| linc-CHAD-2 | Upregulated |
| linc-CHCHD6-1 | Upregulated |
| linc-CHRD | Upregulated |
| linc-CILP | Upregulated |
| linc-CITED2-4 | Upregulated |
| linc-CLLU1-11 | Upregulated |
| linc-CMPK2-1 | Upregulated |
| linc-CNOT2 | Upregulated |
| linc-CNOT6-1 | Upregulated |
| linc-COL28A1 | Upregulated |
| linc-COL9A2-2 | Upregulated |
| linc-CTNND2-2 | Upregulated |
| linc-CYR61-1 | Upregulated |
| linc-DHRS4-2 | Upregulated |
| linc-DHRS7 | Upregulated |
| linc-DHX37-20 | Upregulated |
| linc-DKFZp761E198 | Upregulated |
| linc-DKK3-3 | Upregulated |
| linc-DMRTB1-2 | Upregulated |
| linc-DSP | Upregulated |
| linc-DTHD1-7 | Upregulated |
| linc-DTL-1 | Upregulated |
| linc-DUSP4-1 | Upregulated |
| linc-DUSP4-6 | Upregulated |
| linc-DUT-1 | Upregulated |
| linc-DUT-2 | Upregulated |
| linc-DYNLL2-2 | Upregulated |
| linc-E2F7-4 | Upregulated |
| linc-ECHDC3-1 | Upregulated |
| linc-EIF5 | Upregulated |
| linc-ELK3-1 | Upregulated |
| linc-EPHA7-5 | Upregulated |
| linc-ERG-11 | Upregulated |
| linc-FAAH2-3 | Upregulated |
| linc-FAF1-1 | Upregulated |
| linc-FAM101A-4 | Upregulated |
| linc-FAM156A-1 | Upregulated |
| linc-FAM174B | Upregulated |
| linc-FAM183A | Upregulated |
| linc-FAM186A-2 | Upregulated |
| linc-FAM187B-2 | Upregulated |
| linc-FAM75C2-2 | Upregulated |
| linc-FAM78A-2 | Upregulated |
| linc-FBXL14 | Upregulated |
| linc-FBXO16-2 | Upregulated |
| linc-FER-2 | Upregulated |
| linc-FOXA1-2 | Upregulated |
| linc-FOXP3 | Upregulated |
| linc-FRG1-2 | Upregulated |
| linc-FRG1-5 | Upregulated |
| linc-FSCB | Upregulated |
| linc-G3BP1 | Upregulated |
| linc-GARS-1 | Upregulated |
| linc-GATA3-1 | Upregulated |
| linc-GATA3-2 | Upregulated |
| linc-GATA5-4 | Upregulated |
| linc-GGPS1-4 | Upregulated |
| linc-GGTLC1-7 | Upregulated |
| linc-GIN1-3 | Upregulated |
| linc-GPATCH2-1 | Upregulated |
| linc-GPHN-3 | Upregulated |
| linc-GPR183-3 | Upregulated |
| linc-GPR45-1 | Upregulated |
| linc-GPR45-3 | Upregulated |
| linc-GREM2-1 | Upregulated |
| linc-GRHL2-5 | Upregulated |
| linc-GRIK3-2 | Upregulated |
| linc-GTF2F2-1 | Upregulated |
| linc-GTPBP8 | Upregulated |
| linc-HAS2-2 | Upregulated |
| linc-HIVEP2-1 | Upregulated |
| linc-HNRNPA1-2 | Upregulated |
| linc-HNRNPA2B1-1 | Upregulated |
| linc-HOXC13-3 | Upregulated |
| linc-HRH3-3 | Upregulated |
| linc-HS6ST1-2 | Upregulated |
| linc-ID2-1 | Upregulated |
| linc-IFT74-5 | Upregulated |
| linc-IGFBP2-3 | Upregulated |
| linc-IGHMBP2 | Upregulated |
| linc-IL1F7 | Upregulated |
| linc-IL2RG | Upregulated |
| linc-INSIG1-2 | Upregulated |
| linc-INSIG1-3 | Upregulated |
| linc-IQSEC2 | Upregulated |
| linc-IRX3-1 | Upregulated |
| linc-IRX3-5 | Upregulated |
| linc-ISX-3 | Upregulated |
| linc-ITIH2-3 | Upregulated |
| linc-JAM3-1 | Upregulated |
| linc-JAM3-2 | Upregulated |
| linc-JPH1-2 | Upregulated |
| linc-KARS-2 | Upregulated |
| linc-KCNV1-1 | Upregulated |
| linc-KIAA1024 | Upregulated |
| linc-KIAA1257 | Upregulated |
| linc-KIAA1429-1 | Upregulated |
| linc-KIF26B | Upregulated |
| linc-KIN-5 | Upregulated |
| linc-KLHL31-1 | Upregulated |
| linc-KRT7-1 | Upregulated |
| linc-KRTAP13-1-5 | Upregulated |
| linc-LAMA1-5 | Upregulated |
| linc-LCTL-1 | Upregulated |
| linc-LEMD3 | Upregulated |
| linc-LEPROTL1-3 | Upregulated |
| linc-LOC100129636-2 | Upregulated |
| linc-LPP-2 | Upregulated |
| linc-LRFN2-1 | Upregulated |
| linc-LRP12-1 | Upregulated |
| linc-LRP1B-2 | Upregulated |
| linc-LRP8-1 | Upregulated |
| linc-LRPPRC-1 | Upregulated |
| linc-LRRC41-2 | Upregulated |
| linc-LRRC49-2 | Upregulated |
| linc-LRRC49-3 | Upregulated |
| linc-LRRC49-4 | Upregulated |
| linc-LRRC58 | Upregulated |
| linc-LSMD1-1 | Upregulated |
| linc-LYZL1-4 | Upregulated |
| linc-MAN2A2 | Upregulated |
| linc-MAP3K5 | Upregulated |
| linc-MARCKS-3 | Upregulated |
| linc-MARK1-3 | Upregulated |
| linc-MBL2-2 | Upregulated |
| linc-MBOAT2-1 | Upregulated |
| linc-MCCC2 | Upregulated |
| linc-MECR-4 | Upregulated |
| linc-MEF2D | Upregulated |
| linc-MELK | Upregulated |
| linc-MFSD9-13 | Upregulated |
| linc-MGAT4C-1 | Upregulated |
| linc-MGAT5B-3 | Upregulated |
| linc-MMADHC-3 | Upregulated |
| linc-MMD-1 | Upregulated |
| linc-MTHFD2 | Upregulated |
| linc-MYT1L-1 | Upregulated |
| linc-NADSYN1-2 | Upregulated |
| linc-NANOS1-2 | Upregulated |
| linc-NAP1L2-1 | Upregulated |
| linc-NDUFA5-2 | Upregulated |
| linc-NDUFAF2-3 | Upregulated |
| linc-NID2-4 | Upregulated |
| linc-NLRP3-3 | Upregulated |
| linc-NUDT4-4 | Upregulated |
| linc-NUF2-1 | Upregulated |
| linc-ODF3B | Upregulated |
| linc-ODZ4-3 | Upregulated |
| linc-OLFML2A | Upregulated |
| linc-OPCML | Upregulated |
| linc-OR5AU1 | Upregulated |
| linc-P2RX5 | Upregulated |
| linc-P4HA1 | Upregulated |
| linc-PAM-5 | Upregulated |
| linc-PAPPA-4 | Upregulated |
| linc-PARP10-2 | Upregulated |
| linc-PDCD1-1 | Upregulated |
| linc-PDE6H-2 | Upregulated |
| linc-PEX13 | Upregulated |
| linc-PFDN4-3 | Upregulated |
| linc-PHACTR2-2 | Upregulated |
| linc-PHB-2 | Upregulated |
| linc-PICK1 | Upregulated |
| linc-PLTP | Upregulated |
| linc-POLR3G-1 | Upregulated |
| linc-PPA2-1 | Upregulated |
| linc-PPAP2A-2 | Upregulated |
| linc-PPP2R2C-1 | Upregulated |
| linc-PPT1-1 | Upregulated |
| linc-PRDM11-2 | Upregulated |
| linc-PRDM9-4 | Upregulated |
| linc-PSMD7-7 | Upregulated |
| linc-PTBP1 | Upregulated |
| linc-PTHLH-3 | Upregulated |
| linc-RAI2 | Upregulated |
| linc-RANBP3L-5 | Upregulated |
| linc-RASSF4 | Upregulated |
| linc-RC3H1-2 | Upregulated |
| linc-RCBTB1 | Upregulated |
| linc-REPIN1 | Upregulated |
| linc-RFPL4A | Upregulated |
| linc-RGS18-3 | Upregulated |
| linc-RHOB-5 | Upregulated |
| linc-RHOB-7 | Upregulated |
| linc-RHOXF1-4 | Upregulated |
| linc-RHOXF2 | Upregulated |
| linc-RILPL1-2 | Upregulated |
| linc-RIOK2-2 | Upregulated |
| linc-RIPK4 | Upregulated |
| linc-RPGRIP1L | Upregulated |
| linc-RPL38-3 | Upregulated |
| linc-RPRM-2 | Upregulated |
| linc-RYBP-3 | Upregulated |
| linc-SALL4 | Upregulated |
| linc-SDCCAG1-1 | Upregulated |
| linc-SEC24D | Upregulated |
| linc-SEP15-3 | Upregulated |
| linc-SERHL2-1 | Upregulated |
| linc-SGCG-1 | Upregulated |
| linc-SGCG-2 | Upregulated |
| linc-SH3D19 | Upregulated |
| linc-SH3GL3 | Upregulated |
| linc-SIP1-1 | Upregulated |
| linc-SLAMF9 | Upregulated |
| linc-SLC1A4-2 | Upregulated |
| linc-SLC25A45-4 | Upregulated |
| linc-SLC35F3 | Upregulated |
| linc-SLITRK5-16 | Upregulated |
| linc-SMC1B-4 | Upregulated |
| linc-SMG1 | Upregulated |
| linc-SNAP25-1 | Upregulated |
| linc-SOD3-1 | Upregulated |
| linc-SORCS1-2 | Upregulated |
| linc-SOX10 | Upregulated |
| linc-SOX6 | Upregulated |
| linc-SPAM1 | Upregulated |
| linc-SPANXB1-1 | Upregulated |
| linc-SPO11-1 | Upregulated |
| linc-SPRY1-1 | Upregulated |
| linc-SPRY1-2 | Upregulated |
| linc-SSTR1-4 | Upregulated |
| linc-ST8SIA3-3 | Upregulated |
| linc-STK32B-1 | Upregulated |
| linc-SUZ12-3 | Upregulated |
| linc-SV2C-2 | Upregulated |
| linc-SYNC-2 | Upregulated |
| linc-SYNJ2BP | Upregulated |
| linc-TACC2-1 | Upregulated |
| linc-TBC1D20 | Upregulated |
| linc-TBX20-1 | Upregulated |
| linc-TCEAL1 | Upregulated |
| linc-TET3 | Upregulated |
| linc-TEX28 | Upregulated |
| linc-TFB2M | Upregulated |
| linc-THBS1-3 | Upregulated |
| linc-THPO-2 | Upregulated |
| linc-THSD7B-4 | Upregulated |
| linc-THUMPD3 | Upregulated |
| linc-TLE3-2 | Upregulated |
| linc-TM4SF1 | Upregulated |
| linc-TM4SF4-1 | Upregulated |
| linc-TM6SF1 | Upregulated |
| linc-TMC3-1 | Upregulated |
| linc-TMCC1-2 | Upregulated |
| linc-TMEM150C | Upregulated |
| linc-TMEM161B-1 | Upregulated |
| linc-TMEM18-13 | Upregulated |
| linc-TMEM18-5 | Upregulated |
| linc-TMEM18-7 | Upregulated |
| linc-TMEM18-8 | Upregulated |
| linc-TMEM63B-2 | Upregulated |
| linc-TMEM74 | Upregulated |
| linc-TMPRSS15-18 | Upregulated |
| linc-TNPO3-3 | Upregulated |
| linc-TNR-2 | Upregulated |
| linc-TOX3-3 | Upregulated |
| linc-TPT1-1 | Upregulated |
| linc-TPT1-2 | Upregulated |
| linc-TRAPPC1 | Upregulated |
| linc-TRIM36 | Upregulated |
| linc-TRIML2-5 | Upregulated |
| linc-TRPS1-1 | Upregulated |
| linc-TSHZ2-4 | Upregulated |
| linc-TTLL7-3 | Upregulated |
| linc-TXNRD1 | Upregulated |
| linc-U2AF1-4 | Upregulated |
| linc-UBR2-1 | Upregulated |
| linc-UBR2-2 | Upregulated |
| linc-UGDH | Upregulated |
| linc-USP25-3 | Upregulated |
| linc-USP47-1 | Upregulated |
| linc-USPL1-1 | Upregulated |
| linc-USPL1-2 | Upregulated |
| linc-VLDLR-2 | Upregulated |
| linc-VPS36-1 | Upregulated |
| linc-VSIG10 | Upregulated |
| linc-VTA1-2 | Upregulated |
| linc-WDR60-2 | Upregulated |
| linc-WNT8B | Upregulated |
| linc-XIRP2-1 | Upregulated |
| linc-YOD1-2 | Upregulated |
| linc-ZBTB44 | Upregulated |
| linc-ZDHHC11 | Upregulated |
| linc-ZFC3H1-2 | Upregulated |
| linc-ZNF131-1 | Upregulated |
| linc-ZNF131-3 | Upregulated |
| linc-ZNF131-4 | Upregulated |
| linc-ZNF157 | Upregulated |
| linc-ZNF167 | Upregulated |
| linc-ZNF217 | Upregulated |
| linc-ZNF516-4 | Upregulated |
| linc-ZP4-1 | Upregulated |
| LOC147976 | Upregulated |
| MIR205HG | Upregulated |
| MIRLET7DHG | Upregulated |
| NKAPP1 | Upregulated |
| RP11-1000B6.5 | Upregulated |
| RP11-1049H7.2 | Upregulated |
| RP11-1084I9.2 | Upregulated |
| RP11-108K3.1 | Upregulated |
| RP11-1149M10.1 | Upregulated |
| RP11-114G22.1 | Upregulated |
| RP11-114M1.2 | Upregulated |
| RP11-117D22.2 | Upregulated |
| RP11-121C2.2 | Upregulated |
| RP11-123O10.4 | Upregulated |
| RP11-124N19.3 | Upregulated |
| RP11-135J2.3 | Upregulated |
| RP11-138J23.1 | Upregulated |
| RP11-172E10.1 | Upregulated |
| RP11-175B12.2 | Upregulated |
| RP11-183E9.3 | Upregulated |
| RP11-196G11.2 | Upregulated |
| RP11-207N4.3 | Upregulated |
| RP11-211C9.1 | Upregulated |
| RP11-211G23.2 | Upregulated |
| RP11-217B7.3 | Upregulated |
| RP11-221N13.3 | Upregulated |
| RP11-231E19.1 | Upregulated |
| RP11-236P24.3 | Upregulated |
| RP11-240A16.1 | Upregulated |
| RP11-247L20.3 | Upregulated |
| RP11-248E9.5 | Upregulated |
| RP11-257I8.2 | Upregulated |
| RP11-258C19.5 | Upregulated |
| RP11-267L5.1 | Upregulated |
| RP11-268I9.3 | Upregulated |
| RP11-269C23.3 | Upregulated |
| RP11-270M14.5 | Upregulated |
| RP11-278L15.2 | Upregulated |
| RP11-279F6.1 | Upregulated |
| RP11-279F6.2 | Upregulated |
| RP11-279F6.3 | Upregulated |
| RP11-279N8.3 | Upregulated |
| RP11-281P23.2 | Upregulated |
| RP11-297L17.3 | Upregulated |
| RP11-298A8.2 | Upregulated |
| RP11-2C15.1 | Upregulated |
| RP11-302J23.1 | Upregulated |
| RP11-306I1.2 | Upregulated |
| RP11-308B16.2 | Upregulated |
| RP11-30O15.1 | Upregulated |
| RP11-313F23.3 | Upregulated |
| RP11-318C24.2 | Upregulated |
| RP11-320N21.1 | Upregulated |
| RP11-321C24.1 | Upregulated |
| RP11-324E6.6 | Upregulated |
| RP11-332J15.3 | Upregulated |
| RP11-333O1.1 | Upregulated |
| RP11-344A16.1 | Upregulated |
| RP11-346C20.2 | Upregulated |
| RP11-346D14.1 | Upregulated |
| RP11-351E7.1 | Upregulated |
| RP11-359M6.1 | Upregulated |
| RP11-359N5.1 | Upregulated |
| RP11-363G2.5 | Upregulated |
| RP11-363J20.1 | Upregulated |
| RP11-371I1.2 | Upregulated |
| RP11-379F12.3 | Upregulated |
| RP11-379F12.4 | Upregulated |
| RP11-379F12.7 | Upregulated |
| RP11-37N22.1 | Upregulated |
| RP11-385J1.2 | Upregulated |
| RP11-38G5.2 | Upregulated |
| RP11-398E10.1 | Upregulated |
| RP11-415D17.4 | Upregulated |
| RP11-416N4.4 | Upregulated |
| RP11-428C6.2 | Upregulated |
| RP11-439E19.9 | Upregulated |
| RP11-439H13.2 | Upregulated |
| RP11-439L18.2 | Upregulated |
| RP11-446H18.3 | Upregulated |
| RP11-448P19.1 | Upregulated |
| RP11-450K4.1 | Upregulated |
| RP11-452L6.1 | Upregulated |
| RP11-454K7.3 | Upregulated |
| RP11-46A10.4 | Upregulated |
| RP11-48B3.3 | Upregulated |
| RP11-493L12.4 | Upregulated |
| RP11-496D24.2 | Upregulated |
| RP11-497D6.4 | Upregulated |
| RP11-49C24.1 | Upregulated |
| RP11-506M13.3 | Upregulated |
| RP11-511B23.1 | Upregulated |
| RP11-515E23.2 | Upregulated |
| RP11-530C5.1 | Upregulated |
| RP11-543F8.2 | Upregulated |
| RP11-55K22.5 | Upregulated |
| RP11-566K8.2 | Upregulated |
| RP11-571E6.3 | Upregulated |
| RP11-571O6.1 | Upregulated |
| RP11-579D7.4 | Upregulated |
| RP11-57C13.4 | Upregulated |
| RP11-584P21.2 | Upregulated |
| RP11-58E21.4 | Upregulated |
| RP11-610P16.1 | Upregulated |
| RP11-612B6.1 | Upregulated |
| RP11-612B6.2 | Upregulated |
| RP11-638I2.6 | Upregulated |
| RP11-646E18.4 | Upregulated |
| RP11-655M14.13 | Upregulated |
| RP11-65D17.1 | Upregulated |
| RP11-675F6.3 | Upregulated |
| RP11-680C21.1 | Upregulated |
| RP11-686G23.2 | Upregulated |
| RP11-6B19.1 | Upregulated |
| RP11-702F3.1 | Upregulated |
| RP11-711M9.1 | Upregulated |
| RP11-713M6.2 | Upregulated |
| RP11-713P17.3 | Upregulated |
| RP11-713P17.4 | Upregulated |
| RP11-713P17.5 | Upregulated |
| RP11-715J22.4 | Upregulated |
| RP11-716O23.1 | Upregulated |
| RP11-718G2.4 | Upregulated |
| RP11-726O12.3 | Upregulated |
| RP11-757G1.6 | Upregulated |
| RP11-766N7.3 | Upregulated |
| RP11-776H12.1 | Upregulated |
| RP11-77M5.1 | Upregulated |
| RP11-790J24.1 | Upregulated |
| RP11-79N23.1 | Upregulated |
| RP1-180E22.3 | Upregulated |
| RP11-818F20.4 | Upregulated |
| RP11-838N2.5 | Upregulated |
| RP11-849N15.2 | Upregulated |
| RP11-874J12.1 | Upregulated |
| RP11-888D10.3 | Upregulated |
| RP11-946L16.1 | Upregulated |
| RP11-946L20.2 | Upregulated |
| RP11-94C24.6 | Upregulated |
| RP11-963H4.5 | Upregulated |
| RP11-9G1.3 | Upregulated |
| RP1-209A6.1 | Upregulated |
| RP1-228P16.3 | Upregulated |
| RP1-238O23.4 | Upregulated |
| RP1-265C24.8 | Upregulated |
| RP3-410B11.1 | Upregulated |
| RP3-437I16.1 | Upregulated |
| RP3-461P17.10 | Upregulated |
| RP4-544H6.2 | Upregulated |
| RP4-601P9.2 | Upregulated |
| RP4-655J12.4 | Upregulated |
| RP4-660H19.1 | Upregulated |
| RP4-680D5.2 | Upregulated |
| RP4-724E16.2 | Upregulated |
| RP4-777D9.2 | Upregulated |
| RP4-777O23.1 | Upregulated |
| RP4-777O23.2 | Upregulated |
| RP4-781K5.4 | Upregulated |
| RP4-784A16.5 | Upregulated |
| RP4-813D12.3 | Upregulated |
| RP5-1086K13.1 | Upregulated |
| RP5-1109J22.2 | Upregulated |
| RP5-1112F19.2 | Upregulated |
| RP5-894A10.5 | Upregulated |
| RP5-899B16.1 | Upregulated |
| RP5-902P8.12 | Upregulated |
| RP5-907D15.3 | Upregulated |
| RP5-916L7.1 | Upregulated |
| RP5-916L7.2 | Upregulated |
| RP5-995J12.2 | Upregulated |
| RP6-109B7.2 | Upregulated |
| RP6-201G10.2 | Upregulated |
| UBAC2-AS1 | Upregulated |
| WI2-85898F10.1 | Upregulated |
| dblinc-uc004cjw.2 | Upregulated |
| linc-CEBPG | Upregulated |
| AC113607.2 | Downregulated |
| CTA-397C4.2 | Downregulated |
| CTB-164N12.1 | Downregulated |
| AC004383.4 | Downregulated |
| AC004383.5 | Downregulated |
| AC004448.5 | Downregulated |
| AC004840.9 | Downregulated |
| AC004863.6 | Downregulated |
| AC004947.2 | Downregulated |
| AC005013.5 | Downregulated |
| AC005355.2 | Downregulated |
| AC006028.9 | Downregulated |
| AC006445.8 | Downregulated |
| AC007182.6 | Downregulated |
| AC007255.8 | Downregulated |
| AC007386.2 | Downregulated |
| AC007464.1 | Downregulated |
| AC008940.1 | Downregulated |
| AC010132.10 | Downregulated |
| AC011648.1 | Downregulated |
| AC012065.4 | Downregulated |
| AC012531.25 | Downregulated |
| AC017002.4 | Downregulated |
| AC017060.1 | Downregulated |
| AC017076.4 | Downregulated |
| AC019349.2 | Downregulated |
| AC022182.3 | Downregulated |
| AC046143.7 | Downregulated |
| AC051649.6 | Downregulated |
| AC053503.4 | Downregulated |
| AC058791.1 | Downregulated |
| AC062028.1 | Downregulated |
| AC073636.1 | Downregulated |
| AC074389.7 | Downregulated |
| AC079776.3 | Downregulated |
| AC083864.3 | Downregulated |
| AC084082.3 | Downregulated |
| AC087590.3 | Downregulated |
| AC087859.1 | Downregulated |
| AC092296.1 | Downregulated |
| AC092635.1 | Downregulated |
| AC092687.3 | Downregulated |
| AC093326.3 | Downregulated |
| AC093382.1 | Downregulated |
| AC098828.2 | Downregulated |
| AC104135.2 | Downregulated |
| AC104135.3 | Downregulated |
| AC105053.4 | Downregulated |
| AC105760.2 | Downregulated |
| AC108058.1 | Downregulated |
| AC124861.1 | Downregulated |
| AC124944.2 | Downregulated |
| AC124944.5 | Downregulated |
| AC139666.1 | Downregulated |
| AC147651.1 | Downregulated |
| AC195454.1 | Downregulated |
| ADAMTS9-AS2 | Downregulated |
| AF127577.12 | Downregulated |
| AJ003147.8 | Downregulated |
| AJ009632.3 | Downregulated |
| AL122127.25 | Downregulated |
| AL161668.12 | Downregulated |
| AP000251.2 | Downregulated |
| AP000439.1 | Downregulated |
| AP000442.4 | Downregulated |
| AP001044.2 | Downregulated |
| AP001048.4 | Downregulated |
| AP001055.6 | Downregulated |
| AP001056.1 | Downregulated |
| AP002856.4 | Downregulated |
| BX322557.13 | Downregulated |
| BX571672.1 | Downregulated |
| C9orf106 | Downregulated |
| COL18A1-AS1 | Downregulated |
| CPB2-AS1 | Downregulated |
| CTA-14H9.5 | Downregulated |
| CTB-118N6.2 | Downregulated |
| CTB-17P3.4 | Downregulated |
| CTB-35F21.1 | Downregulated |
| CTC-276P9.1 | Downregulated |
| CTC-276P9.2 | Downregulated |
| CTC-293G12.1 | Downregulated |
| CTC-448D22.1 | Downregulated |
| CTC-542B22.1 | Downregulated |
| CTD-2005H7.1 | Downregulated |
| CTD-2089N3.2 | Downregulated |
| CTD-2187J20.1 | Downregulated |
| CTD-2515H24.3 | Downregulated |
| CTD-2515H24.4 | Downregulated |
| CTD-2555A7.2 | Downregulated |
| CTD-2588J6.2 | Downregulated |
| CTD-3023L14.1 | Downregulated |
| CTD-3023L14.2 | Downregulated |
| CTD-3224K15.2 | Downregulated |
| dblinc-ENST00000420523 | Downregulated |
| dblinc-ENST00000499000 | Downregulated |
| dblinc-ENST00000499525 | Downregulated |
| dblinc-ENST00000500076 | Downregulated |
| dblinc-ENST00000500113 | Downregulated |
| dblinc-ENST00000500181 | Downregulated |
| dblinc-ENST00000500381 | Downregulated |
| dblinc-ENST00000500975 | Downregulated |
| dblinc-ENST00000501288 | Downregulated |
| dblinc-ENST00000501387 | Downregulated |
| dblinc-ENST00000501730 | Downregulated |
| dblinc-ENST00000501751 | Downregulated |
| dblinc-ENST00000502085 | Downregulated |
| dblinc-ENST00000502101 | Downregulated |
| dblinc-ENST00000502737 | Downregulated |
| dblinc-ENST00000503956 | Downregulated |
| dblinc-ENST00000504269 | Downregulated |
| dblinc-ENST00000505771 | Downregulated |
| dblinc-ENST00000506222 | Downregulated |
| dblinc-ENST00000507040 | Downregulated |
| dblinc-ENST00000509267 | Downregulated |
| dblinc-ENST00000509303 | Downregulated |
| dblinc-ENST00000509942 | Downregulated |
| dblinc-ENST00000511013 | Downregulated |
| dblinc-ENST00000511655 | Downregulated |
| dblinc-ENST00000511756 | Downregulated |
| dblinc-ENST00000512879 | Downregulated |
| dblinc-ENST00000512967 | Downregulated |
| dblinc-ENST00000513207 | Downregulated |
| dblinc-ENST00000513305 | Downregulated |
| dblinc-ENST00000515029 | Downregulated |
| dblinc-ENST00000515276 | Downregulated |
| dblinc-NR_001298 | Downregulated |
| dblinc-NR_001447 | Downregulated |
| dblinc-NR_001459 | Downregulated |
| dblinc-NR_002710 | Downregulated |
| dblinc-NR_003245 | Downregulated |
| dblinc-NR_003574 | Downregulated |
| dblinc-NR_015419 | Downregulated |
| dblinc-NR_015450 | Downregulated |
| dblinc-NR_023938 | Downregulated |
| dblinc-NR_024065 | Downregulated |
| dblinc-NR_024100 | Downregulated |
| dblinc-NR_024187 | Downregulated |
| dblinc-NR_024253 | Downregulated |
| dblinc-NR_024280 | Downregulated |
| dblinc-NR_024351 | Downregulated |
| dblinc-NR_024362 | Downregulated |
| dblinc-NR_024394 | Downregulated |
| dblinc-NR_024431 | Downregulated |
| dblinc-NR_024480 | Downregulated |
| dblinc-NR_024493 | Downregulated |
| dblinc-NR_024585 | Downregulated |
| dblinc-NR_026655 | Downregulated |
| dblinc-NR_026658 | Downregulated |
| dblinc-NR_026764 | Downregulated |
| dblinc-NR_026845 | Downregulated |
| dblinc-NR_026852 | Downregulated |
| dblinc-NR_026863 | Downregulated |
| dblinc-NR_026951 | Downregulated |
| dblinc-NR_027026 | Downregulated |
| dblinc-NR_027058 | Downregulated |
| dblinc-NR_027108 | Downregulated |
| dblinc-NR_027148 | Downregulated |
| dblinc-NR_027241 | Downregulated |
| dblinc-NR_027341 | Downregulated |
| dblinc-NR_027345 | Downregulated |
| dblinc-NR_027441 | Downregulated |
| dblinc-NR_027451 | Downregulated |
| dblinc-NR_027793 | Downregulated |
| dblinc-NR_027906 | Downregulated |
| dblinc-NR_027906_ | Downregulated |
| dblinc-NR_028594 | Downregulated |
| dblinc-NR_033374 | Downregulated |
| dblinc-NR_033380 | Downregulated |
| dblinc-NR_033733 | Downregulated |
| dblinc-NR_033848 | Downregulated |
| dblinc-NR_033851 | Downregulated |
| dblinc-NR_033876 | Downregulated |
| dblinc-NR_033917 | Downregulated |
| dblinc-NR_033944 | Downregulated |
| dblinc-NR_033984 | Downregulated |
| dblinc-NR_034003 | Downregulated |
| dblinc-NR_034088 | Downregulated |
| dblinc-NR_034120 | Downregulated |
| dblinc-NR_036488 | Downregulated |
| dblinc-NR_036498 | Downregulated |
| dblinc-NR_036575 | Downregulated |
| dblinc-NR_036677 | Downregulated |
| dblinc-NR_036678 | Downregulated |
| dblinc-NR_036679 | Downregulated |
| dblinc-NR_037191 | Downregulated |
| dblinc-uc001acl.1 | Downregulated |
| dblinc-uc001ayw.2 | Downregulated |
| dblinc-uc001chr.2 | Downregulated |
| dblinc-uc001clo.1 | Downregulated |
| dblinc-uc001dbw.2 | Downregulated |
| dblinc-uc001dnt.1 | Downregulated |
| dblinc-uc001gqe.1 | Downregulated |
| dblinc-uc001hje.1 | Downregulated |
| dblinc-uc001hwk.2 | Downregulated |
| dblinc-uc001mmu.1 | Downregulated |
| dblinc-uc001ooz.1 | Downregulated |
| dblinc-uc001pcd.2 | Downregulated |
| dblinc-uc001qfd.1 | Downregulated |
| dblinc-uc001stq.1 | Downregulated |
| dblinc-uc001ugx.1 | Downregulated |
| dblinc-uc001wxl.2 | Downregulated |
| dblinc-uc001xov.1 | Downregulated |
| dblinc-uc001xrp.2 | Downregulated |
| dblinc-uc001xta.1 | Downregulated |
| dblinc-uc001zlf.1 | Downregulated |
| dblinc-uc002huv.1 | Downregulated |
| dblinc-uc002igh.2 | Downregulated |
| dblinc-uc002jdy.1 | Downregulated |
| dblinc-uc002kol.1 | Downregulated |
| dblinc-uc002krz.2 | Downregulated |
| dblinc-uc002nqn.1 | Downregulated |
| dblinc-uc002nsc.1 | Downregulated |
| dblinc-uc002nxe.2 | Downregulated |
| dblinc-uc002odw.1 | Downregulated |
| dblinc-uc002rae.1 | Downregulated |
| dblinc-uc002raz.1 | Downregulated |
| dblinc-uc002sqr.2 | Downregulated |
| dblinc-uc002vnc.2 | Downregulated |
| dblinc-uc002woe.1 | Downregulated |
| dblinc-uc003dde.2 | Downregulated |
| dblinc-uc003eir.2 | Downregulated |
| dblinc-uc003fax.1 | Downregulated |
| dblinc-uc003fto.1 | Downregulated |
| dblinc-uc003hrl.1 | Downregulated |
| dblinc-uc003hro.2 | Downregulated |
| dblinc-uc003hsn.1 | Downregulated |
| dblinc-uc003ito.1 | Downregulated |
| dblinc-uc003jss.2 | Downregulated |
| dblinc-uc003kwt.2 | Downregulated |
| dblinc-uc003kwu.1 | Downregulated |
| dblinc-uc003kyl.2 | Downregulated |
| dblinc-uc003kzj.2 | Downregulated |
| dblinc-uc003opf.1 | Downregulated |
| dblinc-uc003qir.2 | Downregulated |
| dblinc-uc003qye.1 | Downregulated |
| dblinc-uc003sop.1 | Downregulated |
| dblinc-uc004bad.1 | Downregulated |
| dblinc-uc004bdd.1 | Downregulated |
| dblinc-uc004bxt.1 | Downregulated |
| dblinc-uc004bxu.2 | Downregulated |
| dblinc-uc004bxy.1 | Downregulated |
| dblinc-uc004ecf.1 | Downregulated |
| dblinc-uc004eqg.1 | Downregulated |
| dblinc-uc004evu.2 | Downregulated |
| dblinc-uc004exn.1 | Downregulated |
| dblinc-uc010gig.1 | Downregulated |
| dblinc-uc010hbl.1 | Downregulated |
| dblinc-uc010jub.1 | Downregulated |
| DIO3OS | Downregulated |
| FKBP1A-SDCBP2 | Downregulated |
| FRY-AS1 | Downregulated |
| GNG12-AS1 | Downregulated |
| H19 | Downregulated |
| HAR1B | Downregulated |
| KB-1047C11.1 | Downregulated |
| KB-1448A5.1 | Downregulated |
| KB-1460A1.1 | Downregulated |
| KB-1732A1.1 | Downregulated |
| KIAA0040 | Downregulated |
| LA16c-325D7.2 | Downregulated |
| LINC00087 | Downregulated |
| LINC00102 | Downregulated |
| LINC00160 | Downregulated |
| LINC00173 | Downregulated |
| LINC00273 | Downregulated |
| LINC00311 | Downregulated |
| LINC00312 | Downregulated |
| LINC00313 | Downregulated |
| LINC00316 | Downregulated |
| LINC00324 | Downregulated |
| LINC00327 | Downregulated |
| LINC00338 | Downregulated |
| LINC00350 | Downregulated |
| LINC00354 | Downregulated |
| LINC00445 | Downregulated |
| LINC00454 | Downregulated |
| LINC00475 | Downregulated |
| LINC00511 | Downregulated |
| LINC00518 | Downregulated |
| LINC00533 | Downregulated |
| LINC00565 | Downregulated |
| LINC00593 | Downregulated |
| linc-ABCA5-3 | Downregulated |
| linc-ABT1 | Downregulated |
| linc-ACO1-2 | Downregulated |
| linc-ADCY9 | Downregulated |
| linc-ADRB1-1 | Downregulated |
| linc-AIRE-4 | Downregulated |
| linc-AKT3-2 | Downregulated |
| linc-ALDH1L1-1 | Downregulated |
| linc-ALG2-5 | Downregulated |
| linc-AMD1-2 | Downregulated |
| linc-AMN1 | Downregulated |
| linc-ANKRD55-3 | Downregulated |
| linc-ANKRD55-4 | Downregulated |
| linc-ANKRD55-6 | Downregulated |
| linc-ANP32A-2 | Downregulated |
| linc-APOA5 | Downregulated |
| linc-APOC3-1 | Downregulated |
| linc-ARF6-1 | Downregulated |
| linc-ARHGAP26-2 | Downregulated |
| linc-ARL1-2 | Downregulated |
| linc-ATF3-1 | Downregulated |
| linc-ATF3-2 | Downregulated |
| linc-ATHL1 | Downregulated |
| linc-ATP13A4-2 | Downregulated |
| linc-ATP13A4-8 | Downregulated |
| linc-ATP1A1 | Downregulated |
| linc-ATP6V1C1-3 | Downregulated |
| linc-ATP6V1C2-1 | Downregulated |
| linc-B4GALT1 | Downregulated |
| linc-BARHL2 | Downregulated |
| linc-BBS12-1 | Downregulated |
| linc-BBS5 | Downregulated |
| linc-BCL3 | Downregulated |
| linc-BEST3-1 | Downregulated |
| linc-BHLHE23-3 | Downregulated |
| linc-BOD1-3 | Downregulated |
| linc-BRD9-1 | Downregulated |
| linc-BRI3BP-1 | Downregulated |
| linc-BRI3BP-2 | Downregulated |
| linc-BTBD6-2 | Downregulated |
| linc-C10orf137-1 | Downregulated |
| linc-C10orf90-2 | Downregulated |
| linc-C14orf101-2 | Downregulated |
| linc-C14orf102-1 | Downregulated |
| linc-C14orf102-2 | Downregulated |
| linc-C14orf159-2 | Downregulated |
| linc-C14orf159-3 | Downregulated |
| linc-C14orf4-2 | Downregulated |
| linc-C14orf43-2 | Downregulated |
| linc-C1orf65-2 | Downregulated |
| linc-C1orf65-3 | Downregulated |
| linc-C20orf166 | Downregulated |
| linc-C22orf9 | Downregulated |
| linc-C2orf54-1 | Downregulated |
| linc-C2orf65-1 | Downregulated |
| linc-C5orf43-1 | Downregulated |
| linc-C5orf43-4 | Downregulated |
| linc-C7orf27-1 | Downregulated |
| linc-C7orf27-3 | Downregulated |
| linc-C7orf65-2 | Downregulated |
| linc-C8orf22-2 | Downregulated |
| linc-C9orf106-3 | Downregulated |
| linc-CA14 | Downregulated |
| linc-CA8-5 | Downregulated |
| linc-CACNA2D4 | Downregulated |
| linc-CALCA | Downregulated |
| linc-CBR1-3 | Downregulated |
| linc-CBX4 | Downregulated |
| linc-CCDC37-3 | Downregulated |
| linc-CCDC90A-6 | Downregulated |
| linc-CCND1-2 | Downregulated |
| linc-CCT5-2 | Downregulated |
| linc-CD9-1 | Downregulated |
| linc-CD93-1 | Downregulated |
| linc-CD93-2 | Downregulated |
| linc-CDC16-5 | Downregulated |
| linc-CDC42EP1-1 | Downregulated |
| linc-CDCA4-1 | Downregulated |
| linc-CDCA4-2 | Downregulated |
| linc-CDH11-1 | Downregulated |
| linc-CDYL-3 | Downregulated |
| linc-CEACAM20-1 | Downregulated |
| linc-CEBPB-1 | Downregulated |
| linc-CENPP-4 | Downregulated |
| linc-CENPP-5 | Downregulated |
| linc-CERK-4 | Downregulated |
| linc-CGNL1-1 | Downregulated |
| linc-CHAC1 | Downregulated |
| linc-CHD1-1 | Downregulated |
| linc-CHIC2-3 | Downregulated |
| linc-CHRNA9-2 | Downregulated |
| linc-CIB2 | Downregulated |
| linc-CLASP2-1 | Downregulated |
| linc-CLRN2-1 | Downregulated |
| linc-CMPK2-7 | Downregulated |
| linc-CNN1 | Downregulated |
| linc-CNTLN-5 | Downregulated |
| linc-CNTNAP2-3 | Downregulated |
| linc-COIL-2 | Downregulated |
| linc-COL18A1-1 | Downregulated |
| linc-COL18A1-2 | Downregulated |
| linc-COL18A1-3 | Downregulated |
| linc-COX4NB-4 | Downregulated |
| linc-CPB2 | Downregulated |
| linc-CPEB2-16 | Downregulated |
| linc-CPEB4-3 | Downregulated |
| linc-CREBBP | Downregulated |
| linc-CSTB-1 | Downregulated |
| linc-CSTB-2 | Downregulated |
| linc-CSTB-6 | Downregulated |
| linc-CSTB-7 | Downregulated |
| linc-CSTB-9 | Downregulated |
| linc-CTSD-2 | Downregulated |
| linc-CTTNBP2NL | Downregulated |
| linc-CXCR4-1 | Downregulated |
| linc-DCAF4L2 | Downregulated |
| linc-DHCR7-2 | Downregulated |
| linc-DHRS2-1 | Downregulated |
| linc-DHX37-22 | Downregulated |
| linc-DIRAS2-1 | Downregulated |
| linc-DIRAS2-3 | Downregulated |
| linc-DLGAP2-5 | Downregulated |
| linc-DLGAP2-8 | Downregulated |
| linc-DLX4-1 | Downregulated |
| linc-DMRT2 | Downregulated |
| linc-DTNBP1-2 | Downregulated |
| linc-DUSP26-1 | Downregulated |
| linc-DYRK2-1 | Downregulated |
| linc-EEPD1-2 | Downregulated |
| linc-EFR3A-7 | Downregulated |
| linc-EGR4-3 | Downregulated |
| linc-ELOVL6-1 | Downregulated |
| linc-EMB-3 | Downregulated |
| linc-ENPP4-1 | Downregulated |
| linc-ERICH1-2 | Downregulated |
| linc-ERICH1-4 | Downregulated |
| linc-EVI2A | Downregulated |
| linc-EVX2-7 | Downregulated |
| linc-EXD2-1 | Downregulated |
| linc-F13A1-2 | Downregulated |
| linc-FAIM3 | Downregulated |
| linc-FAM102A | Downregulated |
| linc-FAM122C-2 | Downregulated |
| linc-FAM194A | Downregulated |
| linc-FAM43A-5 | Downregulated |
| linc-FAM92B-1 | Downregulated |
| linc-FAM92B-4 | Downregulated |
| linc-FCHSD2-1 | Downregulated |
| linc-FGF3-1 | Downregulated |
| linc-FGFR1OP-7 | Downregulated |
| linc-FIGNL2-2 | Downregulated |
| linc-FLJ44606-2 | Downregulated |
| linc-FMOD-3 | Downregulated |
| linc-FOS-1 | Downregulated |
| linc-FOXC1-1 | Downregulated |
| linc-FOXF1-3 | Downregulated |
| linc-FRG1-4 | Downregulated |
| linc-GARNL3 | Downregulated |
| linc-GAS6-2 | Downregulated |
| linc-GBP6-4 | Downregulated |
| linc-GCNT4 | Downregulated |
| linc-GCNT7-1 | Downregulated |
| linc-GJA8-2 | Downregulated |
| linc-GLI3-2 | Downregulated |
| linc-GLT25D2-1 | Downregulated |
| linc-GPATCH4 | Downregulated |
| linc-GPR132 | Downregulated |
| linc-GPR26-1 | Downregulated |
| linc-GPRIN3-3 | Downregulated |
| linc-GPS1 | Downregulated |
| linc-GRAMD4-1 | Downregulated |
| linc-GRAMD4-2 | Downregulated |
| linc-GREB1-1 | Downregulated |
| linc-GRHL2-2 | Downregulated |
| linc-GRID1 | Downregulated |
| linc-GRIP1-7 | Downregulated |
| linc-GRPEL1-1 | Downregulated |
| linc-GTF3A | Downregulated |
| linc-GUCA2B | Downregulated |
| linc-GZMK | Downregulated |
| linc-HDDC2-3 | Downregulated |
| linc-HERC6-2 | Downregulated |
| linc-HES1-1 | Downregulated |
| linc-HHIPL1 | Downregulated |
| linc-HMGA1-2 | Downregulated |
| linc-HMGB1-2 | Downregulated |
| linc-HNRNPA2B1-3 | Downregulated |
| linc-HNRNPA3-2 | Downregulated |
| linc-HSPB8 | Downregulated |
| linc-ID4-2 | Downregulated |
| linc-IDH3A | Downregulated |
| linc-IER5L-5 | Downregulated |
| linc-IGFL1-2 | Downregulated |
| linc-JAKMIP1 | Downregulated |
| linc-JDP2 | Downregulated |
| linc-KCNMB1-1 | Downregulated |
| linc-KCTD1-2 | Downregulated |
| linc-KCTD6 | Downregulated |
| linc-KHDRBS3-5 | Downregulated |
| linc-KIAA0040 | Downregulated |
| linc-KIAA0182-1 | Downregulated |
| linc-KIAA0182-2 | Downregulated |
| linc-KIAA0232 | Downregulated |
| linc-KIAA1737-2 | Downregulated |
| linc-KIAA1737-3 | Downregulated |
| linc-KIAA1755-4 | Downregulated |
| linc-KIF13B | Downregulated |
| linc-KITLG-2 | Downregulated |
| linc-KLF11 | Downregulated |
| linc-KLHL18-3 | Downregulated |
| linc-KSR1 | Downregulated |
| linc-LAPTM4A-1 | Downregulated |
| linc-LAPTM5 | Downregulated |
| linc-LARGE-3 | Downregulated |
| linc-LCMT1 | Downregulated |
| linc-LFNG | Downregulated |
| linc-LHX1-1 | Downregulated |
| linc-LINS-4 | Downregulated |
| linc-LOC100129636-4 | Downregulated |
| linc-LOC642587-6 | Downregulated |
| linc-LOXL1 | Downregulated |
| linc-LRCH2-1 | Downregulated |
| linc-LRRC31-1 | Downregulated |
| linc-LRRC32-1 | Downregulated |
| linc-LRRC32-2 | Downregulated |
| linc-LRRC49-1 | Downregulated |
| linc-LSR | Downregulated |
| linc-LYZL2-4 | Downregulated |
| linc-LZTS1-1 | Downregulated |
| linc-MAMLD1 | Downregulated |
| linc-MAP1LC3B2-2 | Downregulated |
| linc-MAP1LC3B2-4 | Downregulated |
| linc-MAP3K1-1 | Downregulated |
| linc-MAP3K9-1 | Downregulated |
| linc-MAPKAP1-3 | Downregulated |
| linc-MBOAT2-2 | Downregulated |
| linc-MBP-4 | Downregulated |
| linc-MDM2 | Downregulated |
| linc-MED13L-2 | Downregulated |
| linc-MEGF10 | Downregulated |
| linc-MERTK-1 | Downregulated |
| linc-METAP1-2 | Downregulated |
| linc-METTL10 | Downregulated |
| linc-MKI67-4 | Downregulated |
| linc-MLLT4-1 | Downregulated |
| linc-MLN | Downregulated |
| linc-MOCS1-3 | Downregulated |
| linc-MPHOSPH8-1 | Downregulated |
| linc-MPHOSPH8-4 | Downregulated |
| linc-MPPE1-4 | Downregulated |
| linc-MRPS18A-1 | Downregulated |
| linc-MRPS31 | Downregulated |
| linc-MTERFD1 | Downregulated |
| linc-MTPAP | Downregulated |
| linc-MYOD1 | Downregulated |
| linc-NAB2 | Downregulated |
| linc-NADSYN1-1 | Downregulated |
| linc-NAIF1 | Downregulated |
| linc-NANOS1-3 | Downregulated |
| linc-NBPF15-1 | Downregulated |
| linc-NEURL1B-1 | Downregulated |
| linc-NFAM1 | Downregulated |
| linc-NIPSNAP1 | Downregulated |
| linc-NOP14-3 | Downregulated |
| linc-NOTCH1-1 | Downregulated |
| linc-NRIP1-3 | Downregulated |
| linc-NRSN1-2 | Downregulated |
| linc-NTM-4 | Downregulated |
| linc-NTM-5 | Downregulated |
| linc-NUDCD2-5 | Downregulated |
| linc-OBFC1 | Downregulated |
| linc-OR4F6-2 | Downregulated |
| linc-OSTalpha-1 | Downregulated |
| linc-OTOS-1 | Downregulated |
| linc-OTOS-2 | Downregulated |
| linc-OXGR1-2 | Downregulated |
| linc-P2RY2-1 | Downregulated |
| linc-P2RY6 | Downregulated |
| linc-PAX9-1 | Downregulated |
| linc-PCDH10-1 | Downregulated |
| linc-PCSK6-2 | Downregulated |
| linc-PDLIM4 | Downregulated |
| linc-PELI1-5 | Downregulated |
| linc-PET117 | Downregulated |
| linc-PFKFB3 | Downregulated |
| linc-PGS1-2 | Downregulated |
| linc-PGS1-3 | Downregulated |
| linc-PHF15-1 | Downregulated |
| linc-PIK3R1-3 | Downregulated |
| linc-PITX1-3 | Downregulated |
| linc-PLCB2 | Downregulated |
| linc-PLEKHG6-2 | Downregulated |
| linc-PLXNA2-3 | Downregulated |
| linc-POFUT2-1 | Downregulated |
| linc-POFUT2-4 | Downregulated |
| linc-POLE4 | Downregulated |
| linc-PPAP2A-1 | Downregulated |
| linc-PPARGC1A | Downregulated |
| linc-PPM1M-1 | Downregulated |
| linc-PPM1M-2 | Downregulated |
| linc-PPP4R1-5 | Downregulated |
| linc-PRB2-1 | Downregulated |
| linc-PRDM1-1 | Downregulated |
| linc-PRICKLE2-2 | Downregulated |
| linc-PRKACG-2 | Downregulated |
| linc-PRLR | Downregulated |
| linc-PRMT8 | Downregulated |
| linc-PRNP | Downregulated |
| linc-PRR5-2 | Downregulated |
| linc-PRSS3-3 | Downregulated |
| linc-PSMG4 | Downregulated |
| linc-PTGER4-4 | Downregulated |
| linc-PTGR2 | Downregulated |
| linc-PTPN1-1 | Downregulated |
| linc-PTPN2-2 | Downregulated |
| linc-RAB7A-2 | Downregulated |
| linc-RAB7A-4 | Downregulated |
| linc-RAD23B-1 | Downregulated |
| linc-RAD23B-2 | Downregulated |
| linc-RAP1GAP2-1 | Downregulated |
| linc-RASL11B | Downregulated |
| linc-RBKS-5 | Downregulated |
| linc-RBM45-2 | Downregulated |
| linc-RBMS1-4 | Downregulated |
| linc-RBPJ | Downregulated |
| linc-RHOBTB2-2 | Downregulated |
| linc-ROPN1B-3 | Downregulated |
| linc-RREB1-4 | Downregulated |
| linc-RRP1B-1 | Downregulated |
| linc-RRP1B-4 | Downregulated |
| linc-RTL1-3 | Downregulated |
| linc-RUFY4-1 | Downregulated |
| linc-RYK | Downregulated |
| linc-SARS | Downregulated |
| linc-SCAND1 | Downregulated |
| linc-SCAND3-4 | Downregulated |
| linc-SCPEP1 | Downregulated |
| linc-SCTR-7 | Downregulated |
| linc-SCUBE1 | Downregulated |
| linc-SCUBE2 | Downregulated |
| linc-SDCCAG1-2 | Downregulated |
| linc-SDK2-1 | Downregulated |
| linc-SEC62 | Downregulated |
| linc-SEMA3B | Downregulated |
| linc-SEMA4D-1 | Downregulated |
| linc-SEMA4D-2 | Downregulated |
| linc-SEMA6A-4 | Downregulated |
| linc-SEPP1-2 | Downregulated |
| linc-SEPT9 | Downregulated |
| linc-SERTAD2-1 | Downregulated |
| linc-SH2B3 | Downregulated |
| linc-SH3BP2 | Downregulated |
| linc-SHB | Downregulated |
| linc-SIRT1 | Downregulated |
| linc-SKIL | Downregulated |
| linc-SLC6A19-2 | Downregulated |
| linc-SLK-1 | Downregulated |
| linc-SMAD7 | Downregulated |
| linc-SNTG2-4 | Downregulated |
| linc-SOX1-1 | Downregulated |
| linc-SPATA18 | Downregulated |
| linc-SPATA8-2 | Downregulated |
| linc-SPO11-3 | Downregulated |
| linc-SPO11-4 | Downregulated |
| linc-SPOCD1 | Downregulated |
| linc-SRBD1-1 | Downregulated |
| linc-SRBD1-5 | Downregulated |
| linc-SRD5A3-2 | Downregulated |
| linc-SRD5A3-3 | Downregulated |
| linc-SRSF7 | Downregulated |
| linc-STARD10 | Downregulated |
| linc-STON2 | Downregulated |
| linc-STRBP | Downregulated |
| linc-SUZ12-4 | Downregulated |
| linc-SYNGR1-1 | Downregulated |
| linc-SYNM-2 | Downregulated |
| linc-SYNM-3 | Downregulated |
| linc-SYNPO2-1 | Downregulated |
| linc-SYT4-1 | Downregulated |
| linc-TAF1A | Downregulated |
| linc-TCP10-5 | Downregulated |
| linc-TCP11L2-1 | Downregulated |
| linc-TFAP2A-2 | Downregulated |
| linc-TFAP2C-1 | Downregulated |
| linc-TFAP2C-3 | Downregulated |
| linc-TGFBR2-2 | Downregulated |
| linc-TGM3-1 | Downregulated |
| linc-THSD4 | Downregulated |
| linc-TMCC1-3 | Downregulated |
| linc-TMEM18-9 | Downregulated |
| linc-TMEM194A | Downregulated |
| linc-TMEM99-4 | Downregulated |
| linc-TNFAIP3-6 | Downregulated |
| linc-TNFRSF19-4 | Downregulated |
| linc-TNKS2-2 | Downregulated |
| linc-TNKS-3 | Downregulated |
| linc-TNKS-5 | Downregulated |
| linc-TNS4 | Downregulated |
| linc-TOX3-2 | Downregulated |
| linc-TP53INP1 | Downregulated |
| linc-TRIM7-1 | Downregulated |
| linc-TRPC4-1 | Downregulated |
| linc-TSC22D2-1 | Downregulated |
| linc-TSKU-1 | Downregulated |
| linc-TSPAN11-2 | Downregulated |
| linc-TSPAN32-2 | Downregulated |
| linc-TSPO2-2 | Downregulated |
| linc-UBAC2-2 | Downregulated |
| linc-UBE2A | Downregulated |
| linc-UBL3-3 | Downregulated |
| linc-UMODL1-1 | Downregulated |
| linc-USP14-2 | Downregulated |
| linc-USP25-1 | Downregulated |
| linc-USP3-1 | Downregulated |
| linc-USP35 | Downregulated |
| linc-VCX3A-1 | Downregulated |
| linc-VIPR2-5 | Downregulated |
| linc-VPS8-1 | Downregulated |
| linc-VWA5B1-2 | Downregulated |
| linc-VWF | Downregulated |
| linc-WIPF3 | Downregulated |
| linc-WNT4 | Downregulated |
| linc-XPC-2 | Downregulated |
| linc-XPO7-1 | Downregulated |
| linc-XRCC6 | Downregulated |
| linc-YWHAQ-1 | Downregulated |
| linc-YWHAZ | Downregulated |
| linc-ZBED1-3 | Downregulated |
| linc-ZBTB40 | Downregulated |
| linc-ZCCHC10 | Downregulated |
| linc-ZCCHC17-1 | Downregulated |
| linc-ZCCHC17-7 | Downregulated |
| linc-ZFAND3-2 | Downregulated |
| linc-ZFHX3-4 | Downregulated |
| linc-ZFP42-10 | Downregulated |
| linc-ZKSCAN1-1 | Downregulated |
| linc-ZMIZ1-1 | Downregulated |
| linc-ZNF107-1 | Downregulated |
| linc-ZNF30-1 | Downregulated |
| linc-ZNF30-2 | Downregulated |
| linc-ZNF354B-1 | Downregulated |
| linc-ZNF354B-2 | Downregulated |
| linc-ZNF382-1 | Downregulated |
| linc-ZNF507-5 | Downregulated |
| linc-ZNF703-1 | Downregulated |
| linc-ZNF703-2 | Downregulated |
| linc-ZNF726-1 | Downregulated |
| linc-ZNF91-1 | Downregulated |
| linc-ZSWIM5 | Downregulated |
| MIR7-3HG | Downregulated |
| MLLT4-AS1 | Downregulated |
| MYLK-AS1 | Downregulated |
| PVT1 | Downregulated |
| RERG-AS1 | Downregulated |
| RP11-100G15.10 | Downregulated |
| RP11-1029J19.5 | Downregulated |
| RP11-1078H9.1 | Downregulated |
| RP11-10L7.1 | Downregulated |
| RP11-1105O14.1 | Downregulated |
| RP11-110L15.2 | Downregulated |
| RP11-112L6.4 | Downregulated |
| RP11-125I23.3 | Downregulated |
| RP11-127L20.3 | Downregulated |
| RP11-128L5.1 | Downregulated |
| RP11-128M1.1 | Downregulated |
| RP11-133K1.6 | Downregulated |
| RP11-135A1.3 | Downregulated |
| RP11-13A1.1 | Downregulated |
| RP11-146I2.1 | Downregulated |
| RP11-148B18.1 | Downregulated |
| RP11-148B18.3 | Downregulated |
| RP11-148L24.1 | Downregulated |
| RP11-150O12.1 | Downregulated |
| RP11-150O12.3 | Downregulated |
| RP11-150O12.5 | Downregulated |
| RP11-156K13.3 | Downregulated |
| RP11-15G16.1 | Downregulated |
| RP11-161M6.4 | Downregulated |
| RP11-161M6.5 | Downregulated |
| RP11-166P13.3 | Downregulated |
| RP11-167H9.6 | Downregulated |
| RP11-170M17.1 | Downregulated |
| RP11-173M1.4 | Downregulated |
| RP11-173M1.8 | Downregulated |
| RP11-174I12.2 | Downregulated |
| RP11-180M15.3 | Downregulated |
| RP11-182I10.3 | Downregulated |
| RP11-197N18.7 | Downregulated |
| RP11-199B17.1 | Downregulated |
| RP11-202P11.1 | Downregulated |
| RP11-211G3.2 | Downregulated |
| RP11-214C8.5 | Downregulated |
| RP11-227F8.2 | Downregulated |
| RP11-229C3.2 | Downregulated |
| RP11-234K24.3 | Downregulated |
| RP11-239E10.2 | Downregulated |
| RP11-245J24.1 | Downregulated |
| RP11-253D19.1 | Downregulated |
| RP11-253I19.3 | Downregulated |
| RP11-254F7.1 | Downregulated |
| RP11-254F7.2 | Downregulated |
| RP11-260E18.1 | Downregulated |
| RP11-266L9.5 | Downregulated |
| RP11-273B19.1 | Downregulated |
| RP11-273B19.2 | Downregulated |
| RP11-273G15.2 | Downregulated |
| RP11-274H24.1 | Downregulated |
| RP11-282I1.2 | Downregulated |
| RP11-285A1.1 | Downregulated |
| RP11-285E9.5 | Downregulated |
| RP11-290F20.3 | Downregulated |
| RP11-293M10.5 | Downregulated |
| RP11-299L17.3 | Downregulated |
| RP11-300M24.1 | Downregulated |
| RP11-304F15.4 | Downregulated |
| RP11-309L24.4 | Downregulated |
| RP11-321G12.1 | Downregulated |
| RP11-325L7.2 | Downregulated |
| RP11-326C3.12 | Downregulated |
| RP11-342C23.4 | Downregulated |
| RP11-344B5.4 | Downregulated |
| RP11-351M16.3 | Downregulated |
| RP11-353N14.2 | Downregulated |
| RP11-353N14.5 | Downregulated |
| RP11-353N4.4 | Downregulated |
| RP11-358M11.3 | Downregulated |
| RP11-359E8.3 | Downregulated |
| RP11-367F23.1 | Downregulated |
| RP11-367J11.3 | Downregulated |
| RP11-367J11.4 | Downregulated |
| RP11-378A12.1 | Downregulated |
| RP11-380D23.2 | Downregulated |
| RP11-384P7.7 | Downregulated |
| RP11-387H17.6 | Downregulated |
| RP11-392A22.2 | Downregulated |
| RP11-395I6.2 | Downregulated |
| RP11-399E6.1 | Downregulated |
| RP11-403P17.3 | Downregulated |
| RP11-417L19.2 | Downregulated |
| RP11-417L19.4 | Downregulated |
| RP11-419J16.1 | Downregulated |
| RP11-420G6.4 | Downregulated |
| RP11-421L21.3 | Downregulated |
| RP11-424I19.2 | Downregulated |
| RP11-430H10.3 | Downregulated |
| RP11-447B18.1 | Downregulated |
| RP11-447D11.2 | Downregulated |
| RP11-449H11.1 | Downregulated |
| RP11-452H21.4 | Downregulated |
| RP11-463H12.2 | Downregulated |
| RP11-469A15.2 | Downregulated |
| RP11-473O4.3 | Downregulated |
| RP11-475N22.4 | Downregulated |
| RP11-476M19.2 | Downregulated |
| RP11-483E7.1 | Downregulated |
| RP11-485M7.3 | Downregulated |
| RP11-488C13.4 | Downregulated |
| RP1-148H17.1 | Downregulated |
| RP11-492E3.1 | Downregulated |
| RP11-497G19.3 | Downregulated |
| RP11-4C20.3 | Downregulated |
| RP11-503G7.2 | Downregulated |
| RP11-513G11.1 | Downregulated |
| RP11-522B15.3 | Downregulated |
| RP11-522B15.4 | Downregulated |
| RP11-526K17.2 | Downregulated |
| RP11-528A4.2 | Downregulated |
| RP11-533K9.2 | Downregulated |
| RP11-533N14.3 | Downregulated |
| RP11-533O20.2 | Downregulated |
| RP11-535C7.1 | Downregulated |
| RP1-153P14.8 | Downregulated |
| RP11-542B15.1 | Downregulated |
| RP11-543C4.1 | Downregulated |
| RP11-543C4.3 | Downregulated |
| RP11-543G18.1 | Downregulated |
| RP11-548L20.1 | Downregulated |
| RP11-552E20.3 | Downregulated |
| RP11-554A11.8 | Downregulated |
| RP11-554F20.1 | Downregulated |
| RP11-557H15.3 | Downregulated |
| RP11-560A15.4 | Downregulated |
| RP11-58E21.3 | Downregulated |
| RP11-58E21.5 | Downregulated |
| RP11-598F7.6 | Downregulated |
| RP11-60A24.3 | Downregulated |
| RP11-616M22.6 | Downregulated |
| RP11-616M22.7 | Downregulated |
| RP11-619A14.3 | Downregulated |
| RP11-61J19.3 | Downregulated |
| RP11-61J19.4 | Downregulated |
| RP11-624M8.1 | Downregulated |
| RP11-626H12.3 | Downregulated |
| RP11-627G18.3 | Downregulated |
| RP11-638L3.1 | Downregulated |
| RP11-648F7.1 | Downregulated |
| RP11-65L3.1 | Downregulated |
| RP11-663N22.1 | Downregulated |
| RP11-665C16.6 | Downregulated |
| RP11-669I1.1 | Downregulated |
| RP11-670E13.2 | Downregulated |
| RP11-67M1.1 | Downregulated |
| RP11-68L18.1 | Downregulated |
| RP11-699L21.1 | Downregulated |
| RP11-69G7.1 | Downregulated |
| RP11-700H6.2 | Downregulated |
| RP11-705O3.1 | Downregulated |
| RP11-709A23.1 | Downregulated |
| RP11-71E19.1 | Downregulated |
| RP11-72M17.1 | Downregulated |
| RP11-731F5.1 | Downregulated |
| RP11-752D24.3 | Downregulated |
| RP11-761E20.1 | Downregulated |
| RP11-76C10.5 | Downregulated |
| RP11-770E5.3 | Downregulated |
| RP11-77I22.2 | Downregulated |
| RP11-7F17.3 | Downregulated |
| RP11-7F17.7 | Downregulated |
| RP11-800A3.3 | Downregulated |
| RP11-806O11.1 | Downregulated |
| RP11-834C11.3 | Downregulated |
| RP11-83B20.1 | Downregulated |
| RP11-849I19.1 | Downregulated |
| RP11-84A19.2 | Downregulated |
| RP11-865I6.2 | Downregulated |
| RP11-87C12.5 | Downregulated |
| RP11-881M11.8 | Downregulated |
| RP11-887P2.1 | Downregulated |
| RP11-88H9.1 | Downregulated |
| RP11-88H9.2 | Downregulated |
| RP11-90J7.2 | Downregulated |
| RP11-923I11.1 | Downregulated |
| RP11-923I11.4 | Downregulated |
| RP11-95P2.3 | Downregulated |
| RP11-991C1.2 | Downregulated |
| RP11-9M16.2 | Downregulated |
| RP1-224A6.3 | Downregulated |
| RP1-261G23.5 | Downregulated |
| RP1-274L7.1 | Downregulated |
| RP1-302G2.5 | Downregulated |
| RP13-25N22.1 | Downregulated |
| RP1-41C23.1 | Downregulated |
| RP1-62D2.3 | Downregulated |
| RP1-80N2.2 | Downregulated |
| RP3-416J7.4 | Downregulated |
| RP3-468B3.2 | Downregulated |
| RP3-468B3.3 | Downregulated |
| RP3-470L22.1 | Downregulated |
| RP4-534N18.2 | Downregulated |
| RP4-550H1.4 | Downregulated |
| RP4-550H1.5 | Downregulated |
| RP4-550H1.6 | Downregulated |
| RP4-561L24.3 | Downregulated |
| RP4-564F22.2 | Downregulated |
| RP4-644L1.2 | Downregulated |
| RP4-665J23.1 | Downregulated |
| RP4-754E20__A.5 | Downregulated |
| RP4-781K5.5 | Downregulated |
| RP4-794H19.4 | Downregulated |
| RP5-1021I20.2 | Downregulated |
| RP5-1028K7.2 | Downregulated |
| RP5-1050E16.1 | Downregulated |
| RP5-1096D14.2 | Downregulated |
| RP5-1114G22.2 | Downregulated |
| RP5-1125A11.4 | Downregulated |
| RP5-1125N11.1 | Downregulated |
| RP5-1157M23.2 | Downregulated |
| RP5-1185H19.2 | Downregulated |
| RP5-1186N24.3 | Downregulated |
| RP5-1198O20.4 | Downregulated |
| RP5-848E13.3 | Downregulated |
| RP5-872K7.7 | Downregulated |
| RP5-994D16.3 | Downregulated |
| RP6-91H8.2 | Downregulated |
| SNHG15 | Downregulated |
| TPRG1-AS1 | Downregulated |
| TSGA10IP | Downregulated |
| TTC28-AS1 | Downregulated |
| WI2-1959D15.1 | Downregulated |
| Z83851.3 | Downregulated |
| ZNF295-AS1 | Downregulated |
