## Supplementary material for "Role of lncRNAs related to NRs in the regulation of gene expression": Table S2

Table S. 2: The 487 transcripts are targets of has-miR-1908-3p according to the TargetScanS (Release 8.0) database. The cumulative weighted context++ score (CWCS) is an option for ranking predicted targets of mammalian miRNAs. For each predicted target, the CWCS evaluates the total suppression expected from multiple sites to the same miRNA. This score is estimated using the 3′-UTR profiles to weigh the minimal effect of each additional site on the miRNA while also taking into account the predicted mRNA depletion resulting from any downstream sites to the same miRNA. The Total context++ score is generated to rank the predicted targets and is calculated by summing the context++ scores for the sites to the representative miRNA (17).

| Target genes | Gene name | Representative miRNA | Cumulative weighted context++ score | Total context++ score |
| --- | --- | --- | --- | --- |
| C19orf26 | chromosome 19 open reading frame 26 | hsa-miR-1908-3p | -1,3 | -1,3 |
| REPIN1 | replication initiator 1 | hsa-miR-1908-3p | -1,27 | -1,27 |
| CTD-2207O23.12 (TEX45 in humans) | Uncharacterized protein | hsa-miR-1908-3p | -1,21 | -1,62 |
| KCTD5 | potassium channel tetramerization domain containing 5 | hsa-miR-1908-3p | -1,13 | -1,13 |
| MZF1 | myeloid zinc finger 1 | hsa-miR-1908-3p | -1,07 | -1,07 |
| MTA1 | metastasis associated 1 | hsa-miR-1908-3p | -1,03 | -1,03 |
| TRMT112 | tRNA methyltransferase 11-2 homolog (S. cerevisiae) | hsa-miR-1908-3p | -1,01 | -1,03 |
| NRG2 | neuregulin 2 | hsa-miR-1908-3p | -0,96 | -0,96 |
| GNPTG | N-acetylglucosamine-1-phosphate transferase, gamma subunit | hsa-miR-1908-3p | -0,9 | -1,59 |
| CRABP2 | cellular retinoic acid binding protein 2 | hsa-miR-1908-3p | -0,9 | -0,9 |
| VWA1 | von Willebrand factor A domain containing 1 | hsa-miR-1908-3p | -0,87 | -0,87 |
| LY6E | lymphocyte antigen 6 complex, locus E | hsa-miR-1908-3p | -0,86 | -0,86 |
| H2AFX | H2A histone family, member X | hsa-miR-1908-3p | -0,85 | -0,85 |
| CACNA1A | calcium channel, voltage-dependent, P/Q type, alpha 1A subunit | hsa-miR-1908-3p | -0,84 | -0,84 |
| GADD45B | growth arrest and DNA-damage-inducible, beta | hsa-miR-1908-3p | -0,84 | -0,84 |
| CASZ1 | castor zinc finger 1 | hsa-miR-1908-3p | -0,82 | -0,82 |
| ONECUT3 | one cut homeobox 3 | hsa-miR-1908-3p | -0,8 | -0,8 |
| ITGB2 | integrin, beta 2 (complement component 3 receptor 3 and 4 subunit) | hsa-miR-1908-3p | -0,8 | -0,8 |
| ATP6V0E2 | ATPase, H+ transporting V0 subunit e2 | hsa-miR-1908-3p | -0,8 | -0,8 |
| PITX1 | paired-like homeodomain 1 | hsa-miR-1908-3p | -0,78 | -0,78 |
| ZDHHC1 | zinc finger, DHHC-type containing 1 | hsa-miR-1908-3p | -0,77 | -0,77 |
| STK11 | serine/threonine kinase 11 | hsa-miR-1908-3p | -0,76 | -0,87 |
| GRB2 | growth factor receptor-bound protein 2 | hsa-miR-1908-3p | -0,74 | -0,76 |
| SSTR5 | somatostatin receptor 5 | hsa-miR-1908-3p | -0,74 | -0,74 |
| C5orf38 | chromosome 5 open reading frame 38 | hsa-miR-1908-3p | -0,73 | -0,73 |
| BTBD19 | BTB (POZ) domain containing 19 | hsa-miR-1908-3p | -0,73 | -0,73 |
| IGFBP3 | insulin-like growth factor binding protein 3 | hsa-miR-1908-3p | -0,73 | -0,76 |
| C1orf229 | chromosome 1 open reading frame 229 | hsa-miR-1908-3p | -0,71 | -0,71 |
| NRARP | NOTCH-regulated ankyrin repeat protein | hsa-miR-1908-3p | -0,71 | -0,71 |
| PARP10 | poly (ADP-ribose) polymerase family, member 10 | hsa-miR-1908-3p | -0,7 | -0,7 |
| DYNLL2 | dynein, light chain, LC8-type 2 | hsa-miR-1908-3p | -0,69 | -0,76 |
| SPON2 | spondin 2, extracellular matrix protein | hsa-miR-1908-3p | -0,67 | -0,67 |
| KIFC2 | kinesin family member C2 | hsa-miR-1908-3p | -0,67 | -0,67 |
| SDF4 | stromal cell derived factor 4 | hsa-miR-1908-3p | -0,67 | -0,67 |
| MAPK4 | mitogen-activated protein kinase 4 | hsa-miR-1908-3p | -0,67 | -0,67 |
| RCAN3 | RCAN family member 3 | hsa-miR-1908-3p | -0,66 | -0,66 |
| H1FX | H1 histone family, member X | hsa-miR-1908-3p | -0,66 | -0,67 |
| ZIC1 | Zic family member 1 | hsa-miR-1908-3p | -0,66 | -0,72 |
| CAMK2N2 | calcium/calmodulin-dependent protein kinase II inhibitor 2 | hsa-miR-1908-3p | -0,65 | -0,65 |
| CDH4 | cadherin 4, type 1, R-cadherin (retinal) | hsa-miR-1908-3p | -0,65 | -0,65 |
| RTN4R | reticulon 4 receptor | hsa-miR-1908-3p | -0,64 | -0,64 |
| DCAF4L1 | DDB1 and CUL4 associated factor 4-like 1 | hsa-miR-1908-3p | -0,64 | -0,64 |
| VPS26B | vacuolar protein sorting 26 homolog B (S. pombe) | hsa-miR-1908-3p | -0,64 | -0,7 |
| ST3GAL2 | ST3 beta-galactoside alpha-2,3-sialyltransferase 2 | hsa-miR-1908-3p | -0,63 | -0,87 |
| LMOD3 | leiomodin 3 (fetal) | hsa-miR-1908-3p | -0,63 | -0,63 |
| CACNB1 | calcium channel, voltage-dependent, beta 1 subunit | hsa-miR-1908-3p | -0,6 | -0,6 |
| FOXL2 | forkhead box L2 | hsa-miR-1908-3p | -0,6 | -0,6 |
| TNRC6C | trinucleotide repeat containing 6C | hsa-miR-1908-3p | -0,59 | -0,91 |
| GDF7 | growth differentiation factor 7 | hsa-miR-1908-3p | -0,59 | -0,59 |
| EFNA2 | ephrin-A2 | hsa-miR-1908-3p | -0,58 | -0,58 |
| TCERG1L | transcription elongation regulator 1-like | hsa-miR-1908-3p | -0,56 | -0,56 |
| ZNF784 | zinc finger protein 784 | hsa-miR-1908-3p | -0,56 | -0,56 |
| ELOVL2 | ELOVL fatty acid elongase 2 | hsa-miR-1908-3p | -0,55 | -0,61 |
| RADIL | Ras association and DIL domains | hsa-miR-1908-3p | -0,55 | -0,55 |
| ADAMTS16 | ADAM metallopeptidase with thrombospondin type 1 motif, 16 | hsa-miR-1908-3p | -0,55 | -0,55 |
| ABHD17A | abhydrolase domain containing 17A | hsa-miR-1908-3p | -0,54 | -0,55 |
| METTL21A | methyltransferase like 21A | hsa-miR-1908-3p | -0,54 | -2,62 |
| ID4 | inhibitor of DNA binding 4, dominant negative helix-loop-helix protein | hsa-miR-1908-3p | -0,52 | -0,52 |
| ADAT3 | adenosine deaminase, tRNA-specific 3 | hsa-miR-1908-3p | -0,52 | -0,52 |
| FAM19A5 | family with sequence similarity 19 (chemokine (C-C motif)-like), member A5 | hsa-miR-1908-3p | -0,51 | -0,58 |
| RPL37 | ribosomal protein L37 | hsa-miR-1908-3p | -0,51 | -0,62 |
| HIST3H2BB | histone cluster 3, H2bb | hsa-miR-1908-3p | -0,5 | -0,88 |
| POLR2J | polymerase (RNA) II (DNA directed) polypeptide J, 13.3kDa | hsa-miR-1908-3p | -0,5 | -0,5 |
| ZDHHC8 | zinc finger, DHHC-type containing 8 | hsa-miR-1908-3p | -0,5 | -0,5 |
| AKAP17A | A kinase (PRKA) anchor protein 17A | hsa-miR-1908-3p | -0,49 | -0,49 |
| RPUSD1 | RNA pseudouridylate synthase domain containing 1 | hsa-miR-1908-3p | -0,49 | -0,49 |
| NNAT | neuronatin | hsa-miR-1908-3p | -0,49 | -0,49 |
| SNX8 | sorting nexin 8 | hsa-miR-1908-3p | -0,48 | -0,49 |
| LTBP4 | latent transforming growth factor beta binding protein 4 | hsa-miR-1908-3p | -0,48 | -0,48 |
| MURC | muscle-related coiled-coil protein | hsa-miR-1908-3p | -0,47 | -0,47 |
| PIGG | phosphatidylinositol glycan anchor biosynthesis, class G | hsa-miR-1908-3p | -0,47 | -0,47 |
| BRSK2 | BR serine/threonine kinase 2 | hsa-miR-1908-3p | -0,47 | -0,61 |
| OSGIN1 | oxidative stress induced growth inhibitor 1 | hsa-miR-1908-3p | -0,47 | -0,47 |
| TST | thiosulfate sulfurtransferase (rhodanese) | hsa-miR-1908-3p | -0,47 | -0,47 |
| NPBWR1 | neuropeptides B/W receptor 1 | hsa-miR-1908-3p | -0,46 | -0,6 |
| POU3F4 | POU class 3 homeobox 4 | hsa-miR-1908-3p | -0,45 | -0,45 |
| CAMSAP3 | calmodulin regulated spectrin-associated protein family, member 3 | hsa-miR-1908-3p | -0,45 | -0,45 |
| LPPR3 | hsa-mir-3187 | hsa-miR-1908-3p | -0,45 | -0,45 |
| CASKIN1 | CASK interacting protein 1 | hsa-miR-1908-3p | -0,45 | -0,45 |
| SH3GLB2 | SH3-domain GRB2-like endophilin B2 | hsa-miR-1908-3p | -0,45 | -0,45 |
| GALNT9 | UDP-N-acetyl-alpha-D-galactosamine:polypeptide N-acetylgalactosaminyltransferase 9 (GalNAc-T9) | hsa-miR-1908-3p | -0,44 | -0,44 |
| RBPJL | recombination signal binding protein for immunoglobulin kappa J region-like | hsa-miR-1908-3p | -0,44 | -0,44 |
| RHBDF1 | rhomboid 5 homolog 1 (Drosophila) | hsa-miR-1908-3p | -0,44 | -0,44 |
| IRF7 | interferon regulatory factor 7 | hsa-miR-1908-3p | -0,44 | -0,44 |
| HAPLN2 | hyaluronan and proteoglycan link protein 2 | hsa-miR-1908-3p | -0,44 | -0,44 |
| MEX3D | mex-3 RNA binding family member D | hsa-miR-1908-3p | -0,43 | -0,43 |
| C8orf82 | chromosome 8 open reading frame 82 | hsa-miR-1908-3p | -0,43 | -0,43 |
| UBE2M | ubiquitin-conjugating enzyme E2M | hsa-miR-1908-3p | -0,43 | -0,43 |
| FAM43B | family with sequence similarity 43, member B | hsa-miR-1908-3p | -0,43 | -0,43 |
| FAM101B | family with sequence similarity 101, member B | hsa-miR-1908-3p | -0,43 | -0,43 |
| HDAC4 | histone deacetylase 4 | hsa-miR-1908-3p | -0,42 | -0,42 |
| ACKR2 | atypical chemokine receptor 2 | hsa-miR-1908-3p | -0,42 | -0,42 |
| AP000695.1 |  | hsa-miR-1908-3p | -0,42 | -0,42 |
| CPLX1 | complexin 1 | hsa-miR-1908-3p | -0,42 | -0,42 |
| SIX5 | SIX homeobox 5 | hsa-miR-1908-3p | -0,42 | -0,42 |
| BEGAIN | brain-enriched guanylate kinase-associated | hsa-miR-1908-3p | -0,42 | -0,42 |
| CERS1 | ceramide synthase 1 | hsa-miR-1908-3p | -0,41 | -0,41 |
| EHBP1L1 | EH domain binding protein 1-like 1 | hsa-miR-1908-3p | -0,41 | -0,41 |
| DMPK | dystrophia myotonica-protein kinase | hsa-miR-1908-3p | -0,41 | -0,42 |
| FAM195A | family with sequence similarity 195, member A | hsa-miR-1908-3p | -0,41 | -0,41 |
| FAM20C | family with sequence similarity 20, member C | hsa-miR-1908-3p | -0,41 | -0,41 |
| TMEM121 | transmembrane protein 121 | hsa-miR-1908-3p | -0,41 | -0,41 |
| ZNF85 | zinc finger protein 85 | hsa-miR-1908-3p | -0,41 | -0,72 |
| TLX3 | T-cell leukemia homeobox 3 | hsa-miR-1908-3p | -0,4 | -0,48 |
| CRTC1 | CREB regulated transcription coactivator 1 | hsa-miR-1908-3p | -0,4 | -0,42 |
| SLC25A47 | solute carrier family 25, member 47 | hsa-miR-1908-3p | -0,4 | -0,4 |
| FKSG62 |  | hsa-miR-1908-3p | -0,4 | -0,4 |
| C8orf17 | chromosome 8 open reading frame 17 | hsa-miR-1908-3p | -0,4 | -0,4 |
| FOXF1 | forkhead box F1 | hsa-miR-1908-3p | -0,4 | -0,4 |
| FOXK1 | forkhead box K1 | hsa-miR-1908-3p | -0,4 | -0,4 |
| OSR1 | odd-skipped related 1 (Drosophila) | hsa-miR-1908-3p | -0,4 | -0,4 |
| TNFSF13 | tumor necrosis factor (ligand) superfamily, member 13 | hsa-miR-1908-3p | -0,4 | -0,4 |
| SCNN1D | sodium channel, non-voltage-gated 1, delta subunit | hsa-miR-1908-3p | -0,4 | -0,4 |
| EVX1 | even-skipped homeobox 1 | hsa-miR-1908-3p | -0,4 | -0,49 |
| NKX2-5 | NK2 homeobox 5 | hsa-miR-1908-3p | -0,39 | -0,39 |
| ING5 | inhibitor of growth family, member 5 | hsa-miR-1908-3p | -0,39 | -0,39 |
| CCNB1IP1 | cyclin B1 interacting protein 1, E3 ubiquitin protein ligase | hsa-miR-1908-3p | -0,39 | -0,86 |
| WNT7B | wingless-type MMTV integration site family, member 7B | hsa-miR-1908-3p | -0,39 | -0,39 |
| NELFB | negative elongation factor complex member B | hsa-miR-1908-3p | -0,39 | -0,39 |
| HOXA7 | homeobox A7 | hsa-miR-1908-3p | -0,39 | -0,39 |
| AP000974.1 | CDNA FLJ26432 fis, clone KDN01418; Uncharacterized protein | hsa-miR-1908-3p | -0,39 | -0,39 |
| LRRC45 | leucine rich repeat containing 45 | hsa-miR-1908-3p | -0,39 | -0,39 |
| HMCN2 | hemicentin 2 | hsa-miR-1908-3p | -0,39 | -0,39 |
| BARHL1 | BarH-like homeobox 1 | hsa-miR-1908-3p | -0,39 | -0,39 |
| TNFSF12-TNFSF13 | TNFSF12-TNFSF13 readthrough | hsa-miR-1908-3p | -0,39 | -0,39 |
| SLC12A5 | solute carrier family 12 (potassium/chloride transporter), member 5 | hsa-miR-1908-3p | -0,39 | -0,39 |
| MGRN1 | mahogunin ring finger 1, E3 ubiquitin protein ligase | hsa-miR-1908-3p | -0,38 | -0,38 |
| CHFR | checkpoint with forkhead and ring finger domains, E3 ubiquitin protein ligase | hsa-miR-1908-3p | -0,38 | -0,38 |
| SHOX | short stature homeobox | hsa-miR-1908-3p | -0,38 | -0,38 |
| TMEM43 | transmembrane protein 43 | hsa-miR-1908-3p | -0,38 | -0,38 |
| WDR4 | WD repeat domain 4 | hsa-miR-1908-3p | -0,38 | -0,38 |
| GLTPD2 | glycolipid transfer protein domain containing 2 | hsa-miR-1908-3p | -0,38 | -0,38 |
| FBXW5 | F-box and WD repeat domain containing 5 | hsa-miR-1908-3p | -0,38 | -0,38 |
| NTN3 | netrin 3 | hsa-miR-1908-3p | -0,38 | -0,38 |
| KRBA2 | KRAB-A domain containing 2 | hsa-miR-1908-3p | -0,37 | -0,62 |
| PPARA | peroxisome proliferator-activated receptor alpha | hsa-miR-1908-3p | -0,37 | -0,45 |
| HOXB8 | homeobox B8 | hsa-miR-1908-3p | -0,37 | -0,37 |
| ANO8 | anoctamin 8 | hsa-miR-1908-3p | -0,37 | -0,37 |
| TPGS1 | tubulin polyglutamylase complex subunit 1 | hsa-miR-1908-3p | -0,37 | -0,37 |
| UBE2J2 | ubiquitin-conjugating enzyme E2, J2 | hsa-miR-1908-3p | -0,36 | -0,49 |
| PFKFB3 | 6-phosphofructo-2-kinase/fructose-2,6-biphosphatase 3 | hsa-miR-1908-3p | -0,36 | -0,36 |
| ZNF791 | zinc finger protein 791 | hsa-miR-1908-3p | -0,36 | -0,37 |
| AC005481.5 | Uncharacterized protein | hsa-miR-1908-3p | -0,36 | -0,36 |
| ZFYVE28 | zinc finger, FYVE domain containing 28 | hsa-miR-1908-3p | -0,36 | -0,74 |
| PIK3R2 | phosphoinositide-3-kinase, regulatory subunit 2 (beta) | hsa-miR-1908-3p | -0,36 | -0,36 |
| WDR24 | WD repeat domain 24 | hsa-miR-1908-3p | -0,36 | -0,36 |
| AP005482.1 | Uncharacterized protein | hsa-miR-1908-3p | -0,36 | -0,36 |
| AC008267.1 | HCG1983814; Uncharacterized protein | hsa-miR-1908-3p | -0,36 | -0,36 |
| CEMP1 | cementum protein 1 | hsa-miR-1908-3p | -0,35 | -0,35 |
| IL17RE | interleukin 17 receptor E | hsa-miR-1908-3p | -0,35 | -0,35 |
| LGI4 | leucine-rich repeat LGI family, member 4 | hsa-miR-1908-3p | -0,35 | -0,35 |
| MVD | mevalonate (diphospho) decarboxylase | hsa-miR-1908-3p | -0,35 | -0,35 |
| DDR1 | discoidin domain receptor tyrosine kinase 1 | hsa-miR-1908-3p | -0,35 | -0,35 |
| UAP1L1 | UDP-N-acteylglucosamine pyrophosphorylase 1-like 1 | hsa-miR-1908-3p | -0,35 | -0,35 |
| NKAIN4 | Na+/K+ transporting ATPase interacting 4 | hsa-miR-1908-3p | -0,34 | -0,34 |
| MIF4GD | MIF4G domain containing | hsa-miR-1908-3p | -0,34 | -0,34 |
| HDAC10 | histone deacetylase 10 | hsa-miR-1908-3p | -0,34 | -0,34 |
| ZIC4 | Zic family member 4 | hsa-miR-1908-3p | -0,34 | -0,34 |
| GLTPD1 | glycolipid transfer protein domain containing 1 | hsa-miR-1908-3p | -0,34 | -0,34 |
| ZNF383 | zinc finger protein 383 | hsa-miR-1908-3p | -0,34 | -0,4 |
| AMDHD2 | amidohydrolase domain containing 2 | hsa-miR-1908-3p | -0,34 | -0,38 |
| MKRN2 | makorin ring finger protein 2 | hsa-miR-1908-3p | -0,34 | -0,34 |
| EBF3 | early B-cell factor 3 | hsa-miR-1908-3p | -0,34 | -0,34 |
| TNFRSF18 | tumor necrosis factor receptor superfamily, member 18 | hsa-miR-1908-3p | -0,34 | -0,34 |
| HRAS | Harvey rat sarcoma viral oncogene homolog | hsa-miR-1908-3p | -0,34 | -0,34 |
| PTBP1 | polypyrimidine tract binding protein 1 | hsa-miR-1908-3p | -0,34 | -0,34 |
| RP11-156E8.1 |  | hsa-miR-1908-3p | -0,33 | -0,41 |
| ANO4 | anoctamin 4 | hsa-miR-1908-3p | -0,33 | -0,33 |
| SLC5A10 | solute carrier family 5 (sodium/sugar cotransporter), member 10 | hsa-miR-1908-3p | -0,33 | -0,33 |
| AL591479.1 | Uncharacterized protein | hsa-miR-1908-3p | -0,33 | -0,33 |
| EFCAB1 | EF-hand calcium binding domain 1 | hsa-miR-1908-3p | -0,33 | -0,33 |
| REM2 | RAS (RAD and GEM)-like GTP binding 2 | hsa-miR-1908-3p | -0,33 | -0,33 |
| SNPH | syntaphilin | hsa-miR-1908-3p | -0,33 | -0,33 |
| PTTG1IP | pituitary tumor-transforming 1 interacting protein | hsa-miR-1908-3p | -0,33 | -0,33 |
| CHTF8 | CTF8, chromosome transmission fidelity factor 8 homolog (S. cerevisiae) | hsa-miR-1908-3p | -0,33 | -0,33 |
| HMX1 | H6 family homeobox 1 | hsa-miR-1908-3p | -0,33 | -0,5 |
| NBEAL2 | neurobeachin-like 2 | hsa-miR-1908-3p | -0,33 | -0,33 |
| C14orf180 | chromosome 14 open reading frame 180 | hsa-miR-1908-3p | -0,33 | -0,33 |
| UBE4B | ubiquitination factor E4B | hsa-miR-1908-3p | -0,32 | -0,32 |
| NLRP1 | NLR family, pyrin domain containing 1 | hsa-miR-1908-3p | -0,32 | -0,33 |
| COL8A2 | collagen, type VIII, alpha 2 | hsa-miR-1908-3p | -0,32 | -0,32 |
| HDLBP | high density lipoprotein binding protein | hsa-miR-1908-3p | -0,32 | -0,33 |
| KCNAB3 | potassium voltage-gated channel, shaker-related subfamily, beta member 3 | hsa-miR-1908-3p | -0,32 | -0,32 |
| DYNLT3 | dynein, light chain, Tctex-type 3 | hsa-miR-1908-3p | -0,32 | -0,32 |
| HTRA1 | HtrA serine peptidase 1 | hsa-miR-1908-3p | -0,32 | -0,32 |
| PPP1R27 | protein phosphatase 1, regulatory subunit 27 | hsa-miR-1908-3p | -0,32 | -0,32 |
| WHSC1 | Wolf-Hirschhorn syndrome candidate 1 | hsa-miR-1908-3p | -0,32 | -0,32 |
| AL162389.1 | Uncharacterized protein | hsa-miR-1908-3p | -0,31 | -0,32 |
| MLST8 | MTOR associated protein, LST8 homolog (S. cerevisiae) | hsa-miR-1908-3p | -0,31 | -0,32 |
| DGCR2 | DiGeorge syndrome critical region gene 2 | hsa-miR-1908-3p | -0,31 | -0,33 |
| TUBB3 | Tubulin beta-3 chain | hsa-miR-1908-3p | -0,31 | -0,31 |
| JUN | jun proto-oncogene | hsa-miR-1908-3p | -0,31 | -0,31 |
| HIVEP3 | human immunodeficiency virus type I enhancer binding protein 3 | hsa-miR-1908-3p | -0,31 | -0,31 |
| RGMA | RGM domain family, member A | hsa-miR-1908-3p | -0,31 | -0,32 |
| VDR | vitamin D (1,25- dihydroxyvitamin D3) receptor | hsa-miR-1908-3p | -0,31 | -0,31 |
| C6orf195 | chromosome 6 open reading frame 195 | hsa-miR-1908-3p | -0,31 | -0,31 |
| DLL4 | delta-like 4 (Drosophila) | hsa-miR-1908-3p | -0,31 | -0,31 |
| PAX2 | paired box 2 | hsa-miR-1908-3p | -0,31 | -0,31 |
| MYADML2 | myeloid-associated differentiation marker-like 2 | hsa-miR-1908-3p | -0,31 | -0,31 |
| UBE2H | ubiquitin-conjugating enzyme E2H | hsa-miR-1908-3p | -0,3 | -0,32 |
| OR2C3 | olfactory receptor, family 2, subfamily C, member 3 | hsa-miR-1908-3p | -0,3 | -0,3 |
| JUND | jun D proto-oncogene | hsa-miR-1908-3p | -0,3 | -0,3 |
| UNCX | UNC homeobox | hsa-miR-1908-3p | -0,3 | -0,3 |
| AGAP1 | ArfGAP with GTPase domain, ankyrin repeat and PH domain 1 | hsa-miR-1908-3p | -0,3 | -0,3 |
| C9orf47 | chromosome 9 open reading frame 47 | hsa-miR-1908-3p | -0,3 | -0,3 |
| CHMP1A | charged multivesicular body protein 1A | hsa-miR-1908-3p | -0,3 | -0,3 |
| CAMKK2 | calcium/calmodulin-dependent protein kinase kinase 2, beta | hsa-miR-1908-3p | -0,3 | -0,3 |
| BOK | BCL2-related ovarian killer | hsa-miR-1908-3p | -0,3 | -0,3 |
| LARGE | like-glycosyltransferase | hsa-miR-1908-3p | -0,3 | -0,3 |
| TPPP | tubulin polymerization promoting protein | hsa-miR-1908-3p | -0,3 | -0,3 |
| CPEB3 | cytoplasmic polyadenylation element binding protein 3 | hsa-miR-1908-3p | -0,3 | -0,3 |
| TRABD | TraB domain containing | hsa-miR-1908-3p | -0,29 | -0,3 |
| GAS2L1 | growth arrest-specific 2 like 1 | hsa-miR-1908-3p | -0,29 | -0,29 |
| NRBP1 | nuclear receptor binding protein 1 | hsa-miR-1908-3p | -0,29 | -0,29 |
| COMT | catechol-O-methyltransferase | hsa-miR-1908-3p | -0,29 | -0,29 |
| AGPAT4 | 1-acylglycerol-3-phosphate O-acyltransferase 4 | hsa-miR-1908-3p | -0,29 | -0,59 |
| PLXNA3 | plexin A3 | hsa-miR-1908-3p | -0,29 | -0,29 |
| RAP2B | RAP2B, member of RAS oncogene family | hsa-miR-1908-3p | -0,29 | -0,29 |
| C21orf2 | chromosome 21 open reading frame 2 | hsa-miR-1908-3p | -0,29 | -0,29 |
| RAB40C | RAB40C, member RAS oncogene family | hsa-miR-1908-3p | -0,29 | -0,29 |
| CCDC180 | coiled-coil domain containing 180 | hsa-miR-1908-3p | -0,29 | -0,29 |
| LDLRAD4 | low density lipoprotein receptor class A domain containing 4 | hsa-miR-1908-3p | -0,29 | -0,29 |
| RDH13 | retinol dehydrogenase 13 (all-trans/9-cis) | hsa-miR-1908-3p | -0,29 | -0,29 |
| C17orf70 | chromosome 17 open reading frame 70 | hsa-miR-1908-3p | -0,29 | -0,29 |
| COL18A1 | collagen, type XVIII, alpha 1 | hsa-miR-1908-3p | -0,28 | -0,28 |
| RGS19 | regulator of G-protein signaling 19 | hsa-miR-1908-3p | -0,28 | -0,28 |
| RNF151 | ring finger protein 151 | hsa-miR-1908-3p | -0,28 | -0,28 |
| ASCL2 | achaete-scute complex homolog 2 (Drosophila) | hsa-miR-1908-3p | -0,28 | -0,28 |
| ZDHHC15 | zinc finger, DHHC-type containing 15 | hsa-miR-1908-3p | -0,28 | -0,28 |
| CES3 | carboxylesterase 3 | hsa-miR-1908-3p | -0,28 | -0,46 |
| KCNQ4 | potassium voltage-gated channel, KQT-like subfamily, member 4 | hsa-miR-1908-3p | -0,28 | -0,28 |
| PIP5K1C | phosphatidylinositol-4-phosphate 5-kinase, type I, gamma | hsa-miR-1908-3p | -0,28 | -0,28 |
| CTD-2162K18.4 | Uncharacterized protein | hsa-miR-1908-3p | -0,27 | -0,27 |
| ALKBH6 | alkB, alkylation repair homolog 6 (E. coli) | hsa-miR-1908-3p | -0,27 | -0,27 |
| ULK1 | unc-51 like autophagy activating kinase 1 | hsa-miR-1908-3p | -0,27 | -0,27 |
| LMTK2 | lemur tyrosine kinase 2 | hsa-miR-1908-3p | -0,27 | -0,4 |
| PLXNA1 | plexin A1 | hsa-miR-1908-3p | -0,27 | -0,27 |
| ACTRT3 | actin-related protein T3 | hsa-miR-1908-3p | -0,27 | -0,45 |
| EPN3 | epsin 3 | hsa-miR-1908-3p | -0,27 | -0,27 |
| ZFHX2 | zinc finger homeobox 2 | hsa-miR-1908-3p | -0,27 | -0,27 |
| ARFGAP1 | ADP-ribosylation factor GTPase activating protein 1 | hsa-miR-1908-3p | -0,27 | -0,27 |
| UPF1 | UPF1 regulator of nonsense transcripts homolog (yeast) | hsa-miR-1908-3p | -0,27 | -0,28 |
| UNC5A | unc-5 homolog A (C. elegans) | hsa-miR-1908-3p | -0,26 | -0,26 |
| SLC11A1 | solute carrier family 11 (proton-coupled divalent metal ion transporter), member 1 | hsa-miR-1908-3p | -0,26 | -0,26 |
| KLF2 | Kruppel-like factor 2 (lung) | hsa-miR-1908-3p | -0,26 | -0,26 |
| STARD10 | StAR-related lipid transfer (START) domain containing 10 | hsa-miR-1908-3p | -0,26 | -0,36 |
| EID2B | EP300 interacting inhibitor of differentiation 2B | hsa-miR-1908-3p | -0,26 | -0,35 |
| TSGA13 | testis specific, 13 | hsa-miR-1908-3p | -0,26 | -0,26 |
| PHOX2B | paired-like homeobox 2b | hsa-miR-1908-3p | -0,26 | -0,26 |
| GPR115 | G protein-coupled receptor 115 | hsa-miR-1908-3p | -0,26 | -0,26 |
| DHRS12 | dehydrogenase/reductase (SDR family) member 12 | hsa-miR-1908-3p | -0,26 | -0,26 |
| PRKAR1B | protein kinase, cAMP-dependent, regulatory, type I, beta | hsa-miR-1908-3p | -0,26 | -0,27 |
| SALL3 | sal-like 3 (Drosophila) | hsa-miR-1908-3p | -0,26 | -0,4 |
| TEX22 | testis expressed 22 | hsa-miR-1908-3p | -0,26 | -0,66 |
| NCLN | nicalin | hsa-miR-1908-3p | -0,25 | -0,38 |
| EIF4H | eukaryotic translation initiation factor 4H | hsa-miR-1908-3p | -0,25 | -0,25 |
| IGF2 | insulin-like growth factor 2 (somatomedin A) | hsa-miR-1908-3p | -0,25 | -0,33 |
| TBL1X | transducin (beta)-like 1X-linked | hsa-miR-1908-3p | -0,25 | -0,29 |
| SMOC2 | SPARC related modular calcium binding 2 | hsa-miR-1908-3p | -0,25 | -0,25 |
| KLC2 | kinesin light chain 2 | hsa-miR-1908-3p | -0,25 | -0,25 |
| VASH2 | vasohibin 2 | hsa-miR-1908-3p | -0,25 | -0,26 |
| MBP | myelin basic protein | hsa-miR-1908-3p | -0,25 | -0,3 |
| CDC34 | cell division cycle 34 | hsa-miR-1908-3p | -0,25 | -0,3 |
| NTN1 | netrin 1 | hsa-miR-1908-3p | -0,25 | -0,25 |
| SCRT1 | scratch homolog 1, zinc finger protein (Drosophila) | hsa-miR-1908-3p | -0,25 | -0,25 |
| NFYC | nuclear transcription factor Y, gamma | hsa-miR-1908-3p | -0,25 | -0,25 |
| MMP25 | matrix metallopeptidase 25 | hsa-miR-1908-3p | -0,25 | -0,25 |
| NUDT16L1 | nudix (nucleoside diphosphate linked moiety X)-type motif 16-like 1 | hsa-miR-1908-3p | -0,25 | -0,25 |
| ATP8B3 | ATPase, aminophospholipid transporter, class I, type 8B, member 3 | hsa-miR-1908-3p | -0,24 | -0,24 |
| KCTD15 | potassium channel tetramerization domain containing 15 | hsa-miR-1908-3p | -0,24 | -0,25 |
| ISM2 | isthmin 2 | hsa-miR-1908-3p | -0,24 | -0,24 |
| TMEM184A | transmembrane protein 184A | hsa-miR-1908-3p | -0,24 | -0,37 |
| HMX2 | H6 family homeobox 2 | hsa-miR-1908-3p | -0,24 | -0,73 |
| SFT2D3 | SFT2 domain containing 3 | hsa-miR-1908-3p | -0,24 | -0,24 |
| ATHL1 | ATH1, acid trehalase-like 1 (yeast) | hsa-miR-1908-3p | -0,24 | -0,93 |
| B3GNT7 | UDP-GlcNAc:betaGal beta-1,3-N-acetylglucosaminyltransferase 7 | hsa-miR-1908-3p | -0,24 | -0,24 |
| REXO1L1 | REX1, RNA exonuclease 1 homolog (S. cerevisiae)-like 1 | hsa-miR-1908-3p | -0,24 | -0,24 |
| BBS1 | Bardet-Biedl syndrome 1 | hsa-miR-1908-3p | -0,23 | -0,23 |
| TAF1C | TATA box binding protein (TBP)-associated factor, RNA polymerase I, C, 110kDa | hsa-miR-1908-3p | -0,23 | -0,32 |
| MFRP | membrane frizzled-related protein | hsa-miR-1908-3p | -0,23 | -0,23 |
| APC2 | adenomatosis polyposis coli 2 | hsa-miR-1908-3p | -0,23 | -0,23 |
| HOXA10 | homeobox A10 | hsa-miR-1908-3p | -0,23 | -0,23 |
| SOX12 | SRY (sex determining region Y)-box 12 | hsa-miR-1908-3p | -0,23 | -0,23 |
| CHRNA4 | cholinergic receptor, nicotinic, alpha 4 (neuronal) | hsa-miR-1908-3p | -0,23 | -0,46 |
| TANGO2 | transport and golgi organization 2 homolog (Drosophila) | hsa-miR-1908-3p | -0,23 | -0,24 |
| ATP11A | ATPase, class VI, type 11A | hsa-miR-1908-3p | -0,23 | -0,28 |
| HOXD9 | homeobox D9 | hsa-miR-1908-3p | -0,23 | -0,23 |
| MAPT | microtubule-associated protein tau | hsa-miR-1908-3p | -0,23 | -0,23 |
| MUC5B | mucin 5B, oligomeric mucus/gel-forming | hsa-miR-1908-3p | -0,23 | -0,23 |
| MAML3 | mastermind-like 3 (Drosophila) | hsa-miR-1908-3p | -0,23 | -0,23 |
| CCM2 | cerebral cavernous malformation 2 | hsa-miR-1908-3p | -0,23 | -0,23 |
| NDUFS7 | NADH dehydrogenase (ubiquinone) Fe-S protein 7, 20kDa (NADH-coenzyme Q reductase) | hsa-miR-1908-3p | -0,23 | -0,23 |
| PITPNM2 | phosphatidylinositol transfer protein, membrane-associated 2 | hsa-miR-1908-3p | -0,23 | -0,23 |
| MRGBP | MRG/MORF4L binding protein | hsa-miR-1908-3p | -0,22 | -0,31 |
| EXTL1 | exostosin-like glycosyltransferase 1 | hsa-miR-1908-3p | -0,22 | -0,33 |
| STON1 | stonin 1 | hsa-miR-1908-3p | -0,22 | -0,23 |
| SIX6 | SIX homeobox 6 | hsa-miR-1908-3p | -0,22 | -0,22 |
| CCDC42B | coiled-coil domain containing 42B | hsa-miR-1908-3p | -0,22 | -0,22 |
| ASB16 | ankyrin repeat and SOCS box containing 16 | hsa-miR-1908-3p | -0,22 | -0,34 |
| SLC44A2 | solute carrier family 44 (choline transporter), member 2 | hsa-miR-1908-3p | -0,21 | -0,21 |
| CATSPERG | catsper channel auxiliary subunit gamma | hsa-miR-1908-3p | -0,21 | -0,21 |
| NUP210 | nucleoporin 210kDa | hsa-miR-1908-3p | -0,21 | -0,21 |
| ITPRIPL2 | inositol 1,4,5-trisphosphate receptor interacting protein-like 2 | hsa-miR-1908-3p | -0,21 | -0,23 |
| ARHGAP26 | Rho GTPase activating protein 26 | hsa-miR-1908-3p | -0,21 | -0,39 |
| NFIC | nuclear factor I/C (CCAAT-binding transcription factor) | hsa-miR-1908-3p | -0,2 | -0,2 |
| NDUFA11 | NADH dehydrogenase (ubiquinone) 1 alpha subcomplex, 11, 14.7kDa | hsa-miR-1908-3p | -0,2 | -0,2 |
| ZNF460 | zinc finger protein 460 | hsa-miR-1908-3p | -0,2 | -0,21 |
| LRRC14 | leucine rich repeat containing 14 | hsa-miR-1908-3p | -0,2 | -0,27 |
| C1QTNF7 | C1q and tumor necrosis factor related protein 7 | hsa-miR-1908-3p | -0,2 | -0,2 |
| DACT2 | dishevelled-binding antagonist of beta-catenin 2 | hsa-miR-1908-3p | -0,2 | -0,2 |
| IRAK2 | interleukin-1 receptor-associated kinase 2 | hsa-miR-1908-3p | -0,19 | -0,19 |
| SETD1B | SET domain containing 1B | hsa-miR-1908-3p | -0,19 | -0,19 |
| FAM213B | family with sequence similarity 213, member B | hsa-miR-1908-3p | -0,19 | -0,2 |
| CAMK2B | calcium/calmodulin-dependent protein kinase II beta | hsa-miR-1908-3p | -0,18 | -0,18 |
| PRDM16 | PR domain containing 16 | hsa-miR-1908-3p | -0,18 | -0,2 |
| FAM83F | family with sequence similarity 83, member F | hsa-miR-1908-3p | -0,18 | -0,18 |
| DAB2IP | DAB2 interacting protein | hsa-miR-1908-3p | -0,18 | -0,18 |
| ALG1 | ALG1, chitobiosyldiphosphodolichol beta-mannosyltransferase | hsa-miR-1908-3p | -0,18 | -0,35 |
| ATP2A3 | ATPase, Ca++ transporting, ubiquitous | hsa-miR-1908-3p | -0,18 | -0,18 |
| LINGO1 | leucine rich repeat and Ig domain containing 1 | hsa-miR-1908-3p | -0,17 | -0,17 |
| RNF44 | ring finger protein 44 | hsa-miR-1908-3p | -0,17 | -0,17 |
| PHF15 | PHD finger protein 15 | hsa-miR-1908-3p | -0,17 | -0,25 |
| ALX4 | ALX homeobox 4 | hsa-miR-1908-3p | -0,17 | -0,17 |
| PPP1R3G | protein phosphatase 1, regulatory subunit 3G | hsa-miR-1908-3p | -0,17 | -0,24 |
| PDPK1 | 3-phosphoinositide dependent protein kinase-1 | hsa-miR-1908-3p | -0,17 | -0,17 |
| SPECC1L | sperm antigen with calponin homology and coiled-coil domains 1-like | hsa-miR-1908-3p | -0,16 | -0,16 |
| HES4 | hairy and enhancer of split 4 (Drosophila) | hsa-miR-1908-3p | -0,16 | -0,22 |
| RBM33 | RNA binding motif protein 33 | hsa-miR-1908-3p | -0,16 | -0,39 |
| SZT2 | seizure threshold 2 homolog (mouse) | hsa-miR-1908-3p | -0,16 | -0,26 |
| DOT1L | DOT1-like histone H3K79 methyltransferase | hsa-miR-1908-3p | -0,16 | -0,19 |
| VAMP1 | vesicle-associated membrane protein 1 (synaptobrevin 1) | hsa-miR-1908-3p | -0,16 | -0,16 |
| EGR3 | early growth response 3 | hsa-miR-1908-3p | -0,16 | -0,16 |
| SLC22A11 | solute carrier family 22 (organic anion/urate transporter), member 11 | hsa-miR-1908-3p | -0,16 | -0,24 |
| FANCC | Fanconi anemia, complementation group C | hsa-miR-1908-3p | -0,16 | -0,22 |
| FAM78A | family with sequence similarity 78, member A | hsa-miR-1908-3p | -0,16 | -0,24 |
| C5orf45 | chromosome 5 open reading frame 45 | hsa-miR-1908-3p | -0,15 | -0,15 |
| OTUD7A | OTU domain containing 7A | hsa-miR-1908-3p | -0,15 | -0,25 |
| KIAA1614 | KIAA1614 | hsa-miR-1908-3p | -0,15 | -0,15 |
| ZNF646 | zinc finger protein 646 | hsa-miR-1908-3p | -0,15 | -0,15 |
| SLC7A5 | solute carrier family 7 (amino acid transporter light chain, L system), member 5 | hsa-miR-1908-3p | -0,15 | -0,27 |
| HMX3 | H6 family homeobox 3 | hsa-miR-1908-3p | -0,15 | -0,36 |
| KIF13B | kinesin family member 13B | hsa-miR-1908-3p | -0,14 | -0,14 |
| ARHGAP27 | Rho GTPase activating protein 27 | hsa-miR-1908-3p | -0,14 | -0,14 |
| POU3F3 | POU class 3 homeobox 3 | hsa-miR-1908-3p | -0,14 | -0,14 |
| SLC35F1 | solute carrier family 35, member F1 | hsa-miR-1908-3p | -0,14 | -0,14 |
| MLXIP | MLX interacting protein | hsa-miR-1908-3p | -0,14 | -0,14 |
| PLD5 | phospholipase D family, member 5 | hsa-miR-1908-3p | -0,13 | -0,13 |
| SLC17A5 | solute carrier family 17 (acidic sugar transporter), member 5 | hsa-miR-1908-3p | -0,13 | -0,29 |
| UTP11L | UTP11-like, U3 small nucleolar ribonucleoprotein, (yeast) | hsa-miR-1908-3p | -0,13 | -0,23 |
| HOXD11 | homeobox D11 | hsa-miR-1908-3p | -0,13 | -0,42 |
| ISL2 | ISL LIM homeobox 2 | hsa-miR-1908-3p | -0,13 | -0,97 |
| SLC9A3R2 | solute carrier family 9, subfamily A (NHE3, cation proton antiporter 3), member 3 regulator 2 | hsa-miR-1908-3p | -0,13 | -0,43 |
| ATP8A2 | ATPase, aminophospholipid transporter, class I, type 8A, member 2 | hsa-miR-1908-3p | -0,12 | -0,26 |
| ZNF665 | zinc finger protein 665 | hsa-miR-1908-3p | -0,12 | -0,14 |
| EPB41 | erythrocyte membrane protein band 4.1 (elliptocytosis 1, RH-linked) | hsa-miR-1908-3p | -0,12 | -0,12 |
| EXOSC2 | exosome component 2 | hsa-miR-1908-3p | -0,12 | -0,37 |
| HIST1H2BO | histone cluster 1, H2bo | hsa-miR-1908-3p | -0,12 | -0,39 |
| ZNF512B | zinc finger protein 512B | hsa-miR-1908-3p | -0,12 | -0,12 |
| GLIPR2 | GLI pathogenesis-related 2 | hsa-miR-1908-3p | -0,11 | -0,44 |
| SLC33A1 | solute carrier family 33 (acetyl-CoA transporter), member 1 | hsa-miR-1908-3p | -0,11 | -0,33 |
| DSEL | dermatan sulfate epimerase-like | hsa-miR-1908-3p | -0,11 | -0,15 |
| TRPM3 | transient receptor potential cation channel, subfamily M, member 3 | hsa-miR-1908-3p | -0,11 | -0,25 |
| KIF21B | kinesin family member 21B | hsa-miR-1908-3p | -0,1 | -0,1 |
| GPM6B | glycoprotein M6B | hsa-miR-1908-3p | -0,1 | -0,26 |
| N4BP1 | NEDD4 binding protein 1 | hsa-miR-1908-3p | -0,09 | -0,13 |
| NSMF | NMDA receptor synaptonuclear signaling and neuronal migration factor | hsa-miR-1908-3p | -0,09 | -0,39 |
| DLGAP2 | discs, large (Drosophila) homolog-associated protein 2 | hsa-miR-1908-3p | -0,09 | -0,09 |
| TRIM67 | tripartite motif containing 67 | hsa-miR-1908-3p | -0,09 | -0,16 |
| CBLN1 | cerebellin 1 precursor | hsa-miR-1908-3p | -0,08 | -0,22 |
| CTD-2368P22.1 | HCG1811579; Uncharacterized protein | hsa-miR-1908-3p | -0,08 | -0,08 |
| PURB | purine-rich element binding protein B | hsa-miR-1908-3p | -0,08 | -0,19 |
| NDOR1 | NADPH dependent diflavin oxidoreductase 1 | hsa-miR-1908-3p | -0,08 | -0,09 |
| FOXK2 | forkhead box K2 | hsa-miR-1908-3p | -0,08 | -0,5 |
| USP40 | ubiquitin specific peptidase 40 | hsa-miR-1908-3p | -0,08 | -0,11 |
| MFN1 | mitofusin 1 | hsa-miR-1908-3p | -0,08 | -0,12 |
| ZDHHC3 | zinc finger, DHHC-type containing 3 | hsa-miR-1908-3p | -0,08 | -0,26 |
| DUSP2 | dual specificity phosphatase 2 | hsa-miR-1908-3p | -0,08 | -0,22 |
| TRAPPC2L | trafficking protein particle complex 2-like | hsa-miR-1908-3p | -0,07 | -0,53 |
| SKI | v-ski avian sarcoma viral oncogene homolog | hsa-miR-1908-3p | -0,07 | -0,07 |
| TTC36 | tetratricopeptide repeat domain 36 | hsa-miR-1908-3p | -0,07 | -0,41 |
| LMNB2 | lamin B2 | hsa-miR-1908-3p | -0,07 | -0,58 |
| SMURF1 | SMAD specific E3 ubiquitin protein ligase 1 | hsa-miR-1908-3p | -0,06 | -0,27 |
| DNM1 | dynamin 1 | hsa-miR-1908-3p | -0,06 | -0,09 |
| PCDHB16 | protocadherin beta 16 | hsa-miR-1908-3p | -0,06 | -0,41 |
| SCAND3 | SCAN domain containing 3 | hsa-miR-1908-3p | -0,06 | -0,12 |
| MMP17 | matrix metallopeptidase 17 (membrane-inserted) | hsa-miR-1908-3p | -0,06 | -0,14 |
| TIMM22 | translocase of inner mitochondrial membrane 22 homolog (yeast) | hsa-miR-1908-3p | -0,06 | -0,19 |
| SOX11 | SRY (sex determining region Y)-box 11 | hsa-miR-1908-3p | -0,05 | -0,27 |
| CYTH2 | cytohesin 2 | hsa-miR-1908-3p | -0,05 | -0,39 |
| TFAP2B | transcription factor AP-2 beta (activating enhancer binding protein 2 beta) | hsa-miR-1908-3p | -0,05 | -0,25 |
| PRSS21 | protease, serine, 21 (testisin) | hsa-miR-1908-3p | -0,05 | -0,24 |
| CLYBL | citrate lyase beta like | hsa-miR-1908-3p | -0,05 | -0,94 |
| TMEM127 | transmembrane protein 127 | hsa-miR-1908-3p | -0,05 | -0,22 |
| APLP2 | amyloid beta (A4) precursor-like protein 2 | hsa-miR-1908-3p | -0,04 | -0,34 |
| TCF21 | transcription factor 21 | hsa-miR-1908-3p | -0,04 | -0,45 |
| MAPK1IP1L | mitogen-activated protein kinase 1 interacting protein 1-like | hsa-miR-1908-3p | -0,04 | -0,2 |
| IRGQ | immunity-related GTPase family, Q | hsa-miR-1908-3p | -0,04 | -0,23 |
| NIPA1 | non imprinted in Prader-Willi/Angelman syndrome 1 | hsa-miR-1908-3p | -0,04 | -0,22 |
| GPR156 | G protein-coupled receptor 156 | hsa-miR-1908-3p | -0,04 | -0,2 |
| HARBI1 | harbinger transposase derived 1 | hsa-miR-1908-3p | -0,04 | -0,34 |
| MYCBP | MYC binding protein | hsa-miR-1908-3p | -0,03 | -0,35 |
| VGLL4 | vestigial like 4 (Drosophila) | hsa-miR-1908-3p | -0,03 | -0,27 |
| PDGFA | platelet-derived growth factor alpha polypeptide | hsa-miR-1908-3p | -0,03 | -0,27 |
| ARHGAP35 | Rho GTPase activating protein 35 | hsa-miR-1908-3p | -0,03 | -0,03 |
| DBP | D site of albumin promoter (albumin D-box) binding protein | hsa-miR-1908-3p | -0,03 | -0,25 |
| TXNDC17 | thioredoxin domain containing 17 | hsa-miR-1908-3p | -0,03 | -0,5 |
| CFD | complement factor D (adipsin) | hsa-miR-1908-3p | -0,03 | -0,7 |
| GJA9 | gap junction protein, alpha 9, 59kDa | hsa-miR-1908-3p | -0,03 | -0,28 |
| NPR1 | natriuretic peptide receptor A/guanylate cyclase A (atrionatriuretic peptide receptor A) | hsa-miR-1908-3p | -0,03 | -0,13 |
| TMBIM4 | transmembrane BAX inhibitor motif containing 4 | hsa-miR-1908-3p | -0,03 | -1,49 |
| PGPEP1 | pyroglutamyl-peptidase I | hsa-miR-1908-3p | -0,02 | -0,2 |
| HOXB13 | homeobox B13 | hsa-miR-1908-3p | -0,02 | -0,45 |
| PANX1 | pannexin 1 | hsa-miR-1908-3p | -0,02 | -0,2 |
| FOXE1 | forkhead box E1 (thyroid transcription factor 2) | hsa-miR-1908-3p | -0,02 | -0,2 |
| CREBBP | CREB binding protein | hsa-miR-1908-3p | -0,02 | -0,17 |
| NFIX | nuclear factor I/X (CCAAT-binding transcription factor) | hsa-miR-1908-3p | -0,02 | -0,1 |
| VPS53 | vacuolar protein sorting 53 homolog (S. cerevisiae) | hsa-miR-1908-3p | -0,02 | -0,28 |
| HEXA | hexosaminidase A (alpha polypeptide) | hsa-miR-1908-3p | -0,02 | -0,15 |
| UNKL | unkempt homolog (Drosophila)-like | hsa-miR-1908-3p | -0,02 | -0,25 |
| RBM28 | RNA binding motif protein 28 | hsa-miR-1908-3p | -0,02 | -0,19 |
| TMEM120B | transmembrane protein 120B | hsa-miR-1908-3p | -0,02 | -0,23 |
| BRD3 | bromodomain containing 3 | hsa-miR-1908-3p | -0,02 | -0,2 |
| LPAR2 | lysophosphatidic acid receptor 2 | hsa-miR-1908-3p | -0,02 | -0,39 |
| VPS18 | vacuolar protein sorting 18 homolog (S. cerevisiae) | hsa-miR-1908-3p | -0,02 | -0,36 |
| PEX10 | peroxisomal biogenesis factor 10 | hsa-miR-1908-3p | -0,02 | -0,21 |
| LRRC27 | leucine rich repeat containing 27 | hsa-miR-1908-3p | -0,01 | -0,22 |
| SMU1 | smu-1 suppressor of mec-8 and unc-52 homolog (C. elegans) | hsa-miR-1908-3p | -0,01 | -0,26 |
| UBD | ubiquitin D | hsa-miR-1908-3p | -0,01 | -0,57 |
| SLC47A1 | solute carrier family 47 (multidrug and toxin extrusion), member 1 | hsa-miR-1908-3p | -0,01 | -0,4 |
| ABCF3 | ATP-binding cassette, sub-family F (GCN20), member 3 | hsa-miR-1908-3p | -0,01 | -0,41 |
| EIF4A3 | eukaryotic translation initiation factor 4A3 | hsa-miR-1908-3p | -0,01 | -0,35 |
| ANGEL1 | angel homolog 1 (Drosophila) | hsa-miR-1908-3p | -0,01 | -0,35 |
| GMPPB | GDP-mannose pyrophosphorylase B | hsa-miR-1908-3p | -0,01 | -0,21 |
| CENPA | centromere protein A | hsa-miR-1908-3p | -0,01 | -0,18 |
| NFYA | nuclear transcription factor Y, alpha | hsa-miR-1908-3p | -0,01 | -0,27 |
| CABLES1 | Cdk5 and Abl enzyme substrate 1 | hsa-miR-1908-3p | -0,01 | -0,21 |
| CENPM | centromere protein M | hsa-miR-1908-3p | -0,01 | -0,34 |
| SNRPD1 | small nuclear ribonucleoprotein D1 polypeptide 16kDa | hsa-miR-1908-3p | -0,01 | -0,21 |
| RP1-170O19.20 | Uncharacterized protein | hsa-miR-1908-3p | -0,01 | -0,47 |
| HOXA9 | homeobox A9 | hsa-miR-1908-3p | -0,01 | -0,47 |
| RBM15B | RNA binding motif protein 15B | hsa-miR-1908-3p | -0,01 | -0,19 |
| TEAD3 | TEA domain family member 3 | hsa-miR-1908-3p | -0,01 | -0,33 |
| RRP1 | ribosomal RNA processing 1 | hsa-miR-1908-3p | -0,01 | -0,2 |
| RPL27A | ribosomal protein L27a | hsa-miR-1908-3p | -0,01 | -0,32 |
| MTG2 | mitochondrial ribosome-associated GTPase 2 | hsa-miR-1908-3p | -0,01 | -0,34 |
| ITGB5 | integrin, beta 5 | hsa-miR-1908-3p | -0,01 | -0,42 |
| HPS1 | Hermansky-Pudlak syndrome 1 | hsa-miR-1908-3p | 0 | -0,34 |
| DBNL | drebrin-like | hsa-miR-1908-3p | 0 | -0,24 |
| PHB2 | prohibitin 2 | hsa-miR-1908-3p | 0 | -0,17 |
| MGAT1 | mannosyl (alpha-1,3-)-glycoprotein beta-1,2-N-acetylglucosaminyltransferase | hsa-miR-1908-3p | 0 | -0,27 |
| C11orf80 | chromosome 11 open reading frame 80 | hsa-miR-1908-3p | 0 | -0,67 |
| CYB5R4 | cytochrome b5 reductase 4 | hsa-miR-1908-3p | 0 | -0,28 |
| CHSY1 | chondroitin sulfate synthase 1 | hsa-miR-1908-3p | 0 | -0,16 |
| PCYT2 | phosphate cytidylyltransferase 2, ethanolamine | hsa-miR-1908-3p | 0 | -0,32 |
| PKNOX1 | PBX/knotted 1 homeobox 1 | hsa-miR-1908-3p | 0 | -0,1 |
| RNPEPL1 | arginyl aminopeptidase (aminopeptidase B)-like 1 | hsa-miR-1908-3p | 0 | -0,17 |
| TRIOBP | TRIO and F-actin binding protein | hsa-miR-1908-3p | 0 | -0,19 |
| MLH1 | mutL homolog 1 | hsa-miR-1908-3p | 0 | -0,43 |
| CCDC127 | coiled-coil domain containing 127 | hsa-miR-1908-3p | 0 | -0,27 |
| PSMF1 | proteasome (prosome, macropain) inhibitor subunit 1 (PI31) | hsa-miR-1908-3p | 0 | -0,33 |
| ZDHHC4 | zinc finger, DHHC-type containing 4 | hsa-miR-1908-3p | 0 | -0,47 |
| TAB1 | TGF-beta activated kinase 1/MAP3K7 binding protein 1 | hsa-miR-1908-3p | 0 | -0,24 |
| WDR33 | WD repeat domain 33 | hsa-miR-1908-3p | 0 | -0,2 |
| NDUFA10 | NADH dehydrogenase (ubiquinone) 1 alpha subcomplex, 10, 42kDa | hsa-miR-1908-3p | 0 | -0,26 |
| CTC1 | CTS telomere maintenance complex component 1 | hsa-miR-1908-3p | 0 | -0,17 |
| RNF126 | ring finger protein 126 | hsa-miR-1908-3p | 0 | -0,23 |
| RP11-210M15.2 | Uncharacterized protein | hsa-miR-1908-3p | 0 | -0,53 |
| JSRP1 | junctional sarcoplasmic reticulum protein 1 | hsa-miR-1908-3p | 0 | -0,62 |
| C9orf3 | chromosome 9 open reading frame 3 | hsa-miR-1908-3p | 0 | -0,36 |
| GEMIN8 | gem (nuclear organelle) associated protein 8 | hsa-miR-1908-3p | 0 | -0,35 |
| CAPZB | capping protein (actin filament) muscle Z-line, beta | hsa-miR-1908-3p | 0 | -0,12 |
| MYADM | myeloid-associated differentiation marker | hsa-miR-1908-3p | 0 | -0,43 |
| SOBP | sine oculis binding protein homolog (Drosophila) | hsa-miR-1908-3p | 0 | -0,41 |
| TRA2B | transformer 2 beta homolog (Drosophila) | hsa-miR-1908-3p | 0 | -0,15 |
| BCAT1 | branched chain amino-acid transaminase 1, cytosolic | hsa-miR-1908-3p | 0 | -0,17 |
| KIAA1551 | KIAA1551 | hsa-miR-1908-3p | 0 | -0,17 |
| PPP2R1A | protein phosphatase 2, regulatory subunit A, alpha | hsa-miR-1908-3p | 0 | -0,33 |
| BCDIN3D | BCDIN3 domain containing | hsa-miR-1908-3p | 0 | -0,31 |
| PSTPIP1 | proline-serine-threonine phosphatase interacting protein 1 | hsa-miR-1908-3p | 0 | -0,12 |
| FAM207A | family with sequence similarity 207, member A | hsa-miR-1908-3p | 0 | -0,19 |
| EHD3 | EH-domain containing 3 | hsa-miR-1908-3p | 0 | -0,09 |
| ADAMTS13 | ADAM metallopeptidase with thrombospondin type 1 motif, 13 | hsa-miR-1908-3p | 0 | 0 |
| PLEKHG5 | pleckstrin homology domain containing, family G (with RhoGef domain) member 5 | hsa-miR-1908-3p | 0 | -0,49 |
| URB1 | URB1 ribosome biogenesis 1 homolog (S. cerevisiae) | hsa-miR-1908-3p | 0 | 0 |
| BCL7A | B-cell CLL/lymphoma 7A | hsa-miR-1908-3p | 0 | 0 |
| DNAH17 | dynein, axonemal, heavy chain 17 | hsa-miR-1908-3p | 0 | -0,03 |
| SIRT5 | sirtuin 5 | hsa-miR-1908-3p | 0 | -0,26 |
